# AbNatiV: VQ-VAE-based assessment of antibody and nanobody nativeness for hit selection, humanisation, and engineering

**DOI:** 10.1101/2023.04.28.538712

**Authors:** Aubin Ramon, Montader Ali, Misha Atkinson, Alessio Saturnino, Kieran Didi, Cristina Visentin, Stefano Ricagno, Xing Xu, Matthew Greenig, Pietro Sormanni

**Author notes:** Faculty of Mathematics, University of Cambridge, Wilberforce Road, Cambridge CB3 0WA, UK.

## Abstract

Monoclonal antibodies have emerged as key therapeutics, and nanobodies are rapidly gaining momentum following the approval of the first nanobody drug in 2019. Nonetheless, the development of these biologics as therapeutics remains a challenge. Despite the availability of established in vitro directed evolution technologies that are relatively fast and cheap to deploy, the gold standard for generating therapeutic antibodies remains discovery from animal immunization or patients. Immune-system derived antibodies tend to have favourable properties in vivo, including long half-life, low reactivity with self-antigens, and low toxicity. Here, we present AbNatiV, a deep-learning tool for assessing the nativeness of antibodies and nanobodies, i.e., their likelihood of belonging to the distribution of immune-system derived human antibodies or camelid nanobodies. AbNatiV is a multi-purpose tool that accurately predicts the nativeness of Fv sequences from any source, including synthetic libraries and computational design. It provides an interpretable score that predicts the likelihood of immunogenicity, and a residue-level profile that can guide the engineering of antibodies and nanobodies indistinguishable from immune-system-derived ones. We further introduce an automated humanisation pipeline, which we applied to two nanobodies. Wet-lab experiments show that AbNatiV-humanized nanobodies retain binding and stability at par or better than their wild type, unlike nanobodies humanised relying on conventional structural and residue-frequency analysis. We make AbNatiV available as downloadable software and as a webserver.

## 2 Introduction

Antibodies are a class of biomolecules with a remarkable ability to bind to molecular targets selectively and tightly. For this reason, they find key applications in biological research (1) and medicine, where they are widely employed as both diagnostic (2) and therapeutic agents (3). Nanobodies (Nbs) are single-domain antibodies (VHH) naturally expressed in camelids (4). They have grown in popularity due to their unique structural characteristics, which include small size, good stability and solubility, long third complementarity determining region (CDR3) that can bind to poorly accessible epitopes, and affinity and specificity at par to those of full-length antibodies (5). Furthermore, their potential as therapeutics has gained increased recognition since the approval of the first nanobody drug, Caplacizumab, in 2019 (6).

Established approaches to discover new antibodies or nanobodies for a target of interest can broadly be classified as first-generation in vivo approaches, for instance relying on animal immunisation (7), and second-generation in vitro techniques, relying on laboratory library construction and screening (8,9). More recently, a third generation of approaches based on computational design has started to emerge (9). Starting from the mid 90s, in vitro methods like phage display from naïve or synthetic libraries showed promise to replace animal immunisation or other in vivo techniques to isolate novel antibodies. In vitro selection is faster and cheaper than in vivo counterparts, has fewer ethical implications, and enables a better control over antigen presentation (10,11). However, despite the added costs and complexity, an increasing number of pharmaceutical and biotech companies prefers to obtain new antibodies by immunising transgenic animals with a humanised immune system (12,13) or by isolating them directly from patients (14,15). The reason for this choice is that, compared with in vitro directed evolution, antibody selection carried out by immune systems usually yields antibodies with higher developability potential and especially better in vivo properties, including long half-life, low immunogenicity, no toxicity, and low cross-reactivity against self-antigens (16,17). Up to now, most therapeutic antibodies continue to come from animal immunization (18). This consideration thus raises the question of whether a computational design strategy will ever rise to meet the challenge of generating antibodies with such properties.

Computational antibody design is still in its infancy. Yet, important advances have been made in the design of antibodies targeting predetermined epitopes of interest (19–23), which remains extremely laborious with laboratory-based approaches, and in the prediction and design of biophysical properties that underpin developability (24). Overall, computational design promises a cheaper and faster route for the discovery and optimisation of antibodies, while in principle affording a much better control than in vivo and in vitro techniques over other key biophysical properties such as stability and solubility (9).

Notwithstanding these advances, the computational prediction of in vivo properties remains hugely problematic. These properties, which include long half-life, low immunogenicity, and no toxicity, are difficult to measure accurately and in good throughput, and their molecular determinants remain poorly understood. This hurdle broadly affects therapeutic antibody development also beyond computational design, and a multitude of in vitro assays, referred to as developability screening assays, have been proposed as proxies for binding specificity or in vivo half-life to de-risk antibody development programmes (24–26). However, these assays typically correlate poorly with each other, and have only been shown to somewhat correlate with selected in vivo properties in limited specific examples (17,24,27). While advances have been made in the computational predictions of the outcome of some of these assays (28–30), or even in the number of such assays in which a lead antibody candidate is likely to perform poorly (31,32), it is quite clear that progress is hindered by the absence of robust well-defined experimental measurements of in vivo properties. These challenges are the key reasons behind the fact that in vivo antibody discovery from immune systems largely remains the gold-standard technology for therapeutic antibody discovery.

In this work, we introduce a novel deep learning method to bypass these challenges, by enabling the computational engineering of antibody and nanobody sequences indistinguishable from those obtained from immune systems. We call our method AbNatiV, as it provides an accurate quantification of the likelihood of a given sequence belonging to the distribution of native variable domain (Fv) sequences derived from human or camelid immune systems. We define this likelihood antibody nativeness, as it reflects the similarity to native antibodies. Therefore, Fv sequences with high nativeness can be expected to have in vivo properties comparable to those of immune-system-derived antibodies. AbNatiV consists of a vector-quantized variational auto-encoder (VQ-VAE) designed to process aligned Fv sequences and trained with masked unsupervised learning on sequences from curated native immune repertoires. Four different models are trained respectively on the Fv sequences of human heavy chains (VH), kappa light chains (Vκ), lambda light chains (Vλ), and camelid heavy chain single-domains (VHH).

AbNatiV can assess separately the degree of humanness and of VHH-nativeness of a given Fv sequence. It provides both an interpretable overall nativeness score and a residue-level nativeness profile of the Fv sequence, which can guide engineering by highlighting sequence regions harbouring liabilities. Therefore, AbNatiV can be useful for computational antibody design, but also to rank Fv sequences of any origin, including from in vitro discovery. The accuracy of AbNatiV in evaluating humanness is demonstrated in several benchmarks. In particular, we show that AbNatiV outperforms alternative methods when classifying antibody therapeutics. Moreover, we find that AbNatiV learns a representation of natural antibodies that captures high-order relationships between positions, which we show to be valuable for CDR grafting. We further introduce an automated humanisation pipeline of antibodies and nanobodies that relies on AbNatiV. For nanobodies, this approach monitors concurrently the humanness and the VHH-nativeness of a sequence. Wet-lab experiments on two nanobodies binding to distinct targets show that AbNatiV-humanised nanobodies retain binding and stability at par or better than their wild type, unlike nanobodies humanised with conventional structural and residue-frequency analysis.

Taken together, our results highlight the potential of AbNatiV in advancing antibody and nanobody engineering, serving as a valuable tool for computational design and ranking of Fv sequences from diverse sources, including in vitro discovery and synthetic libraries.

## 3 Results

### 3.1 The AbNatiV model

AbNatiV is a deep learning model trained on immune-system-derived antibody sequences. It employs an architecture inspired by that of the vector-quantized variational auto-encoder (VQ-VAE), originally proposed for image processing (i.e., for tensors of rank 3) (33). The AbNatiV architecture compresses amino-acid sequences (encoded as tensors of rank 2) into a bottleneck layer, also called embedding, where each latent variable is mapped to the closest code-vector from a learnable codebook, prior to reconstruction with a decoder (**Fig. 1A**). This vector quantisation from the codebook leads to a discrete latent representation rather than a continuous one as in standard VAE. This VQ architecture was chosen because protein sequences are discrete objects and thus may favour a discrete representation, and because it was shown to circumvent issues of posterior collapse that sometimes affect standard VAEs (33). Our model contains in the encoder and decoder both patch convolutional layers and transformers (**Fig. 1B**). These are respectively more suitable to capture local interactions along the sequence (i.e., local motifs), and long-range interactions between such local motifs or individual residues, which may be mediated by tertiary contacts. High codebook usage (i.e., high perplexity) is ensured in the bottleneck by a *k*-means initialisation of the codebook and a cosine similarity search during the nearest neighbour lookup quantisation, as it is needed to prevent poor data representation and maintain a robust training (34) (see Methods and **Supplementary** Fig. 1).

**Figure 1.**
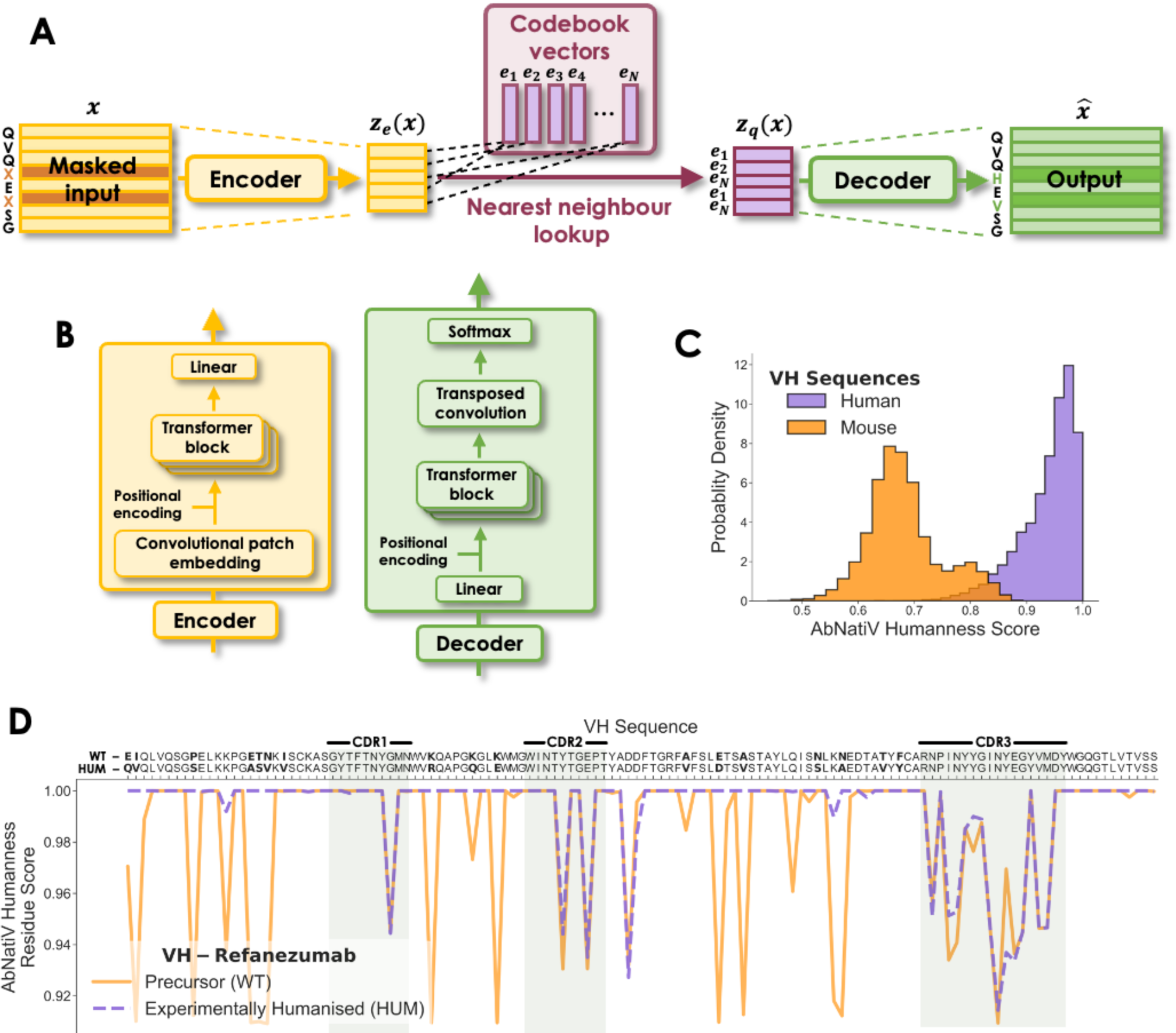
The AbNatiV model. (**A**) Architecture of the VQ-VAE-based AbNatiV model. The one-hot encoded input sequence *x* is encoded into a compressed representation *z*_*e*_(*x*) through an encoder (in yellow). In the latent space (in burgundy), *z*_*e*_(*x*) is discretised with a nearest neighbour lookup on a codebook 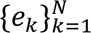 of *N* code-vectors. Each of the components of *z*_*e*_(*x*) is substituted with the closest code-vector to generate the discrete embedding *z*_*q*_(*x*). Finally, the output *x*^^^ is reconstructed through a decoder (in green) from *z*_*q*_(*x*). During training, residue masking is applied to the input *x* by replacing a portion of its residues with a masking vector (in darker shade). (**B**) Architecture of the encoder (in yellow) and decoder (in green) blocks in the AbNatiV model. (**C**) AbNatiV humanness score distributions of the VH Human (Test set, in purple) and Mouse databases (in orange). The ROC-AUC between the two distributions is 0.996. (**D**) AbNatiV humanness profiles of the VH mouse precursor and of the humanised sequence of the Refanezumab antibody therapeutic (the corresponding VL profile is in **Supplementary** Fig. 5).

The model is trained with masked unsupervised learning. Unsupervised learning works on the assumption that every sequenced antibody follows some set of biophysical and evolutionary rules that allow it to be produced by organisms and to carry out its biological function without causing toxicity. AbNatiV is built to impose a bottleneck in the network that forces a compressed representation of the input sequence, which is then reconstructed by the decoder. If the amino acids within the input sequences were fully independent from each other, this compression and subsequent reconstruction would be impossible. However, if some structure exists in the input, as it is the case of natural antibody sequences, this structure should be learnt and consequently leveraged when forcing the input through the network bottleneck. Therefore, the AbNatiV architecture is in principle capable of learning a representation of natural antibodies that captures high-order relationships between residue positions to provide a highly sensitive measure of antibody nativeness.

To ensure that the model learns meaningful high-order relationships, we also employed masked learning. During training the input sequence is masked by removing information on the identity of a random subset of residues, and the training task is to reconstruct the full sequence, including correctly predicting the identity of the masked residues (see Methods). This masking procedure is akin to a noising technique employed in denoising auto-encoders (35). From a theoretical standpoint, the approach is motivated by a manifold learning perspective, which assumes that the input data exists on a low-dimensional manifold embedded in the input space. The noising process – that is the masking/replacing of individual residues during training – shifts each training sequence away from the manifold of native antibodies, and the network is tasked with moving the data back onto the manifold via the output reconstruction of the input sequence. Additionally, the fact that the reconstruction loss also accounts for unmasked regions of the training sequences ensures that the network does not move data away from the manifold. Reconstruction accuracy is quantified with a mean square error (MSE) calculated between one-hot encoded input sequences and reconstructed output sequences. Then, at inference time, the network reconstruction of unmasked sequences represents a transformation of the input that produces an output sequence which lies closer to the manifold on which native antibodies exist. This fact establishes a crucial link between the MSE of the reconstruction and antibody nativeness, as the MSE can be interpreted as the distance of the input sequence from the manifold of native antibodies (see Methods). Reconstruction through the network always introduces some deterioration of the perfect one-hot encoded vectors, meaning that the MSE is never exactly zero, even when no residue is substituted during inference.

Taken together, AbNatiV architecture and masked unsupervised learning strategy drive the model to capture the essential features that are common across a database of native antibody sequences.

AbNatiV is trained on aligned sequences of native antibody from curated immune repertoires from the OAS database (36) and other sources (see Methods). The model is trained for 10 epochs separately on human VH, Vκ, Vλ, and camelid VHH sequences (∼2 million unique sequences in each training set). The κ and λ light chains are treated separately due to their significant differences. AbNatiV takes around 1 hour per epoch to train on a single GPU (NVIDIA RTX 8000). For each model, a validation dataset of 50,000 unique sequences different from those in the training set monitors the absence of overfitting (**Supplementary** Fig. 2) and is used for hyperparameter optimisation. 10,000 further unique sequences, distinct from those in training and validation sets, are kept aside for testing. We observe a near-perfect overlap between the distributions of the AbNatiV scores of the training and test datasets, which supports lack of overfitting (**Supplementary** Fig. 3). We further verified that there is no correlation between the AbNatiV scores of the test sequences and their median or minimum percent sequence difference to the training sequences (R^2^ ≤ 0.002, **Supplementary** Fig. 4).

For each input Fv sequence, the trained AbNatiV models return an antibody nativeness score and a sequence profile.

The nativeness score quantifies how close the input sequence is to the learnt distribution, that is to a native antibody sequence derived from the immune system the model was trained on (human or Camelid in this work). To facilitate the interpretation of this score and the comparison of scores from the different trained models, the AbNatiV score is defined in such a way that it approaches 1 for highly native sequences, and that 0.8 represents the threshold that best separates native and non-native sequences (see Methods). In the case of AbNatiV trained on VH, Vλ and Vκ human chains, this score is referred as to the AbNatiV humanness score (**Fig. 1C**). Similarly, for AbNatiV trained on VHH camelid sequences, this score is referred to as the AbNatiV VHH-nativeness score.

The sequence profile consists of one number per residue position in the aligned input sequence, so it contains a total of 149 entries including gaps. Here too, entries approaching 1 denote high nativeness, and smaller than 1 increasingly lower nativeness. This profile is useful to understand which sequence regions or residues contribute most to the overall nativeness of the sequence, and which may be liabilities. As an example, **Fig. 1D** shows the humanness profile of the VH sequence of a mouse antibody (WT precursor) that contains many low-scoring regions that could be immunogenic in humans, compared to that of its humanised counterpart: the therapeutic antibody Refanezumab. The profile of Refanezumab contains far fewer low-scoring regions, and these are mostly found in the CDR loops, which are of mouse origin and were grafted into a human Fv framework during humanisation (**Fig. 1D** and **Supplementary** Fig. 5). This example shows that sequence profiles can be powerful tools to guide antibody engineering, by facilitating the design of mutations to improve antibody nativeness.

Overall, AbNatiV predictions are highly interpretable, as nativeness scores tend to 1 with a 0.8 threshold that separates native and non-native sequences, and the sequence profile provides single-residue resolution on the sequence-determinants of nativeness.

#### 3.1.1 Classification of human antibodies

To quantify the performance of AbNatiV, we first assessed its ability to discriminate between human antibody Fv sequences and antibody Fv sequences from other species. The area under the receiver operating characteristic curve (ROC-AUC) and that under the precision-recall curve (PR-AUC) are used to quantify the ability of the models to correctly classify sequences (**Fig. 2**, **Extended Data Fig. 1**, and **Supplementary** Fig. 6). For example, AbNatiV can accurately distinguish the VH human sequences of its Test set from VH mouse sequences based on their humanness score distribution with a PR-AUC of 0.996 (**Fig. 2B**) and ROC-AUC of 0.995 (**Supplementary** Fig. 6A). Similarly, AbNatiV can successfully discriminate between human and rhesus (monkey, *Macaca mulatta*) sequences. Despite the high genetic similarity between these two organisms, the model can separate VH sequences very well, with a PR-AUC of 0.965 (**Fig. 2B**) and ROC-AUC of 0.958 (**Supplementary** Fig. 6A).

**Figure 2.**
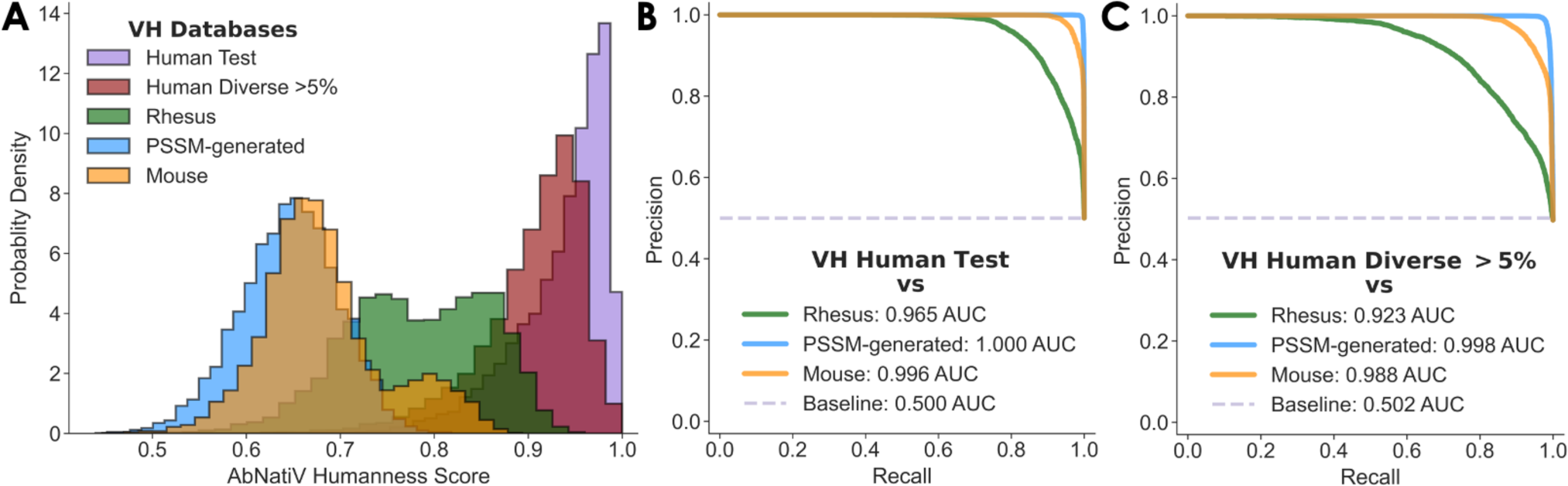
Performance on VH sequence classification. (**A**) The AbNatiV humanness score distributions of the Human Test (purple), Human Diverse >5% (red), Rhesus (green), PSSM-generated (blue), and Mouse (orange) VH antibody datasets. The PSSM-generated database is made of artificial sequences randomly generated using residue positional frequencies from the PSSM of human VH sequences. The Human Diverse >5% dataset is made of VH sequences at least 5% different from their closest sequence in the VH Training set (see Methods). (**B, C**) Plots of the PR curves of the ability of AbNatiV to distinguish the VH Human Test set (**B**) or Human Diverse >5% set (**C**) from the other datasets (see legend, which also reports the area under the curve). The baseline (dashed line) corresponds to the performance of a random classifier. The corresponding ROC curves are given in **Supplementary** Figure 6A-B.

We further employed two control datasets in our benchmark: one for the learning of high-order relationships, and one to confirm the lack of overfitting and the ability of the model to generalise to unseen sequence space. For the latter, we compiled a dataset of highly diverse human Fv sequences that we named Diverse >5% (at least 5% away from any sequence in the training set, see Methods). As expected, classification performances on the Diverse dataset slightly decrease, but overall remain very high. For the VH model, the biggest drop is found with Rhesus sequences from a PR-AUC of 0.965 with the Test set down to 0.923 with the Diverse >5% set (**Fig. 2B and C**). However, the VH model is still able to classify most of the Diverse >5% sequences as human. Only 5.5% of these sequences have a score bellow the nativeness threshold of 0.8, compared with 1.9% for the Test VH sequences. For the light chain models, the performances are even more comparable (**Extended Data Fig. 1**, and **Supplementary** Fig. 6), perhaps because the Diverse >2.5% set is less distant to the training set since diversity is more limited in light chains than in heavy chains. This performance on the control dataset is in line with our assessment of lack of overfitting (**Supplementary** Fig. 2), and it makes us confident in the ability of the model to generalise to sequences distant from those it was trained on.

As a control for the learning of high-order relationships, we generated datasets of artificial Fv sequences constructed by picking residues at random following the positional residue frequencies observed in human Fv sequences (see Methods and **Supplementary** Fig. 7). We call these datasets PSSM-generated sets. If one looks at each residue position individually, these artificial sequences are indistinguishable from real human sequences, as they are constructed only using residues observed in human sequences at each position (with log-likelihood > 0 and following the observed residue frequency distribution, see Methods). However, as residues at each position of the artificial sequences have been chosen independently of residues at other positions, any high-order relationship observed in these sequences should be compatible with random expectation. Remarkably, we find that AbNatiV can perfectly separate real VH Human sequences from PSSM-generated ones (PR-AUC of 1.000 and 0.998 respectively for VH Human Test and Diverse>5%; **Fig. 2**), and that the separation is also excellent for Vκ (PR-AUC of respectively 0.992 and 0.988; **Supplementary** Fig. 6A-C) and Vλ (PR-AUC of respectively of 0.990 and 0.980; **Supplementary** Fig. 6D-F). This performance attests the ability of AbNatiV to learn complex high-order relationships observed within native human Fv sequences beyond their simple amino acid composition.

We then compared the performances of AbNatiV with that of other computational methods developed for the humanisation of antibody sequences (**Table 1, Extended Data Tables 1-2, Supplementary Tables 1-3**, and **Supplementary** Fig. 8-9). More specifically, we focus on the recently introduced OASis 9-mer peptide similarity score (37), the Sapiens transformer model (37), and the AbLSTM model (38), as these approaches were shown to outperform older methods. Our results show that AbNatiV outperforms all alternative approaches on all classification tasks overall (**Table 1**, and **Supplementary Table 1**). The biggest difference is observed in the Human Test vs. Rhesus classification, where for VH sequences the AbNatiV PR-AUC is 0.965 while that of the best alternative method, AbLSTM, is 0.721, which increases to 0.777 once the AbLSTM architecture is re-trained on our training set (**Table 1**). Lower performances of the alternative models are also shown for the Human Test vs. Mouse and vs. PSSM-generated classification tasks. We have not included in this benchmark the recently introduced Hu-mAb method (39), since we could only access it as a webserver that processes a single sequence per run. However, as Hu-mAb is trained with supervised learning for the specific task of distinguishing between human and mouse sequences, we would expect it to do extremely well at the mouse vs. human classification task, and perhaps not as well on other tasks.

**Table 1.** Evaluation of the PR classification and reconstruction tasks for human VH sequences. The assessment is carried out for AbNatiV trained on human VH sequences (first row) and other computational approaches that can assess humanness (other rows). AbLSTM retrained corresponds to the AbLSTM model retrained on the same training set of AbNatiV (see Methods). The first six columns report the area under the PR curve (shown in **Fig. 2** and **Supplementary** Fig. 8), assessing the ability of the models to separate sequences in the Human Test (T) or the Human Diverse >5% (D) sets from those from mouse, rhesus, and PSSM-generated (see column headers). The Human Diverse >5% dataset is used here as a control to specifically assess the ability of the AbNatiV to generalise to sequences distant from those in its training set. The last two columns quantify the ability of each model to reconstruct human sequences in each dataset (column header). The OASis method does not carry out reconstruction. Many sequences of the D datasets belong to the Sapiens training set. Corresponding ROC results are in **Supplementary** Table 1.

| VH | Classification (PR-AUC) |  |  |  |  |  | Reconstruction accuracy |  |
| --- | --- | --- | --- | --- | --- | --- | --- | --- |
|  | Rhesus vs |  | Mouse vs |  | PSSM-generated vs |  | T | D |
|  | T | D | T | D | T | D |  |  |
| AbNatiV | 0.965 | 0.923 | 0.996 | 0.988 | 1.000 | 0.998 | 0.960 | 0.935 |
| OASis (relaxed) | 0.570 | 0.829 | 0.897 | 0.965 | 0.982 | 0.992 | N/A | N/A |
| Sapiens | 0.626 | 0.883 | 0.982 | 0.994 | 0.993 | 0.997 | 0.918 | 0.949 |
| AbLSTM | 0.721 | 0.892 | 0.963 | 0.986 | 0.998 | 0.998 | 0.807 | 0.856 |
| AbLSTM Retrained | 0.777 | 0.866 | 0.967 | 0.979 | 0.997 | 0.996 | 0.822 | 0.849 |

**Table 2.**
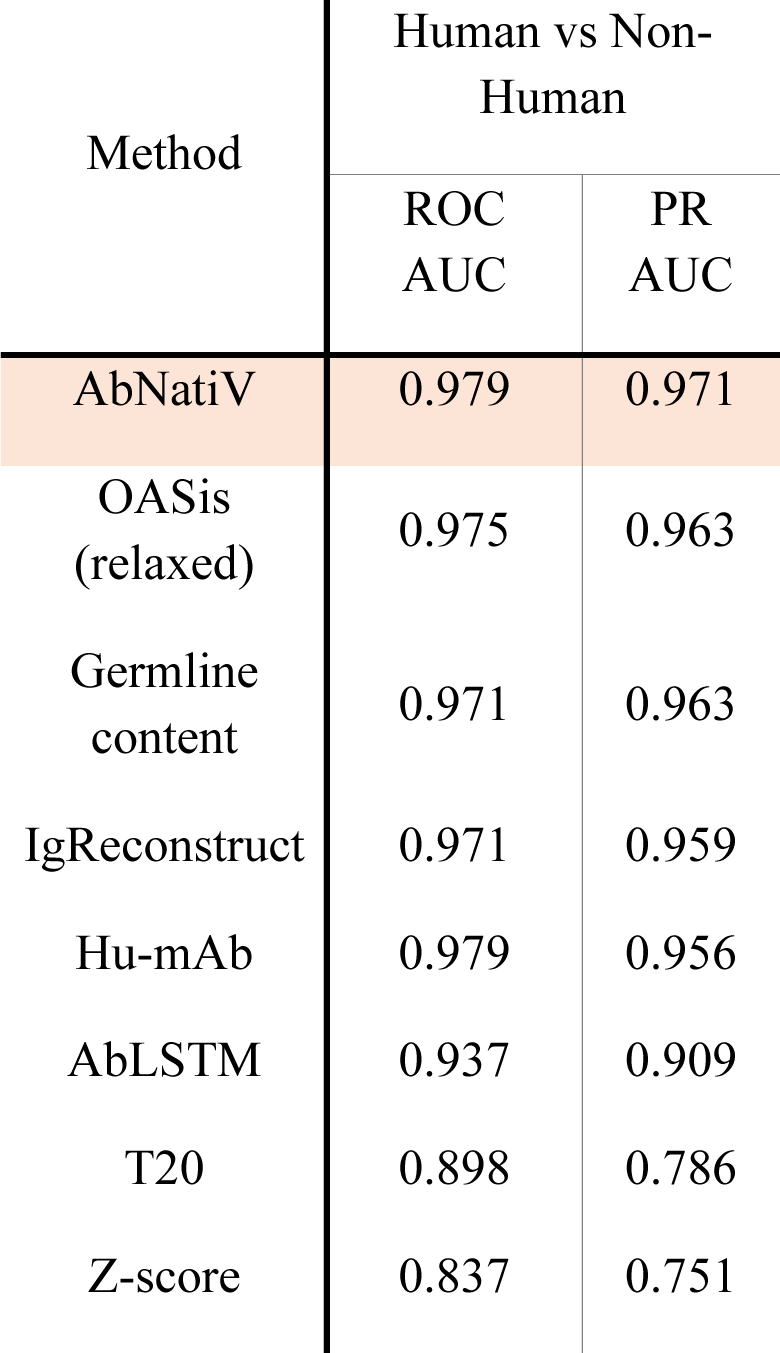
Performance on the classification of antibody therapeutics. The assessment is carried out for AbNatiV (first row) by averaging the AbNatiV humanness scores of the heavy and light chains from the relevant AbNatiV model (i.e., trained either on VH, Vκ, or Vλ, see Methods), and for other computational methods (table rows). The classification task consists in distinguishing 196 human-derived therapeutic antibodies from 353 therapeutic antibodies from a different origin (mouse, chimeric, and humanised). The area under the curve for both ROC and PR curves are reported in the first two columns.

We further carried out the same benchmarks by replacing the human Test set with the human Diverse >5% dataset, which contains sequences that are at least 5% different from any sequence in our training set. AbNatiV remains the best performing model overall. However, Sapiens marginally outperforms AbNatiV in one task: the classification of Mouse sequences (by 0.006 in PR-AUC, **Table 1**). This result is hardly surprising, as the Human Diverse >5% databases were built using sequences from the training set of Sapiens and OASis (37), and hence are overclassified with respect to our Human Test set. In addition, amino acid reconstruction accuracies were computed for all methods (except OASis as the method is not reconstruction-based). The reconstruction accuracy quantifies the ability of a model to reconstruct the initial input from the embedding in the latent space. Both AbNatiV and Sapiens rely on masked learning, while AbLSTM relies on standard unsupervised learning. We find that the former models have higher reconstruction accuracies than the AbLSTM model (96%, 92% and 81% on the Human Test set respectively for AbNatiV, Sapiens and AbLSTM). Sapiens reconstructs slightly better than AbNatiV the VH sequences in the Human Diverse dataset (respectively 94% and 95%). However, it should be noted again that the Human Diverse >5% dataset is contained in the training set of Sapiens (37).

Similar results are found for Vκ and Vλ lights chains, when comparing AbNatiV with the OASis and Sapiens methods (**Extended Data Tables 1-2**, and **Supplementary Tables 2-3**), while the AbLSTM humanness score is not defined for light chains (38). Curiously, in contrast with VH sequences, AbNatiV exhibits higher reconstruction accuracy than Sapiens also for the VL sequences in the Human Diverse >2.5% datasets (respectively 98% vs 94% for Vκ, and 98% vs 93% for Vλ).

Taken together, these results demonstrate that AbNatiV is a precise humanness assessment method that has learned high-order relationship between residues to identify antibody sequences derived from human immune systems.

#### 3.1.2 Application to antibody therapeutics

The assessment of humanness is a critical step of antibody drug development, with the goal of ensuring that drug candidates have minimal risk for administration to patients. Therefore, we ran AbNatiV on therapeutic antibody sequences and averaged the humanness score of the heavy and light chains from the relevant AbNatiV model (i.e., trained either on VH, Vκ, or Vλ, see Methods). More specifically, we evaluated the performance of the method on distinguishing 196 human therapeutics from 353 antibodies therapeutics of non-human origin (mouse, chimeric, and humanised). The precision-recall (PR) curve (**Fig. 3A**) and ROC curve (**Supplementary** Fig. 10) are computed for AbNatiV and 7 other computational approaches (see Methods and **Table 2**). AbNatiV outperforms all other methods when considering both AUCs with a PR-AUC of 0.971 and a ROC-AUC of 0.979. The second-best methods after AbNatiV are OASis with a PR-AUC of 0.963 and a ROC-AUC of 0.975 and Hu-mAb with a ROC-AUC of 0.979 and a PR-AUC of 0.956.

**Figure 3.**
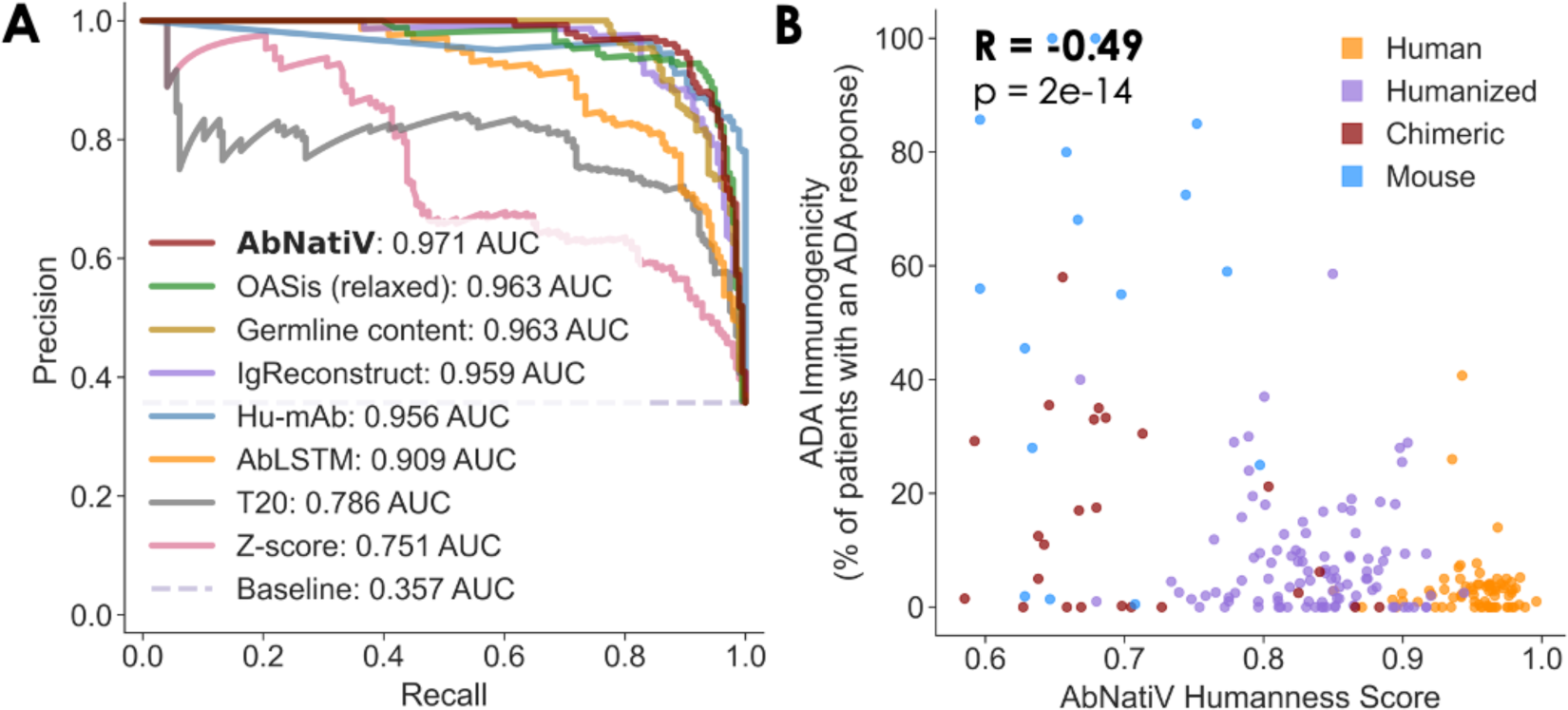
Performance on antibody therapeutics. (**A**) Plot of the PR curves of the classification of 196 human-derived therapeutics from 353 therapeutics of non-human origin (mouse, chimeric, and humanised) carried out with AbNatiV (in red) and seven other computational methods (see legend, which also reports the AUC values). The baseline (dashed line) corresponds to the performance expected from a random classifier. Corresponding ROC curves can be found in **Supplementary** Figure 10. (**B**) Scatter plot of the AbNatiV humanness score of 126 antibody therapeutics and their ADA immunogenicity score, expressed as the percentage of patients developing an ADA response in each study. The Pearson correlation (R) and p-value are reported on top left corner. Sequences are coloured based on their origin (i.e., human in orange, humanised in purple, chimeric in red, and mouse in blue).

Α central interest in humanisation of antibodies is to reduce their immunogenicity in human immune systems. One way to assess immunogenicity in early-stage clinical trials is to assess the number of patients who develop anti-drug antibodies (ADAs) in response to the administration of therapeutic antibodies (40). We find that the AbNatiV humanness score (i.e., the average of the AbNatiV humanness scores of the VH and VL, see Methods) shows a Pearson correlation coefficient (R) of - 0.49 (p-value ∼ 2x10^-14^) with the percent of patients that developed ADAs upon treatment, which is available for 216 different therapeutic antibodies (**Fig. 3B**). We note that these ADA data are highly heterogeneous and therefore there is no reason to expect much stronger correlations. The percent of patients who developed an ADA response is determined in different studies carried out in drastically different ways. In particular, the dosage of the therapeutic antibody candidate and the length of the study (i.e., the number of doses administered and the total study time) can vary widely among different therapeutic candidates. It is therefore foreseeable that a highly immunogenic antibody which is administered only once and at a relatively low dose would elicit a weaker ADA response than a less immunogenic antibody that is administered at high dose for an extended period. The reason for these discrepancies is that these clinical studies are designed around the specific requirements of the drug candidate under scrutiny, rather than to quantitatively compare the immunogenicity of different drug candidates.

#### 3.1.3 Classification of native camelid nanobodies

The development of single-domain antibodies has been gathering even more momentum since the approval of Caplacizumab in 2019, the first nanobody-based therapeutic (6). Nanobodies (VHHs) are naturally expressed in camelids and can exhibit advantageous stability and solubility properties combined with a small size that allows for better tissue penetration, while retaining the affinity and specificity of full-length antibodies (5). When trained on VHH sequences, AbNatiV returns a VHH-nativeness score that quantifies the resemblance of antibody sequences to native camelid single-domain antibody, and hence the ability of a VH sequence to fold independently of a VL counterpart.

We find that AbNatiV accurately discriminates VHH Test sequences from the VH sequences of human (0.983 PR-AUC), mouse (0.995), and rhesus (0.992) (**Fig. 4A-C**, and **Supplementary** Fig. 11). The PR-AUC between PSSM-generated artificial VHH sequences and real camelid VHH sequences from the Test set is 0.942. The VHH model can classify most of the Diverse >5% VHH sequences as native, with a performance at par to that observed on the Test set. 10.4% of Diverse >5% VHH sequences have a score bellow the nativeness characteristic threshold of 0.8, compared with 10.8% for the Test VH sequences. To the best of our knowledge, AbNatiV is the first approach to quantify the nativeness of nanobodies. Therefore, to compare with a different model, we retrained the AbLSTM architecture, originally developed for human VH sequences, on our nanobody training set (see Methods). We find that AbNatiV shows higher classification performance than the retrained AbLSTM model on all tasks, and especially on the classifications with the VHH Diverse >5% dataset (**Table 3, Supplementary Table 4**, and **Supplementary** Figure 12).

**Figure 4.**
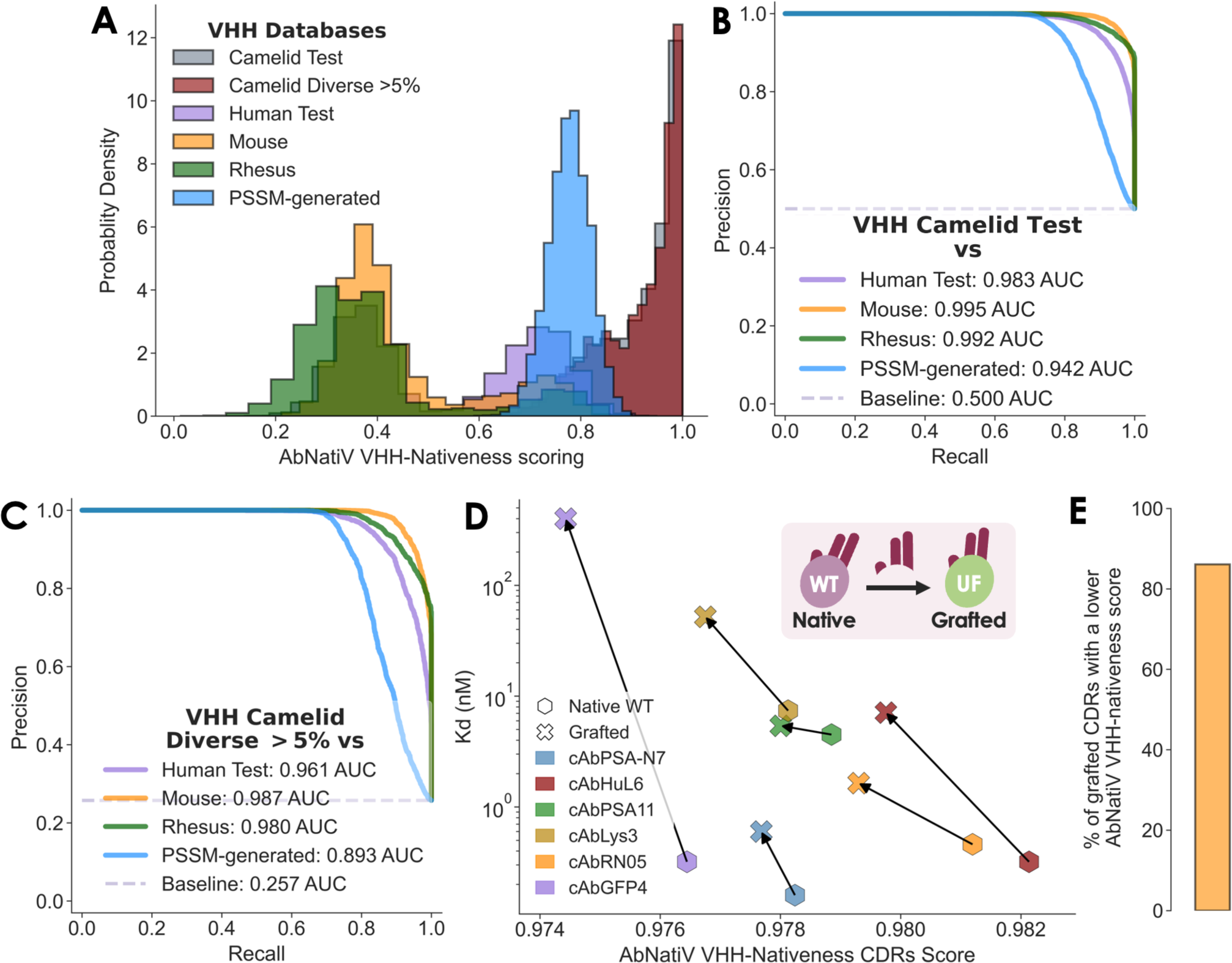
Performance on VHH sequences derived from camelids. (**A**) The AbNatiV VHH-nativeness score distributions of the VHH Camelid Test (in grey), Camelid Diverse >5% (in red), VH Human Test (in purple), VH Mouse (in orange), VH Rhesus (in green), and VHH PSSM-generated (in blue) datasets. The VHH PSSM-generated database is made of artificial sequences randomly generated using residue positional frequencies from the PSSM of VHH sequences. The Camelid Diverse >5% dataset is made of VHH sequences at least 5% different from their respective closest sequence in the VHH Training set (see Methods). Each dataset contains 10,000 sequences except Camelid Diverse >5% which contains 3,468 sequences. (**B, C**) Plot of the PR curves used to quantify the ability of AbNatiV to distinguish the VHH Camelid Test (**B**) or Camelid Diverse >5% (**C**) set from the other datasets (see legend, which also reports the AUC values). The baseline (dashed line) corresponds to the performance of a random classifier. The corresponding ROC curves are given in **Supplementary** Fig. 11. (**D**) Plot of the binding K_D_, as reported in Ref. (41), as a function of the AbNatiV VHH-nativeness score computed across all CDR positions of 6 nanobodies (see legend) before and after grafting of all three CDRs onto a camelid universal framework (UF). An arrow is directed from the native sequence in the WT framework to the grafted one. (**E**) All three CDRs from a test set of 5,000 VHH sequences are computationally grafted onto the UF (see Methods). The bar plot shows that 86% of them have a lower AbNatiV VHH-nativeness score when grafted onto the UF than when they are within their native framework.

**Table 3.** Evaluation of the PR classification and reconstruction tasks for camelid VHH sequences. The assessment is carried out for AbNatiV trained on camelid VHH sequences (first row) and the AbLSTM model retrained on the same training set of AbNatiV (see Methods and second row). The first eight columns report the area under the curve for PR curves (shown in **Fig. 4-C** and **Supplementary** Fig. 12), assessing the ability of the models to separate sequences in the Camelid Test (T) or Human Diverse >5% (D) sets from those from human, mouse, rhesus, and PSSM-generated (see column headers). The Camelid Diverse >5% dataset is used as a control to specifically assess the ability to generalise to sequences distant from those in the training set. The last two columns quantified the ability of each model to reconstruct camelid sequences in each dataset (column header). Corresponding ROC results are in **Supplementary** Table 4.

| VHH | Classification (PR-AUC) |  |  |  |  |  |  |  | Reconstruction accuracy |  |
| --- | --- | --- | --- | --- | --- | --- | --- | --- | --- | --- |
|  | Human Test vs |  | Mouse vs |  | Rhesus vs |  | PSSM-generated vs |  | Camelid Test (T) | Camelid Diverse >5% (D) |
|  | T | D | T | D | T | D | T | D |  |  |
| AbNatiV | 0.983 | 0.961 | 0.995 | 0.987 | 0.992 | 0.980 | 0.942 | 0.893 | 0.954 | 0.954 |
| AbLSTM retrained | 0.956 | 0.900 | 0.988 | 0.969 | 0.983 | 0.956 | 0.916 | 0.839 | 0.847 | 0.846 |

#### 3.1.4 CDR nativeness for grafting experiment

The grafting of target specific CDRs onto a different framework scaffold is a common technique to design antibody with enhanced properties (e.g., lower immunogenicity, higher stability or expressibility, etc.) (41–43). In the case of nanobodies, a specific camelid framework, referred to as universal framework (UF), was shown to retain very high conformational stability and prokaryotic expressibility almost independently of its CDR loops (44). In that study, all three CDRs of 6 unrelated nanobodies targeting different antigens were grafted onto the UF. Binding affinity (K_D_) and conformational stability (ΔG) were experimentally measured for all six wild type (WT) nanobodies, and corresponding UF variants with the grafted CDRs. Upon grafting, the binding K_D_ worsened for most variants, probably because the CDRs now make some non-native interactions with the UF sequence, which affects their conformation and consequently antigen binding, even if the conformational stability improved upon grafting because of the superior stability of the UF (44). AbNatiV provides a direct sequence-based approach to assess the nativeness of these CDRs within the VHH UF and their WT framework, by computing the VHH-nativeness score across all CDR positions (see Methods). We find that for all these 6 grafting examples, AbNatiV scoring anticorrelates with the experimentally measured change in binding K_D_ (**Fig. 4D**). Specifically, AbNatiV attributes a worse (lower) VHH-nativeness score to these sets of CDRs when they are grafted onto the UF than when they are found in their WT framework, in agreement with the experimental measurement of a worse (higher) binding K_D_. An example of the nativeness profile before and after grafting is provided in **Supplementary** Figure 13.

Encouraged by these findings on six experimentally characterised grafting examples, we sought to obtain more robust statistics by computationally grafting all three CDRs of 5,000 different nanobodies from the VHH Test set onto the UF scaffold. We find that in 86% of cases AbNatiV computes a lower VHH-nativeness score for the CDRs grafted in the UF than for the CDRs in their native WT framework (**Fig. 4E**). Taken together, the results of these analyses suggest that AbNatiV can accurately determine whether CDR loops are in the right context.

### 3.2 Humanisation of nanobodies

With the recent surge of interest in the use of nanobodies as therapeutics, the humanisation of nanobodies has emerged as a crucial requirement to improve their therapeutic index and reduce immunogenicity risks for clinical applications (42,45,46). **Extended Data Figure 2** depicts the AbNatiV evaluation of the humanness and VHH-nativeness of 3 nanobody therapeutics, and of 8 WT nanobodies from a SARS-CoV-2 study (47) and their humanised counterpart characterised in a separate study (45). In that study, Sang et al. introduced a computational pipeline named Llamanade (45), which integrates structural information and residue frequency statistics to humanise nanobody sequences. We find that all humanised nanobody sequences are assigned an AbNatiV humanness score higher than their WT counterpart. Importantly, this improvement of humanness impacts their VHH-nativeness only weakly or even improves it (**Extended Data Fig. 2**), which is in line with the non-significant or very small change observed experimentally by Sang et al. (45) in the binding K_D_ of these nanobodies upon humanisation.

Encouraged by these observations, we sought to develop a framework to exploit AbNatiV for the rational humanisation of nanobody sequences. By combining the humanness (VH-AbNatiV) with the VHH-nativeness (VHH-AbNatiV) assessments of AbNatiV, we propose a dual-control humanisation strategy of nanobody sequences. As illustrated in **Supplementary** Figure 14, this strategy begins by identifying liable positions with a low AbNatiV humanness or VHH-nativeness in the residue profile. Then, it suggests potentially humanising mutations derived from the human VH PSSM (**Supplementary** Fig. 7A). Finally, it accepts mutations that improve the AbNatiV humanness score while preserving or further improving the AbNatiV VHH-nativeness score (see Methods for further details).

Two distinct strategies to sample mutational variants are proposed, which we designate as “Enhanced” and “Exhaustive” sampling. The enhanced approach iteratively explores the mutational space, aiming for rapid convergence to identify a promising mutant. In contrast, the exhaustive approach assesses all mutation combinations within the available mutational space and selects the best sequence. It is important to note that the exhaustive sampling is considerably more computationally demanding. For instance, in the case of a sequence with 10 liable positions where 4 mutations are allowed at each position, the mutational space encompasses 4^10^ mutants, exceeding 1 million combinations. On the other end, the enhanced sampling will explore on average less than 100 combinations of mutations. Therefore, to manage the computational complexity of the exhaustive approach, we restrict its mutational space by constraining the allowed mutations to residues enriched in both the human VH and VHH PSSMs. Conversely, the enhanced method’s mutational space is larger as it restricts its allowed mutations to the human VH PSSM only. To minimise the chances of affecting antigen binding, both strategies are limited to the framework regions. For each sampling strategy, we implement both a purely sequence-based approach and a structure-based approach that models the nanobody structure from the input sequence (see Methods). In the latter, buried residues that are not on the nanobody surface are excluded from the list of potential targets for mutations, as commonly done in humanisation strategies based on framework resurfacing (48,49).

To test the effectiveness of these different humanisation pipelines we generated in silico humanised variants of two nanobodies, which we then produced and characterised in vitro. These two nanobodies bind to two distinct proteins of therapeutic relevance: Nb24 targets the β_2_-microglobulin (50), and mNb6 targets the receptor-binding domain (RBD) of the Spike protein of SARS-CoV-2 (matured version of Nb6 in Ref. (51)). Nb24 was obtained from a llama immunisation campaign and exhibits moderate binding with a dissociation constant K_D_ in the mid-nanomolar range (52), while mNb6 was obtained from the screening of a synthetic library and then highly optimised via saturation mutagenesis to reach a high-picomolar-range K_D_ (53). For each WT sequence, we generated four humanised variants using the AbNatiV automated pipelines and a further control variant. Two variants were generated by each sampling method: one limited to solvent-accessible framework sites, and the other encompassing all framework sites. While the crystal structures of Nb24 and Nb6 are solved experimentally (respectively PDB ids 4kdt and 7kkk), solvent-exposed sites were identified by modelling in silico the structures of the WT sequences with Nanobuilder2 (51) to simulate a more general setting in which crystal structures may not be available.

For comparison, we also generated one additional humanised variant for each WT nanobody using the automated humanization tool Llamanade that proposes humanising mutations based on structural and residue frequency analysis (45). We refer to these as Frequency and Structure-based humanised variants. All generated sequences are presented in **Extended Data Table 3**, and the human VH and VHH AbNatiV profiles in **Supplementary** Figures 15 and 16, which also highlight the mutations from the WT. As expected, all humanised sequences have improved humanness and similar VHH-nativeness to their WT, except for the two frequency and structure-based variants that show worsened VHH-nativeness (**Fig. 5A,B**).

**Figure 5.**
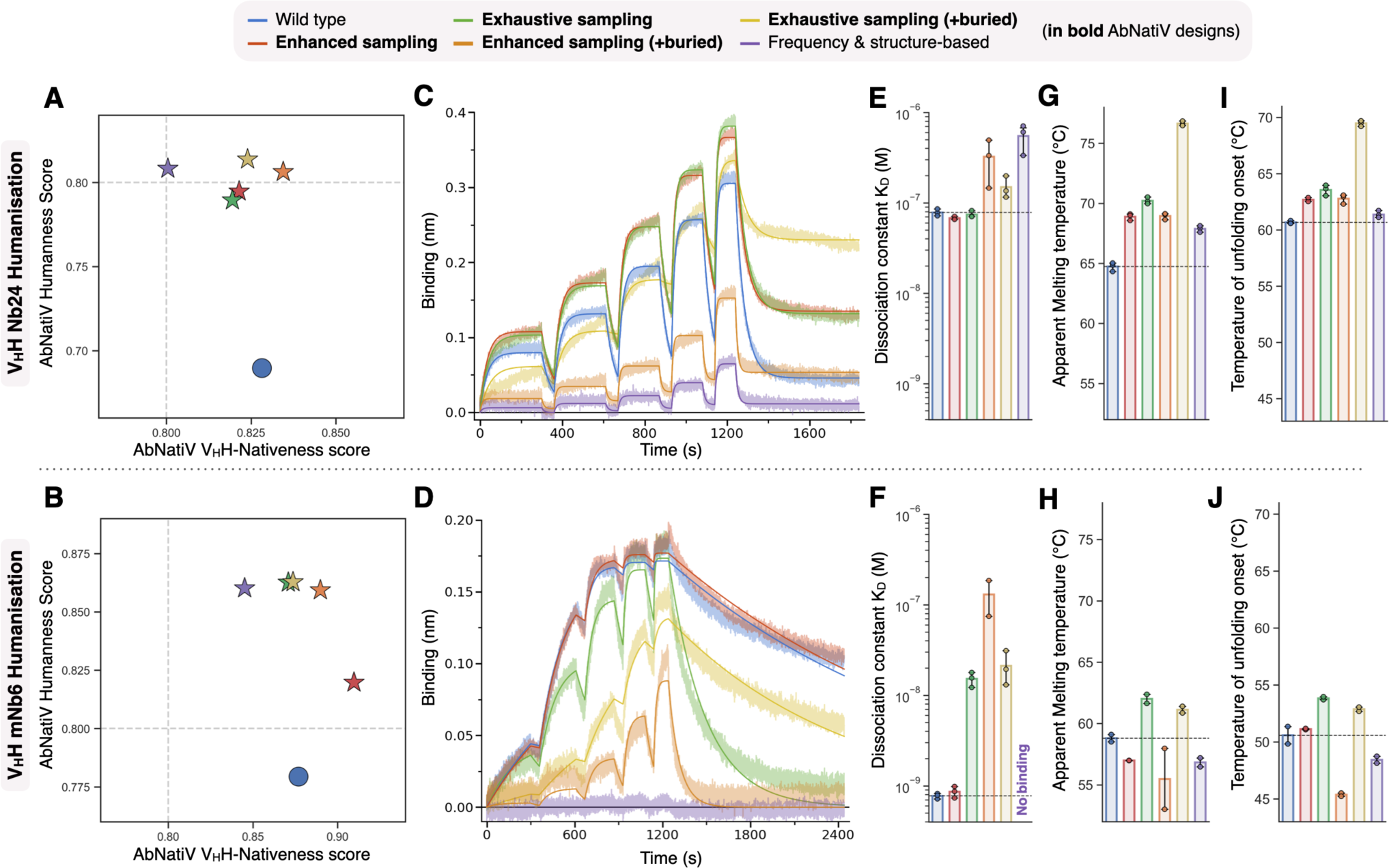
Humanisation of two llama-derived nanobodies. The top row pertains to the humanisation of nanobody Nb24, which binds human β_2_-microglobulin, the lower row to mNb6, which binds SARS-CoV-2 RBD. In the legend, variants in bold font are different AbNatiV design strategies (see text). The Frequency & structure-based designs are done with the Llamanade webserver (45). (**A, B**) Scatter plot of the AbNatiV VH humanness score as a function of the VHH-nativeness score for all characterised variants (legend, the WT is the blue circle). (**C, D**) BLI binding traces (associations and dissociations phases) obtained with SA sensors loaded with biotinylated β_2_-microglobulin (**C**) or biotinylated SARS-CoV-2 RBD (**D**). (**C**) Association was monitored in wells containing 25, 50, 100, 200, and 400 nM of Nb24 nanobody variants (see legend). Data were fitted globally with a 1:1 partial dissociation binding model (solid lines) using R_max_, on rate, and off rate as global parameters and Y_t→inf_ as local parameter. (**D**) Association was monitored in wells containing 3.7, 11.1, 33.3, 100, and 300 nM of the WT and the Enhanced sampling variants (see legend), 4, 12.2, 36.4, 109.3, and 328 nM of the Enhanced sampling (+buried) variant (orange), and 6.2, 18.5, 55.6, 166.7, and 500 nM of all other mNb6 variants (legend). Data were fitted globally with a 1:1 binding model (solid lines) using R_max_, on rate, and off rate as global parameters. Two additional independent BLI experiments per antigen, carried out with different concentrations and times, are presented in **Fig. S24**. (**E, F**) Bar plot of the fitted K_D_ values from the three experiments. (**H, G**) Bar plot of the apparent melting temperatures. (**I, J**) Bar plot of the temperatures of unfolding onset (see Methods). Error bars are standard deviations.

WT nanobodies and all humanised designs were then produced in E. Coli and experimentally characterised (see Methods).

Bio-layer interferometry (BLI) experiments show that Nb24 WT binds β_2_-microglobulin with a K_D_ of 79 ± 6 nM (mean ± standard deviation from 3 independent experiments, **Fig. 5C,E** and **Supplementary** Fig. 17), which is compatible with previously reported values (52). AbNatiV-humanised Nb24 variants obtained from both the Enhanced and the Exhaustive sampling strategies bind the antigen with K_D_ values at par or slightly better than that of the WT (respectively 68 ± 3 and 75 ± 5 nM; **Fig. 5C,E**). Conversely, humanised variants containing mutations also at buried positions showed worsened K_D_ values, and the Nb24 variant with the most compromised binding was that from the Frequency and Structure-based humanisation, with a K_D_ in the high nanomolar range (**Fig. 5C,E**).

We also measured the thermal stability of all produced nanobodies (see Methods). We find that all Nb24 humanised variants have increased apparent melting temperatures and temperatures of unfolding onset over those of the WT (**Fig. 5G**). However, this improvement is the smallest for Frequency and Structure-based humanisation, it is more pronounced for the Enhanced sampling AbNatiV humanisation, and even larger for the Exhaustive sampling strategies (**Fig. 5G,I**).

In agreement with previous reports (53), we find that WT mNb6 binds SARS-CoV-2 RBD with a K_D_ in the high picomolar range (0.78 ± 0.04 nM). The AbNatiV-humanised mNb6 variant from the Enhanced sampling strategy retains this tight K_D_ (K_D_ = 0.86 ± 0.10 nM; **Fig. 5D,F**). However, all other mNb6 humanised variants show a binding compromised to varying degrees. The least affected variant is the one from the AbNatiV Exhaustive sampling, with a K_D_ of 15 ± 2 nM, followed by the two AbNatiV variants that also contain mutations at buried sites. The most affected variant is the one from the Frequency and Structure-based humanisation, which did not yield any binding signal in the assay (**Fig. 5D** and **S24**).

In terms of thermal stability, the Enhanced sampling variants show a slight decrease of apparent melting temperature over that of the WT, but a similar or marginally improved temperature of unfolding onset. Conversely, the Enhanced Sampling variant with mutations at buried positions and the Frequency and Structure-based variant had decrease thermal stability, while both Exhaustive Sampling variants had increased thermal stability (**Fig. 5H,J**).

Taken together, these results underscore the effectiveness of the AbNatiV Enhanced Sampling humanisation pipeline to enhance in silico the humanness of nanobodies by suggesting mutations that are not detrimental to binding and stability.

## 4 Discussion

In this work we have introduced AbNatiV, a VQ-VAE-based antibody nativeness assessment method that can evaluate the likelihood of input sequences belonging to the distribution of immune-system derived antibodies (human VH and VL domains and camelid VHHs). AbNatiV provides both an interpretable overall score for the full sequence, and a nativeness profile at the residue level, which can be exploited to guide antibody engineering and humanisation. The integration of masked and unsupervised learning with the deep VQ-VAE architecture allows AbNatiV to capture complex high-order interactions. AbNatiV successfully discriminates natural sequences from artificial sequences generated following the natural positional residue frequency, and it can distinguish human antibodies or camelid nanobodies from antibodies from other species. Compared to alternative methods developed for antibody humanisation, AbNatiV exhibits higher classification performances, while often being trained on a smaller number of sequences (∼2 million) for fewer epochs (10 epochs). To put these numbers in context, the deep VH transformer model Sapiens was trained on 20 million sequences for 700 epochs (37). The training set size of the AbNatiV VHH model, comprising around 2.2 million sequences, is inherently limited by the number of VHH sequences available in the literature. Conversely, for the human heavy and light chains, 2 million sequences only were used for training despite the abundance of available data for human antibody sequences. Upon investigation, we revealed that the VH model exhibits minimal performance improvement when expanding the training set size from 1 million to 2 million sequences (see **Supplementary** Fig. 18A). This little gain of performance does not justify increasing the dataset training size further as this would substantially increase training time. Furthermore, having a training size comparable with that of the VHH model ensures a fair and meaningful performance evaluation across models.

AbNatiV is trained on aligned sequences. The alignment process is performed with the AHo antibody residue numbering scheme (54), which numbers each residue based on its structural role (e.g., being in a particular CDR loop or in the framework region). Essentially all known antibodies fit into this representation, and we posited that – albeit our method is purely sequence based – using Fv sequences aligned in this way would facilitate the learning of structural features and hence increase performance. To test this hypothesis, we employed the same architecture on non-aligned sequences (see Methods), which, as expected, led to a very notable performance drop. In the case of VH sequences, using non-aligned sequences results in a three-to-four-fold decrease of both training and validation loss performances (see **Supplementary** Fig. 18**.B**). These findings are consistent with those of Hawkins-Hooker et al. (55), who applied a fully connected VAE to a dataset of luciferase sequences. The model trained on aligned sequences captured better the information, leading to a more successful generation of new luciferase-like sequences compared to the model trained on unaligned sequences. Moreover, employing aligned sequences enables AbNatiV to produce residue profiles readily comparable across sequences of different lengths. This feature is highly advantageous for sequence engineering purposes, and for the comparison of different hits from antibody discovery or optimisation campaigns.

We have also observed that AbNatiV outperforms alternative methods when classifying human-derived antibody therapeutics from therapeutic antibodies of non-human origin, which also reflects the robustness of the AbNatiV assessment beyond the span of its training and test sets. We have further shown that AbNatiV humanness score have a statistically significant correlation (R= - 0.5) with the percent of patients that developed ADA in clinical studies. This evaluation of immunogenicity with the ADA database is commonly employed to benchmark immunogenicity assessments methods (37,39,56), and therefore we performed it in our work. However, these ADA data exhibit a substantial level of heterogeneity, as the database was assembled using immunogenicity data from different clinical studies reported in the literature, with experimental conditions (e.g., number of patients, dosage, study length) varying substantially among studies. As an example, Basiliximab was tested on 339 patients (https://www.ema.europa.eu/), while Disitamab only on 58 (57). In the study considered in the ADA dataset that we used, Disitamab is reported to elicit an ADA response in 58.6% of the patients. However, in a more recent publication on a larger study with a more uniform design (80 patients with the same dosage instead of 58 patients with 4 different dosages), Disitamab was shown to elicit ADA response in 23.8% of the participants (58), which is less than half of the number previously estimated. This example shows that the degree of heterogeneity of this ADA database should be considered when expecting quantitative correlations with immunogenicity predictions. Nevertheless, a recently introduce method, called Hu-mAb (39), showed a slightly better correlation with these ADA data (R= - 0.58)(39). Hu-mAb is a random-forest classifier trained in a supervised way to differentiate human from mouse sequences. As supervised learning is well known to typically outperform unsupervised learning, and as the ADA dataset contains only human, mouse, chimeric, or humanised antibodies from mouse precursors, it is perhaps not surprising that a supervised learning approach specifically trained to separate mouse from human antibodies shows a slightly stronger correlation with these data. In this work, we chose to develop a model trained with unsupervised learning because we want it to be applicable to any input Fv sequence, as opposed to just mouse and human sequences. One of the main reasons we developed AbNatiV is to use it in synergy with emerging approaches of de novo antibody design, which typically yield artificial sequences whose latent distribution may be specific to the design method employed.

Alongside humanness, AbNatiV quantifies the nativeness of nanobodies. The resulting model exhibits high classification performance in distinguishing VHH sequences derived from camelids from VH sequences from other species and from PSSM-generated artificial VHH sequences. The ability to discriminate artificial sequences confirms that the correct classification of VHHs does not solely rely on the presence of nanobody hallmark residues (42), as these are also present in the artificial PSSM-generated VHH sequences. However, while the discrimination performance of native nanobody sequences from artificial ones is excellent, it is not as good as that of AbNatiV trained on human sequences (PR-AUC of VHH: 0.942, VH: 1.000, Vκ: 0.992, and Vλ: 0.990). This observation may suggest that a bigger, and especially more diverse, VHH training dataset could be beneficial. While AbNatiV-VHH is trained on slightly more sequences than AbNatiV-Humanness, these come from a much more restricted number of studies. Therefore, our VHH dataset has more limited diversity that the human one and it also comprises nanobodies from different camelid species (llamas, dromedaries, vicugna, etc.; **Supplementary Table 5**), which may slightly confuse the model and demand for a larger training dataset. Quite generally, we expect that the publication of additional camelid immune repertoires will be beneficial for data-driven approaches like AbNatiV, which have the potential to facilitate and accelerate nanobody development and humanisation.

AbNatiV can also be used to assess whether CDR loops are in the right context or not (**Fig. 4D, E)**. This observation demonstrates the ability of the model to capture long-range interactions between CDRs and framework regions and shows that AbNatiV can assist CDR grafting. For example, the CDR nativeness loss calculated by AbNatiV is consistent with the experimentally observed loss of binding affinity upon CDR grafting in a different framework (**Fig. 4D**). Yet, a quantitative correlation with the magnitude of the change in K_D_ is not observed, most likely because only a subset of non-ideal CDR-framework contacts resulting from grafting actually translates to an affinity loss, in a way that is highly specific to the nanobody-antigen binding pose. We envisage that these applications of AbNatiV may increase the effectiveness and success of *de novo* antibody design methods based on the grafting of designed CDR loops (20,21,59). We have focussed our analysis on VHH sequences. However, the exact same approach can be carried out with AbNatiV-humanness to select human scaffold sequences that serve as better receptors for CDR grafting from non-human sources, such as murine CDRs (see **Fig. 1D**), designed CDRs, or CDRs from a synthetic library.

Nanobodies exhibit significant structural differences from human VH domains that enable them to fold independently of a VL counterpart. For instance, the CDR3 of nanobodies is often longer and sometimes folds back to interact with the framework (5,45). During the process of humanisation for therapeutic purposes, it is crucial to improve humanness while preserving these traits, as they translate into high stability and binding affinity. Consequently, we introduce an automated humanisation pipeline that combines the humanness and VHH-nativeness assessments of AbNatiV. We applied this deep-learning-based dual-control strategy on two nanobodies and showed that the humanised variants generated with the Enhanced Sampling pipeline retain their binding activity and biophysical stability. Conversely, both properties are disrupted when conventional structural and residue-frequency humanisation is applied to the same nanobodies.

We selected Nb24 and mNb6 as test nanobodies because they bind two distinct antigens with therapeutic potential, are quite different from each other (for example Nb24 has a non-canonical disulphide and mNb6 has not) and represent respectively a standard and a very challenging test case for humanisation. Nb24 was obtained from immunisation, and with a mid-nanomolar dissociation constant is not a particularly optimised nanobody. Conversely, with a high picomolar dissociation constant, mNb6 is a highly affinity-maturated version of a nanobody (Nb6), which was obtained from the screening of a synthetic library (53). Consequently, one would expect that mutations in mNb6 may be more likely to disrupt affinity and stability than mutations in Nb24. Indeed, our results neatly align with this hypothesis, with both Enhanced and Exhaustive sampling strategy showing excellent results on Nb24, improving both binding affinity (marginally) and stability (substantially). Conversely, only the Enhanced sampling strategy didn’t compromise the binding of mNb6 retaining a comparable stability.

Overall, the Enhanced sampling AbNatiV humanisation yielded the most promising results. Additionally, this sampling approach is the most computationally efficient, adding to its value. Yet, the Exhaustive sampling remains a valuable choice as it generates humanised sequences for different numbers of mutations via its Pareto set selection (see Methods). In our experiments we have tested only the variant with the highest VH-humanness, which is also the one with the highest number of mutations except for the Exhausted +buried strategy ran on mNb6 (**Supplementary** Fig. 19). Yet, this approach offers users the flexibility to pick humanised variants with fewer mutations, lowering the risk of affecting their activity or other biophysical properties. Moreover, we make all the sampling parameters fully adjustable in our software (e.g., tolerance of humanness, VHH-nativeness decrease or buried residues, see Methods). Users can also look at the AbNatiV residue profiles and make in-depth analysis of the expected impact of humanisation. This empowers users to make fully informed decisions when designing their humanised sequences and selecting those for experimental testing.

In addition to nanobodies, AbNatiV can be used to humanise directly paired heavy and light Fv sequences by running the same sampling strategies without the VHH-nativeness constraint. In this way, the pipeline improves both heavy- and light-chain humanness. This option is made available online on our repository.

Finally, we note that the trained AbNatiV models may facilitate applications of semi-supervised learning, even if we have not explored this avenue in this work. Semi-supervised learning, also known as low-N learning, combines a small amount of labelled data with a large amount of unlabelled data during training (60–62). The embedding of the VQ-VAE, and possibly also the last hidden layer of the decoder, can be seen as an effective way to distil the fundamental features of antibody variable domains into a representation that is semantically rich and structurally, evolutionarily, and biophysically grounded (63). The compactness of this representation, and the fact that it was built by learning from many functional sequences, means it can be used as input to train a supervised model (top model) with few free parameters, which therefore may be expected to generalise with relatively few labelled training data (61). Approaches of semi-supervised learning with protein directed-evolution data have recently been successfully deployed and were shown to be able to generalize to unseen regions of sequence space (60,62,64).

In summary, we expect that AbNatiV will facilitate antibody and nanobody development, as it provides a rapid, highly accurate, and interpretable way to quantify humanness and VHH-nativeness from the knowledge of the sequence alone. Looking into the future, it is reasonable to expect that computational approaches of de novo antibody design will be increasingly adopted to generate novel antibodies. In this context, AbNatiV provides a holistic way to select the best designed antibodies or nanobodies to target epitopes of interest, for instance by ensuring high humanness or by facilitating the selection of a framework highly compatible with designed CDR loops. Antibodies designed in this way will have high nativeness, and therefore can be expected to share similar specificities and in vivo properties as immune-system-derived antibodies. Besides low immunogenicity, these properties include favourable half-life and low self-antigen cross-reactivity, which are essential for successful clinical development. Overall, we believe that approaches like AbNatiV will constitute a step-change in our ability to design de novo antibodies with in vivo properties highly competitive with those of antibodies isolated from immune systems.

## 5 Methods

### 5.1 Datasets and antibody sequence processing

The source of all antibody sequences used for training and testing is given in **Supplementary Table 5**, with the full-length antibody sequences coming from the Observed Antibody Space (OAS) (36), and the single-domain camelid VHH sequences coming from various studies (65–68). All sequences were aligned, cleaned, and processed beforehand. Non-redundant sequences were aligned using the AHo numbering scheme (54) resulting in aligned sequences of length 149. The alignment was carried out using the widely employed ANARCI software (69) followed by a custom python script to check for consistency and fix misalignments. More specifically, we found that in some instances gaps may be opened in unexpected positions (sometimes in framework 1 or framework 2) leading to a misalignment of the subsequent part of the sequence, including the fully conserved cysteines that form the intra-domain disulphide bond (AHo positions 23 and 106). Therefore, a script was run to adjust possible inaccuracies in the alignment of each sequence within the multiple sequence alignment (MSA). This script maximises the identity between the MSA consensus sequence and the sequence under scrutiny calculated at all positions with conservation index greater than 0.9, which include the two fully conserved cysteines. Sequences whose alignment could not be fixed, or that didn’t have two cysteines at the conserved positions (because of e.g., sequencing errors) were discarded. Furthermore, Fv sequences with more than one or two missing residues at respectively the N- and the C-terminal were removed. For heavy chains, a Glutamine residue was added at the N-terminus, if missing, and two Serine residues were added at the C-terminus, if missing. For lambda and kappa light chains, respectively a Leucine or a Lysine were added at the C-terminus (AHo position 148), if missing. After alignment a check for unique sequences was repeated (because for example after completing the C-terminus some duplicated sequences may exist) and any duplicate discarded.

Datasets of processed heavy, lambda, kappa (from human, rhesus, and mouse) and VHH antibody sequences from various studies from the literature were assembled (**Supplementary Table 5**) and processed as described above. All the parsed sequence datasets used in this study are available online in the AbNatiV GitLab at https://gitlab.developers.cam.ac.uk/ch/sormanni/abnativ.

#### 5.1.1 Training, Validation, Test and Diverse datasets

2,000,000 sequences from the human heavy, lambda, and kappa databases were used to train three distinct models, respectively. 2,144,185 sequences from the VHH databases (Camelid and PDB-sdAB) were used to train a fourth model. For each model, 50,000 sequences were additionally kept aside for validation, and 10,000 sequences for testing. These training, validation, and test sequences were selected as random splits from the larger database of unique sequences. As we only have unique aligned Fv sequences, this procedure ensures that sequences in training, validation, and test datasets are at least one mutation away, as commonly done in the field when dealing with large databases of sequences.

Furthermore, to be able to assess performance on a dataset of sequences that are more distant from any training sequence, we have built an additional diverse dataset for each model. Such diverse datasets are compiled with sequences that are at least 5% different from any sequences of the training set (2.5% for Vκ and Vλ, as light chains have less diversity). Percent difference is defined as the number of mutations between an aligned test sequence and an aligned training sequence (gap to gap is not considered a mutation), divided by the length of the gapless test sequence. As calculations are memory-intensive (for instance the VH test set vs. training set only would be 2x10^10^ sequence difference value, meaning that each boxplot would need more than 80 Gb of memory), they are carried out in the following way. First, the distance between all sequences in the Training set and each sequence in the Test set (or in any other of the datasets on the x-axis) is calculated. This corresponds to 2 million percent differences (more for VHHs, see **Supplementary Table 5**). Then, only the values of the minimum and of the 5^th^, 25^th^, 50^th^ (i.e., median), 75^th^, and 95^th^ percentiles of these differences are saved. At the end of the calculation, these values are available for each sequence in the Test set, and the same goes for the other datasets examined (x-axis). For the human models (VH, Vκ, and Vλ) diverse sequences are extracted from both Test and BioPhi datasets (subset of the training dataset of the Sapiens transformer from BioPhi (37), see **Supplementary** Fig. 20) to yield the corresponding Diverse >5% (or >2.5% for the light chains) dataset. For the VHH model, diverse sequences are extracted from the Test dataset by requiring at least 5% difference from the closest sequence in the training set **Supplementary** Figure 20 shows the cumulative distribution functions (CDFs) of the minimum percent different to training sequences for each dataset. **Supplementary** Figure 4 the distribution of the sequence difference between training sequences and all sequences in the datasets used to assess AbNatiV performance, as well as the lack of correlation between the AbNatiV nativeness score and the distance of that sequence from the training set.

#### 5.1.2 PSSM-generated datasets of artificial sequences

Position weight matrices (PWM) and corresponding position-specific scoring matrices (PSSMs) were computed from each human and camelid antibody training datasets (**Supplementary** Fig. 7). From these matrices, additional custom datasets of artificial sequences were generated to be used as controls, named PSSM-generated datasets. These sequences were built by randomly filling each residue position using the underlying residue frequency observed in the PWM (that is the matrix of observed residue frequencies, **Supplementary** Fig. 7) considering only those amino acids enriched at that position (i.e., PSSM log-likelihood score > 0).

### 5.2 The AbNatiV model

#### 5.2.1 Vector-quantized variational auto-encoder (VQ-VAE) architecture

The AbNatiV model takes aligned antibody sequences of length 149 as input, and one-hot encodes each into a tensor of dimension 149x21. Each position is represented by a vector of size 21 consisting of zeros and a one at the alphabet index of the residue under scrutiny (20 standard amino acids and a gap token).

The architecture of the models is based on a vector-quantized variational auto-encoder (VQ-VAE) framework (33), which involves a VAE with a discretisation of the dense latent space through code-vectors (**Fig. 1A**). The sequence input *x* ∈ {0,1}^149×21^is first encoded into a compressed sequence representation *z_e_*(*x*) ∈ ℝ^(*l*×*d_c_*^, where *l* represents the compressed sequence length and *d_c_* the dimension of the code-vectors. In order to discretise *z*_*e*_(*x*) in the latent space, a learnable codebook of *N* code-vectors 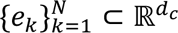 is used. A nearest neighbour lookup is applied, so that each component 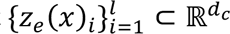 is substituted by the closest code-vector of the codebook, resulting in the quantised embedding *z_q_*(*x*) ⊂ ℝ*^l×d_c_^*. Finally, *z_q_*(*x*) is decoded to generate the reconstructed output *x*^^^ ∈ {0.1}^149×21^ having the original dimensions as the original sequence input *x*.

For increased codebook usage (i.e., higher perplexity), the *N* code-vectors are initialized with the *N*

*k*-means centroids of the first training batch, and code-vectors not assigned for multiple batches are replaced by randomly sampling the current batch as detailed in Ref. (70), where a vector quantizer was applied to sound compression. In addition, the code-vectors 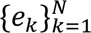 and the encoded inputs *z_e_*(*x*) are *l*_2_-normalised. The Euclidean distance of the *l*_2_-normalised vectors is used during the nearest neighbour lookup resulting in a cosine similarity search as proposed in the image modelling model ViT-VQGAN (71). Furthermore, the code-vectors from the codebook are updated during training by exponential moving average (EMA) with a decay of 0.9 to assure a more stable training (72).

The encoder and decoder layers are illustrated in **Figure 1B**. In the encoder, the input sequence is embedded by a patch convolutional layer (71). A 1D-convolution layer with a kernel size *K* equals to its stride *S* embeds each of the non-overlapping patches of dimension *K*x21 into a single vector of size *d_emb_* (i.e., the number of channels of the 1D-convolution layer). A minimal padding was added to the sequence input beforehand to avoid missing any sequence region. For instance, in the VHH model, with *K* = *S* = 8, a padding of 3 is added to compress the sequence inputs into *l* = 19 embedding vectors of size *d_emb_*. Then, a sinusoidal positional encoding is added before *L* transformer blocks. The transformer blocks are designed as in BERT (73), with *H* heads in the multi-head attentions layer and a hidden dimension *d_ff_* in the feed forward layer. Before quantisation, a linear layer is applied to reduce the embedding dimension *d_emb_* to the size of the code-vectors *d_c_*.

In the decoder, a linear layer is first applied to augment the dimension of the discrete embedding *z_q_*(*x*) to *d_emb_*. Mirroring the encoder, a positional encoding is applied before *L* transformer blocks with the same hyperparameters of the encoder. Ultimately, a transpose 1D-convolution layer with a softmax activation function is applied to reconstruct back the tensor into the same dimension of the original sequence inputs. All the hyperparameters were manually tuned for the VH and VHH models. It has been found empirically that the same hyperparameter values lead to the best performances for both models. Since the hyperparameters do not look to be dependent on the origin of the training set, the same hyperparameter values were used across all models, and their values are given in **Supplementary Table 6**.

#### 5.2.2 Unsupervised masked learning

Like the original VQ-VAE (33) the AbNatiV models are trained to minimize a negative evidence lower bound (NELBO) consisting of three terms as follows:

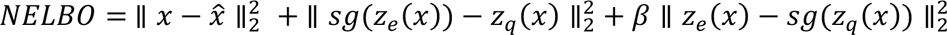

The first term is the negative log-likelihood reconstruction loss, which is characteristic of the variational autoencoders. This term is approximated by the reconstruction mean-squared error (MSE) between the input *x* and the decoder output *x*^^^. The second and third terms are associated with the vector quantisation step in the latent space, enabling the codebook to get trained. Both terms are MSEs between the encoded input *z*_*e*_(*x*) and the quantised latent embedding *z*_*q*_(*x*). In particular, in the second term, stop gradient *sg* is applied to *z*_*e*_(*x*) to detach it from the computational graph, thereby updating only the codebook during back propagation. In the third term, *z*_*q*_(*x*) is conversely ignored during back propagation, which drives the encoder to commit to the codebook vectors. The stop gradient allows the codevectors and the encoder to be updated at different speeds. The relative learning speed between these two terms is imposed by the scaling factor β. In all our models, *β* is set to 0.25. By choosing *β* < 1, the codevectors are updated more rapidly to align with the encoder, preventing an arbitrary growth of the encoder outputs (33).

The neural network is implemented using PyTorch.1.14 (74) and enhanced by the PyTorchLightning.0.7 module. The models are trained with a batch size of 128 by the Adam optimizer (75) with a learning rate of 4e-05. During training, a masking is applied to the one-hot encoded inputs. As in the training of the language transformer model BERT (73), a percentage of positions *p*_*mask*_ is selected for masking. Among these selected positions, 80% are replaced by the uniform vector of size 21 with a probability of 1/21 for each residue, which we use as a mask token. 10% are randomly replaced by another residue or gap. 10% remain unchanged so that the model does not learn to expect a fixed number of masked residues (as all sequences are aligned to 149 positions).

### 5.3 Training with non-aligned sequences

For comparison, we trained the same VQ-VAE architecture (same hyperparameters and number of training epochs) on non-aligned VH sequences. A padding of value 0 has been added to the left and right of the one-hot input vectors of non-aligned sequences to reach a size of 149. If the padding size required is odd, one more pad is added to the right side. The loss function is identical. For the reconstruction accuracy, only the non-padded components are considered.

### 5.4 Antibody nativeness definition

The concept of antibody nativeness is intuitively understood as the extent to which a given sequence resembles those of native antibodies, that is of antibodies derived from the immune system under scrutiny (in this work human or camelid immune systems). Here, we provide a quantitative definition of nativeness as:

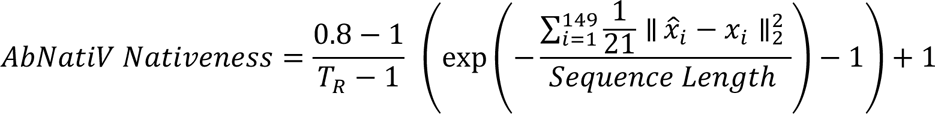

where 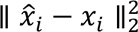 is the MSE at sequence position *i* between the aligned input sequence *x* and the reconstructed output sequence *x*^^^ of a trained AbNatiV model. This MSE is summed over all 149 positions of the aligned sequence and normalised by the length of the input sequence (i.e., without considering the gaps opened by the alignment). As this operation gives a number *X* that in principle ranges in [0, +∞[, where 0 would correspond to a fully native sequence that is perfectly reconstructed, we apply the function *Y* = exp (−*X*). This way, *Y* is now a number in [0,1], where 1 means fully native, thus providing a more intuitive ranking for high and low nativeness. We wish to point out that, for typical antibody sequences from any species, the average MSE *X* is typically a very small number in all the models that we trained. Therefore, in this relevant range of *X*, *Y* = exp (−*X*) is effectively approximated by a simpler linear transformation *Y* = 1 − *X* meaning that the distance between different antibody sequences is only minimally affected by the exponential transformation. Finally, the operation (0.8 − 1) ∗ (*Y* − 1)/(*T_R_* − 1) + 1 linearly rescales the scores so that the final nativeness score becomes a quantity directly and intuitively interpretable as an absolute value for a single sequence, and not just usable to rank different sequences (**Supplementary** Fig. 21). *T_R_* is specific to each trained model, and it denotes the optimal threshold of *Y* that best separates native sequences (positives in the classification) from non-native sequences (negatives in the classification). This linear transformation rescales the values of *Y* so that this threshold on the final nativeness score becomes 0.8 for every model. In other words, this means that a nativeness score greater than 0.8 denotes a sequence classified as native, while a score below 0.8 one classified as non-native. *T_R_* is calculated for each trained model as follows. The precision-recall (PR) curves are generated between human sequences (Human Test & Human BioPhi datasets) as positives, and non-human sequences (Mouse) as negatives for the VH, Vκ and Vλ models. Similarly, the PR curve is also calculated between VHH sequences (Camelid Test) as positives, and non-VHH sequences (Human Test and Mouse) as negatives, all computed on the *Y* = exp (−*X*) scored sequences (**Supplementary** Fig. 21A, 21D, 21G, 21J). For every model, the PR optimal threshold value *T_R_* is extracted as the point closest to (1,1) (**Supplementary** Fig. 21B, 21E, 21H, 21K, *T_R_*(*VH*) = 0.988047, *T_R_*(*VKappa*) = 0.992496, *T_R_*(*VLambda*) = 0.985580, and *T_R_*(*VHH*) = 0.990973). The scores are thus linearly rescaled to shift *T_R_*to 0.8 to return a final value ∈]−∞, 1] for any input Fv sequence (**Supplementary** Fig. 21C, 21F, 21I, 21L). Not only does this rescaling make the nativeness scores from different models interpretable in the same way, but it also future proofs the definition of nativeness. The values of Tr will change if, in the future, the model is retrained on a larger or more diverse dataset, or if the architecture is further improved. However, the interpretation of the final nativeness score, which is what users will rely on, will be the same. We define AbNatiV humanness score the nativeness from AbNatiV trained on VH, Vκ, and Vλ human sequences, and AbNatiV VHH-nativeness score, that from AbNatiV trained on single-domain VHH sequences.

In addition, residue level scoring profiles are defined by applying *Y* = exp (−*X*) to the MSE reconstruction error at each position of the given sequence.

### 5.5 Performance metrics

All the performance metrics reported are computed by analysing 10,000 scored sequences for each database, except for the Diverse datasets (see **Supplementary Table 5**). For datasets smaller than 10,000, the whole dataset is used.

#### 5.5.1 Classification

The area under the curve (AUC) of the receiver operating characteristic (ROC) and of the precision-recall (PR) curves are computed to quantify the ability of a model to classify sequences. For ROC curves, the AUC is equal to 1 when the classification is perfect. It is equal to 0.5 when the model performs as poorly as a classifier that is randomly sampling from a uniform distribution. For PR curves, the AUC is also equal to 1 when the classification is perfect, while it is equal to the ratio of positive entries over the total number of entries in the datasets when the classification is random.

#### 5.5.2 The amino acid reconstruction accuracy

The amino acid reconstruction accuracy quantifies the ability of a model to reconstruct the initial unmasked input from the embedded vector of the latent space. The reconstructed outputs of the model have for each position a probability distribution over the alphabet. For each position, the most probable amino acid is selected. The amino acid reconstruction accuracy corresponds to the ratio of correctly predicted residues for every position over the length of the sequence. It is equal to 1 if all residues have been correctly reconstructed, and 0 if not even one has. It can be expressed, as follows:

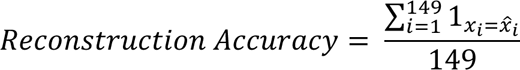

where *x_i_* and *x_i_*^^^. are residue at the position *i* of respectively the input *x* and the reconstructed output *x*^^^ of the model.

#### 5.5.3 Benchmarking with other assessments from the literature

Open-source antibody humanness assessments from the literature were employed to benchmark the performances of AbNatiV. These assessments include OASis and Sapiens from Biophi (37) and AbLSTM (38).

OASis is an average 9-mer peptide similarity searched through the OAS database. Sapiens is an unsupervised human antibody language model based on the transformer encoder BERT (73) network. It is trained on unaligned human antibody sequences from the OAS database. The GitHub implementation (https://github.com/Merck/BioPhi) of OASis and Sapiens is used to score our testing databases. The relaxed stringency level is used for the OASis assessment. The OASis score is not position discrete; hence it cannot be used for the amino acid reconstruction task.

AbLSTM (38) is an unsupervised long-short-term-memory (LSTM) neural network. Human heavy chains sequences from the OAS database are aligned prior training. Here, we used the pre-trained model in the benchmarking, and we also retrained the AbLSTM for 10 epochs from scratch on the same single-domain, and human heavy, lambda and kappa databases used for the training of our VQ-VAE models. In the case of human VH we carried out the benchmark with both retrained AbLSTM and original pre-trained one as downloaded from https://github.com/vkola-lab/peds2019. The original hyperparameters of AbLSTM were used (embedding dimension = 64, hidden dimension= 64, batch size = 128, and learning rate = 2e-03). The negative log sum loss of the AbLSTM model was employed as its humanness or VHH-nativeness scores as done in the original work (38).

### 5.6 Predictions on antibody therapeutics

549 antibody therapeutics from the IMGT database (76) were obtained from the BioPhi dataset (37). This dataset includes 196 fully human therapeutic sequences and 353 therapeutics of non-human origin (mouse, chimeric, and humanised). The AUC of ROC and PR curves are computed to quantify the ability of the models to separate these two groups of sequences.

Similarly, 216 antibody therapeutics with their immunogenicity scores – expressed as the percentage of patients who developed an anti-drug-antibody (ADA) response during clinical trials – were also obtained from the BioPhi dataset (37). These sequences were used to quantify the extent of correlation between the models nativeness scores and the observed ADA response, using the Pearson correlation coefficient and its associated p-value. For each therapeutic, the mean between the scores of VH and VL domain is used as an overall nativeness.

The humanness scores from different methods developed to humanize antibodies with which we compare our approach were obtained as computed by the authors of BioPhi and deposited in their GitHub (https://github.com/Merck/BioPhi) and in the tables of Ref. (37). The alternative methods considered in this work are the BioPhi germline content (37) (sequence identity to closest human germline), HumAb (39) (random forest-based humanness), IgReconstruct (77) (positional nucleotide frequency scoring from back-translated human antibodies), AbLSTM (38), T20 (78) (similarity average among the closest 20 sequences), and Z-score (79) (similarity average across all sequences) assessments. Light-chain-only antibodies (i.e., Istiratumab, Lulizumab Pegol, Placulumab and Tibulizumab) are removed from the IMGT BioPhi parsed dataset as the original pretrained AbLSTM can only scores heavy chains. Because the Fv sequence of Pexeluzimab has missing C-term residues, it is also removed from the ADA dataset and excluded from further analysis. All these sequences with their associated scores are available in **Supplementary Datasets 1** and **2**.

### 5.7 Grafting assessment on nanobodies

In Ref. (41) all three CDRs of 6 nanobodies were grafted onto a camelid VHH framework sequence, referred to as universal framework (UF). Binding K_D_ and conformational stability ΔG were experimentally measured for all six wild type (WT) nanobodies, and corresponding variants with CDRs grafted onto the universal framework. Here, we compute the nativeness scores of the 6 pairs of WT and grafted nanobodies. As the UF has intrinsically better nativeness because of its ideal framework, to understand whether our model predicts the CDRs to be in the right context or not, we compute the VHH-nativeness CDRs scores. These are defined as the sum of the MSE reconstruction scores of all residues at the CDR positions (according to the AHo numbering scheme) normalised by the length of these CDRs without gaps. *Y* = exp (−*X*) is applied to the resulting sum *X* to give a more interpretable number in [0,1]. A nativeness prediction of a CDR context is considered correct when the VHH-nativeness CDRs score of the WT nanobody is higher than that of its UF-grafted counterpart, as reflected by the experimentally measured change in binding K_D_ which is typically worse for the UF-grafted variant (**Fig. 4D**).

We also carried out this assessment on a much bigger scale, by computationally grafting all CDRs of 5,000 different Nbs from the Camelid Test dataset onto the UF scaffold.

### 5.8 Humanness assessment of nanobodies

For the analysis reported in **Figure 5**, 300 VH human sequences and 300 camelid sequences from the test datasets are scored both with the AbNatiV human heavy and camelid heavy models to provide background distributions. Then, we further scored 8 WT nanobodies from a SARS-CoV-2 study (47) and their humanised counterpart as reported in Ref. (45), and 3 therapeutic nanobody sequences (Envafolimab, Caplacizumab, and Rimteravimab) available from the therapeutic database Thera-SAbDab (80).

### 5.9 Automated humanisation of nanobodies

The humanisation process of nanobody sequences by AbNatiV follows a dual-control strategy which seeks to increase the humanness while retaining the VHH-nativeness of a given sequence. Standard antibodies can be humanised exactly as described here, by removing all steps involving the VHH-nativeness.

Given an input sequence, the VH-AbNatiV and VHH-AbNatiV residue profiles are computed along with the solvent accessible surface area (SASA) using the “rolling ball” algorithm (81) on the whole unbound structure modelled with NanoBuilder2 from the ImmuneBuilder software (51). The SASA of each residue is converted into a relative SASA (RASA) value by dividing the SASA of the given residue X under scrutiny with its maximum allowed SASA (82). The latter is obtained as the SASA of residue X in the context of the Gly-X-Gly tripeptide in a fully extended conformation. Structural modelling and SASA calculations are only performed when the user choses to do framework resurfacing, that is to avoid mutating any buried residue, which is the default behaviour.

To reduce the mutational space, we first flag positions for mutation using the residue nativeness profiles. The search is restricted to the framework region, as CDRs typically contain binding residues. Flagged positions have either a VH-AbNatiV or VHH-AbNatiV score smaller or equal to 0.98, or the WT residue is not enriched in the human VH PSSM (i.e., does not have a PSSM log-likelihood score > 0 and a PSSM frequency > 0.01, see **Supplementary** Fig. 7). The latter condition is just an additional safeguard. In our investigations we have never observed a framework residue that was not enriched in human VH PSSM and yet was not flagged as liability by the VH-AbNatiV profile. Furthermore, if framework resurfacing is selected as an option, mutable residues must exhibit a RASA greater or equal to 15%. By comparison, in Chen et al. work (83) a RASA of 20% serves as a cut-off between buried and exposed residues. Starting from these automatically identified mutable positions, we developed two distinct sampling methods to explore the mutational space.

#### 5.9.1 Enhanced sampling

The enhanced sampling is illustrated in **Supplementary** Figure 14**.A**. Convergence towards the best combination of mutations is achieved by mutating each position subsequently one a time, as opposed to exploring all possible combinations. The order at which positions are mutated is defined starting from those mutable positions that are least affected when other positions are mutated. This strategy increases the odds that positions mutated early remain stable even after subsequent mutations along the sequence are performed, leading to a more efficient path towards identifying the best mutational variant. Thereby, a first calculation is performed to sort positions to mutate based on their average interdependence upon mutations at every other position in the sequence. To quantify this dependence, a computational deep mutation scanning is implemented. For a given position, each of the other positions is individually mutated into all available amino acid residues (19 possibilities). For each mutation, and each of the other positions, we calculate the difference between the AbNatiV VHH residue score at the position under scrutiny of the WT sequence and that of the mutated sequence (note, mutations are at other positions but may still affect the score of this position and this is what we are probing for here). These differences are then averaged into a single value quantifying the dependence of the position under scrutiny on mutations elsewhere in the sequence. This procedure is iterated for every liable position.

Subsequently, starting from the position with the least dependence on mutations at other positions, we mutate it with all the amino acids significantly enriched in the human VH PSSM (i.e., with a PSSM log-likelihood score > 0 and a PWM frequency > 0.01, see **Supplementary** Fig. 7) as shown in **Fig. 6.B**. We exclude cysteines and methionines from the list of candidate mutations as these are linked to developability liabilities. The selected mutation at each position is then the one which increases most the multi-objective function: 0.8 Δ *VH* + 0.2 Δ *VHH* and which does not decrease the VHH-AbNatiV score by more than 1.5% of that of the WT (i.e., 1.5% decrease tolerance for ΔVHH). If no such mutation is found (e.g., all screened ones decrease the VHH-nativeness by more than 1.5%), the residue is left to WT and the procedure continues to the next mutable position. If a mutation is found, the sequence is updated and the process of selecting positions for mutation in **Figure 15.A** recommences from the beginning to ensure that no over other positions has become a liability (i.e., residue score <= 0.98) following the introduction of this new mutation.

#### 5.9.2 Exhaustive sampling

The exhaustive sampling is illustrated in **Supplementary** Figure 14**.B**. We generate all the possible combinations of mutations at all liable positions by considering as candidates for each position those amino acids significantly enriched in both human VH and VHH PSSMs (i.e., with a PSSM log-likelihood score > 0 and a PWM frequency > 0.01, see **Supplementary** Fig. 7). Cysteines and methionines are excluded from the list of candidates as these are linked to developability liabilities. The WT residue is retained in the list of candidate amino acids at each liable position. First, we retain only those combinations of mutations that do not decrease the VHH-nativeness score by more than 1.5% over that of the WT. Then, we compute the Pareto front that maximises the VH-humanness score while minimising the number of mutations over all remaining combinations of mutations. In fact, given that WT residues were retained in the list of candidate amino acid substitutions, the method produces mutational variants that have a number of mutations ranging from 0 (the WT, which is one possible combination) and the total number of identified liable positions.

At the end, this approach returns a set of mutational variants with the highest VH-humanness for each number of mutations that are beneficial to the VH-humanness (see **Supplementary** Fig. 19). In the pareto analysis, increasing the number of mutations is beneficial only when it further increases the VH-humanness score. For instance, we see in **Supplementary** Figure 19**.D** that going from 9 to 10 mutations does not increase the VH-humanness further, and therefore the variant with 10 mutations is not selected in the Pareto front. In this work, experimental testing was conducted exclusively on the sequence exhibiting the highest humanness score, which happens to be the one with the highest number of mutations in all Exhaustive sampling designs except for the variant in **Supplementary** Figure 19**.D**.

#### 5.9.3 Frequency- and structure-based nanobody humanisation

To provide a benchmark for the AbNatiV humanisation pipelines described above, we carried out nanobody humanisation also using the recently introduced Llamanade humanisation pipeline (45). This approach builds on a systematic analysis of the sequence and structural properties that distinguish nanobodies from human VH, and proposes humanising mutations based on the analysis of the input nanobody modelled structure and the key differences between its sequence and sequences of human VH domains. These frequency- and structure-based designs were carried out with the Llamanade webserver accessed on 4^th^ of July 2023 (at http://35.208.211.136).

### 5.10 Protein production

Genes encoding the Nb24 and mNb6 WT nanobodies and their humanised variants were synthesized and cloned into an isopropyl-β-D-thiogalactopyranoside (IPTG)–inducible vector (by Genscript in vector pET29a(+)), including a leading PelB sequence to enable translocation to the periplasm, facilitate intra-domain disulphide bond formation, and ultimately the secretion of the protein to the expression media. A C-terminal 6x His tag is added for purification. All expressed amino acid sequences are given in **Extended Data Table 3**. Care was taken to maintain the same codon usage as the WT, except for the mutated amino acid positions. Plasmids were transformed into E. coli Shuffle LysY strain to further facilitate the formation of the disulphide bond, and to enable the secretion to the expression media (which is facilitated by the LysY leakier cell wall). Cultures (0.5 litre) of LB media were inoculated at initial 0.03 OD600 (optical density at 600 nm), grown at 37°C until reaching 0.8 OD600 nm, and then induced with 500 µM IPTG at 30 °C for overnight expression.

His Mag Sepharose Excel magnetic beads (Cytiva) were washed in PBS and added to the cultures (1 mL per 0.5 Litre) about 3 hours before harvesting to capture the secreted his-tagged nanobodies. Loaded beads were then fished out from the expression media using an AmMag™ magnetic wand (Genscript) and purification was performed with an AmMag™ SA Plus Semi-automated System (Genscript) using PBS as running buffer and carrying out washing steps with PBS 4 mM Imidazole, and elution with PBS 200 mM imidazole. Eluted nanobodies were further purified by size exclusion chromatography using a Superdex 75 10/300 column equilibrated in PBS on an Akta Pure System (Cytiva) to remove the imidazole, further increase the purity, and isolate monomeric nanobodies.

Purified nanobodies were aliquoted, flash-frozen in liquid nitrogen, and stored at -80 °C. Each aliquot was used only once, and, following thawing, was centrifuged at 21,000 g at 4°C for 10 minutes to pellet down any precipitate that may have formed during freeze/thawing.

Recombinant β_2_-microglobulin was expressed and purified to homogeneity as previously reported in (84). Briefly, *E. coli* BL21(DE3) cells were transformed with pET29b carrying the coding sequence of β_2_-microglobulin. The transformed cells were grown at 37 °C in LB medium supplemented with kanamycin and protein expression was induced with 1 mM IPTG for 3 h. β2-microglobulin was purified from the inclusion bodies. The cell pellet was resuspended in Triton buffer (100 mM sodium phosphate pH 7.4, 0.1% Triton, 1 mM EDTA, 10 mM DTT) supplemented with lysozyme and Dnase. The cells were lysed by sonication and then centrifuged. The pellet obtained was washed with Triton buffer and then dissolved in 6 M GuHCl. β_2_-microglobulin was refolded by consecutive dialysis (20 mM Sodium phosphate pH 7.4, 150 mM NaCl; 20 mM Sodium phosphate pH 7.4, 75 mM NaCl; 20 mM Sodium phosphate pH 7.4, 35 mM NaCl and 20 mM Tris HCl pH 8.3) and then purified by ion exchange using a Hi Prep Q FF 16/10 column (GE Healthcare Life Sciences) connected to an Akta Pure system (Cytiva). The protein was eluted with a linear 0-1 M NaCl gradient in 20 mM Tris-HCl pH 8.3. Purified β_2_-microglobulin was aliquoted, lyophilized and stored at -80 °C. SarS-CoV-2 RBD was purchased as biotinylated purified protein from CUSABIO (product code CSB-MP3324GMY1-B) and stored at -80 °C.

Protein concentrations were measured using blanked absorbance 280 nm values and extinction coefficients calculated from the amino acid sequence using the Expasy ProtParam tool ((web.expasy.org/protparam/).

### 5.11 Liquid chromatography–mass spectrometry

The mass of all antibodies was verified by LC-MS using an ACQUITY UPLC/VionTM-IMS-QTof system coupled with an electrospray ionization (ESI) source. Liquid chromatographic separation of samples was performed on ACQUITY UPLC Protein BEH C4 column (300 Å pore diameter, 1.7 μm, 2.1 mm × 50 mm, Waters) using gradient elution. 1 ul of sample was injected with a flow rate of 0.3 mL/min and the analysis was carried out at defaulted parameters. The acquired data was processed using UNIFI ™ software. Disulphide bonds (-2 Da per bond) were detected in all variants (see **Extended Data Table 3**).

### 5.12 β2-microglobulin biotinylation

To enable BLI binding assays with streptavidin sensors, β_2_-microglobulin was biotinylated. 10 µM of β_2_-microglobulin were incubated with 1x molar concentration of EZ-Link Sulfo-NHS-LC-Biotin (Thermofisher 21335) for 2 hours, quiescent at room temperature. After this time, unreacted biotin was removed by size exclusion chromatography using a Superdex 75 10/300 column equilibrated in PBS on an Akta Pure System (Cytiva). Biotinylated β_2_-microglobulin was then characterised with LC-MS do determine the degree of labelling (**Supplementary** Fig. 22).

### 5.13 Measurements of thermal stability

Measurements of apparent melting temperature were carried out in PBS at 6 µM nanobody concentration (except for mNb6 Exhausted Sampling + buried, which was at a concentration of 1.5 µM because of insufficient material) on a Tycho system (Nanotemper). Each experiment was repeated 3 times for Nb24 variants and 2 times for mNb6 variants. Each 350/330 fluorescence ratio trace is first smoothed via a Savitzky-Golay filter (window length = 21, polynomial order = 2) and fitted with the two-state thermal denaturation model:

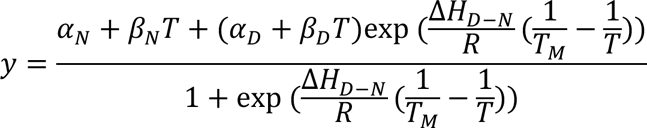

With *α_N_*, *β_N_* and *α_D_, β_D_* the intercept and slope of the linear baselines of respectively the native and denatured states, *R* the gas constant, Δ*H_D-N_* the enthalpy of equilibrium between the native and the denatured state, and *T_M_* the apparent melting temperature. Each 350nm/330nm fluorescence ratio trace is first smoothed via a Savitzky-Golay filter (window length = 21, polynomial order = 2) and then fitted. The temperature of unfolding onset *T*_;<4’=_ is defined as the temperature needed to unfold 5% of the folded population. By definition, *T*_;<4’=_ is a function of *T_M_* and Δ*H_D-N_*:

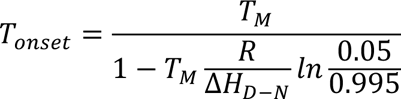

### 5.14 BLI affinity measurements

BLI measurements were performed using an Octet-BLI K2 system (ForteBio). All assays were carried out in PBS supplemented with 0.05% Tween-20 (Sigma) to suppress non-specific interactions with the sensors. All assays were carried out in a black 96-well plate (Greiner 655209), 200 µl per well, and all sensors were subjected to pre-hydration in the assay buffer for at least 15 min before usage. The assay plate was kept at 30°C with a shaking speed of 1000 rpm. The loading wells contained 50 nM of biotinylated β_2_-microglobulin or 30 nM of biotinylated SarS-CoV-2 RBD (purchased from CUSABIO, product code CSB-MP3324GMY1-B). All experiments consisted in a baseline step, a loading step, another baseline step, followed by several association and short dissociation steps. After the last associations step, a long dissociation step is performed. The number of association/dissociation steps, their time, and analyte concentrations employed varied among experiments (see **Fig. 5** and **Supplementary** Fig. 17 and their captions). In all experiments a reference sensor (loaded in the same way as the assay sensors but probing only buffer wells in all association steps) was employed and its signal was subtracted from that of each assay sensor before data analysis. Binding data of all Nb24 nanobody variants were fitted globally with a 1:1 partial dissociation binding model using R_max_, on rate, and off rate as global parameters and Y_t→inf_ as local parameter. Data of all mNb6 variants were fitted globally with a standard 1:1 binding model using R_max_, on rate, and off rate as global parameters.

## 6 Conflict of Interest

The authors declare no conflict of interest.

## 7 Author Contributions

PS conceived and supervised the project. AR developed the deep learning architecture with the guidance of MG and PS. AR parsed and collected the training data with the help of AS and KD. AR built the humanisation pipeline and designed the humanised variants. M. Ali, M. Atkinson and XX produced the nanobody variants and carried out wet-lab experiments. CV and SR produced β_2_-microglobulin and provided expert advice. AR and PS wrote the first version of the paper. All authors analysed data and edited the paper.

## 8 Funding

P.S. is a Royal Society University Research Fellow (URF\R1\201461). We acknowledge funding from UKRI EPSRC (EP/X024733/1 ERC starting grant to PS underwritten by UKRI) and from an Isaac Newton Trust/Wellcome Trust ISSF/University of Cambridge Joint Research Grant (MBAG/624 RG89305).

## 9 Data Availability Statement

All data needed to evaluate the conclusions in this article, or that are necessary to interpret, verify and extend the research in the article are available online in the AbNatiV GitLab at https://gitlab.developers.cam.ac.uk/ch/sormanni/abnativ or in the Methods Section and Supplementary Materials. Additional details are available from the corresponding author on request.

## 10 Code Availability Statement

The AbNatiV code repository including the trained models and the automated humanisation pipeline is available at https://gitlab.developers.cam.ac.uk/ch/sormanni/abnativ. A user-friendly webserver to run AbNatiV is provided at www-cohsoftware.ch.cam.ac.uk/index.php/abnativ. To access the webserver, users need to register a free account and log in.

## 14 Extended Data

**Extended Data Table 1.** Evaluation of the PR classification and reconstruction tasks for human Vκ light-chain sequences. The assessment is carried out for AbNatiV trained on human Vκ sequences (first row) and other computational approaches that can assess humanness (other rows). The first six columns report the PR-AUC (curves shown in **Extended Data Fig. 1**B-C and **Supplementary** Fig. 9A-D), assessing the ability of the models to separate sequences in the Human Test (T) or the Human Diverse >2.5% (D) sets from those from mouse, rhesus, and PSSM-generated (see column headers). The last two columns quantify the ability of each model to reconstruct human sequences in each dataset (column header). The OASis method does not carry out reconstruction. Many sequences of the D datasets belong to the Sapiens training set. See ROC results in **Supplementary** Table 2.

| Vκ | Classification (PR-AUC) | | | | | | Reconstruction accuracy | |
| --- | --- | --- | --- | --- | --- | --- | --- | --- |
|  | Rhesus vs |  | Mouse vs |  | PSSM-generated vs |  | Human Test (T) | Human Diverse >2.5% (D) |
|  | T | D | T | D | T | D |  |  |
| AbNatiV | 0.809 | 0.769 | 0.998 | 0.997 | 0.992 | 0.988 | 0.982 | 0.979 |
| OASis (relaxed) | 0.734 | 0.744 | 0.993 | 0.993 | 0.959 | 0.961 | N/A | N/A |
| Sapiens | 0.848 | 0.860 | 0.993 | 0.993 | 0.989 | 0.989 | 0.935 | 0.939 |

**Extended Data Table 2.**
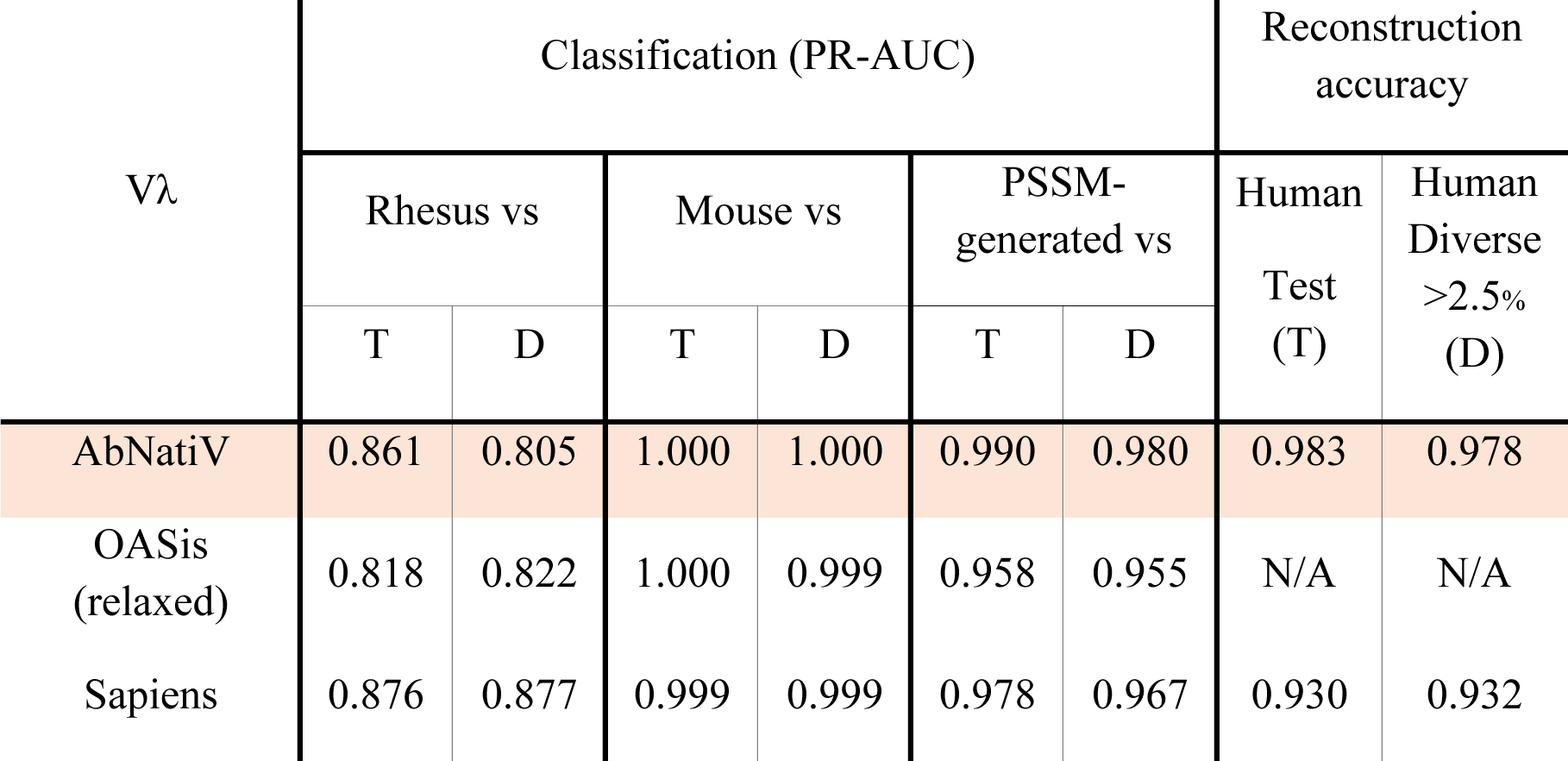
Evaluation of the PR classification and reconstruction tasks for human Vλ light-chain sequences. The assessment is carried out for AbNatiV trained on human Vλ sequences (first row) and other computational approaches that can assess humanness (other rows). The first six columns report the PR-AUC (curves shown in **Extended Data Fig. 1**D-E and **Supplementary** Fig. 9E-H), assessing the ability of the models to separate sequences in the Human Test (T) or the Human Diverse >2.5% (D) sets from those from mouse, rhesus, and PSSM-generated (see column headers). The last two columns quantify the ability of each model to reconstruct human sequences in each dataset (column header). The OASis method does not carry out reconstruction. Many sequences of the D datasets belong to the Sapiens training set. See ROC results in **Supplementary** Table 3.

**Extended Data Table 3.**
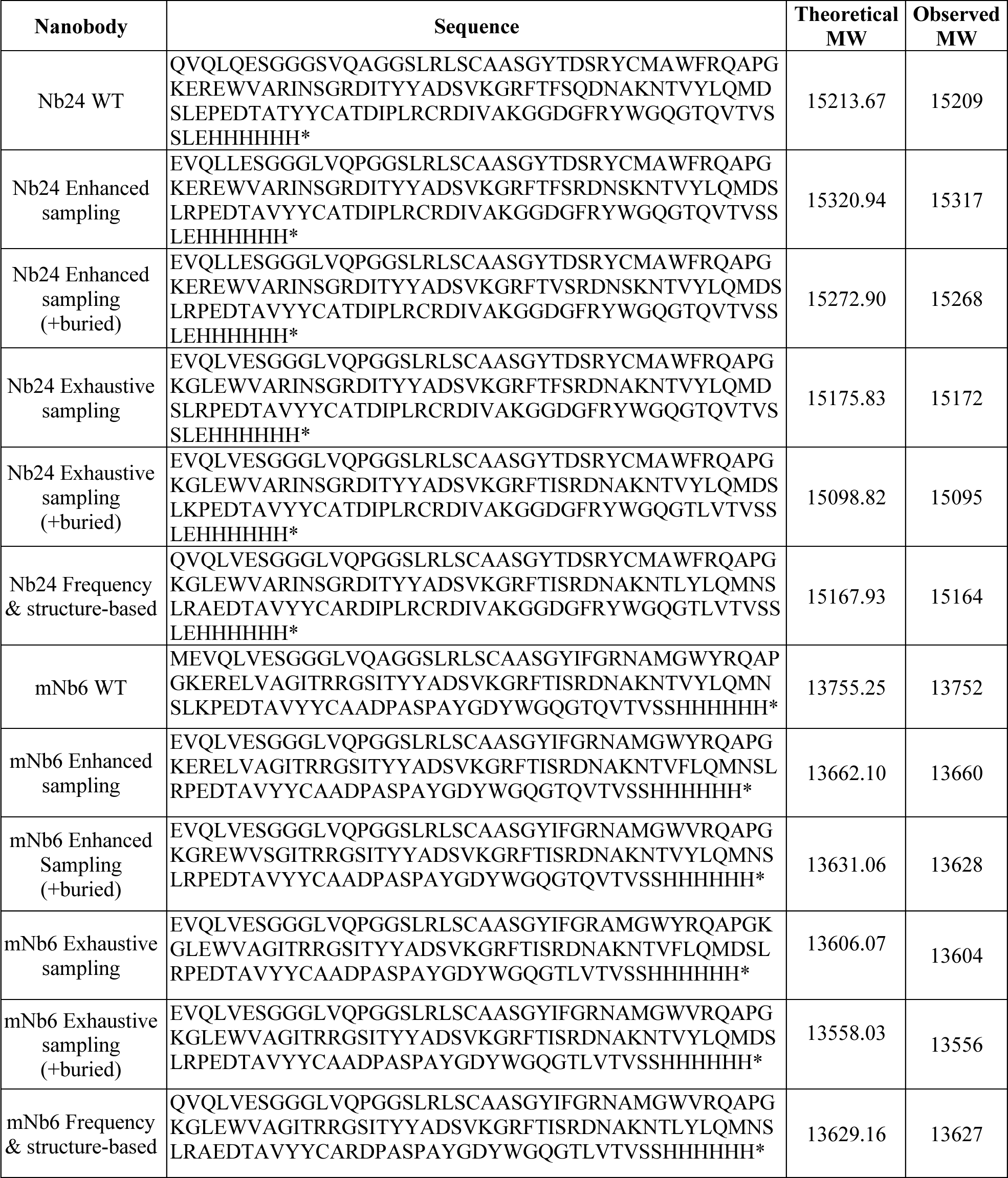
Nanobody sequences experimentally tested. Sequences of the WT nanobodies Nb24 and mNb6 and their humanised variants as used in the wet-lab experiments. A PelB signal sequence was present at the N-terminus of all nanobodies, but this is cleaved upon secretion and hence it is not part of the final protein. All humanised designs are done with AbNatiV except for the Frequency & structure-based designs, which are done with the Llamanade software (45). The theoretical MW is calculated from the amino acid sequence assuming reduced disulphide bonds, observed MW is measured with LC-MS. Nb24 variants have two disulphide bonds and mNb6 have one. Therefore, a difference of -4 Da and -2 Da respectively for Nb24 and mNb6 variants is expected between theoretical and observed MWs.

**Extended Data Fig. 1.**
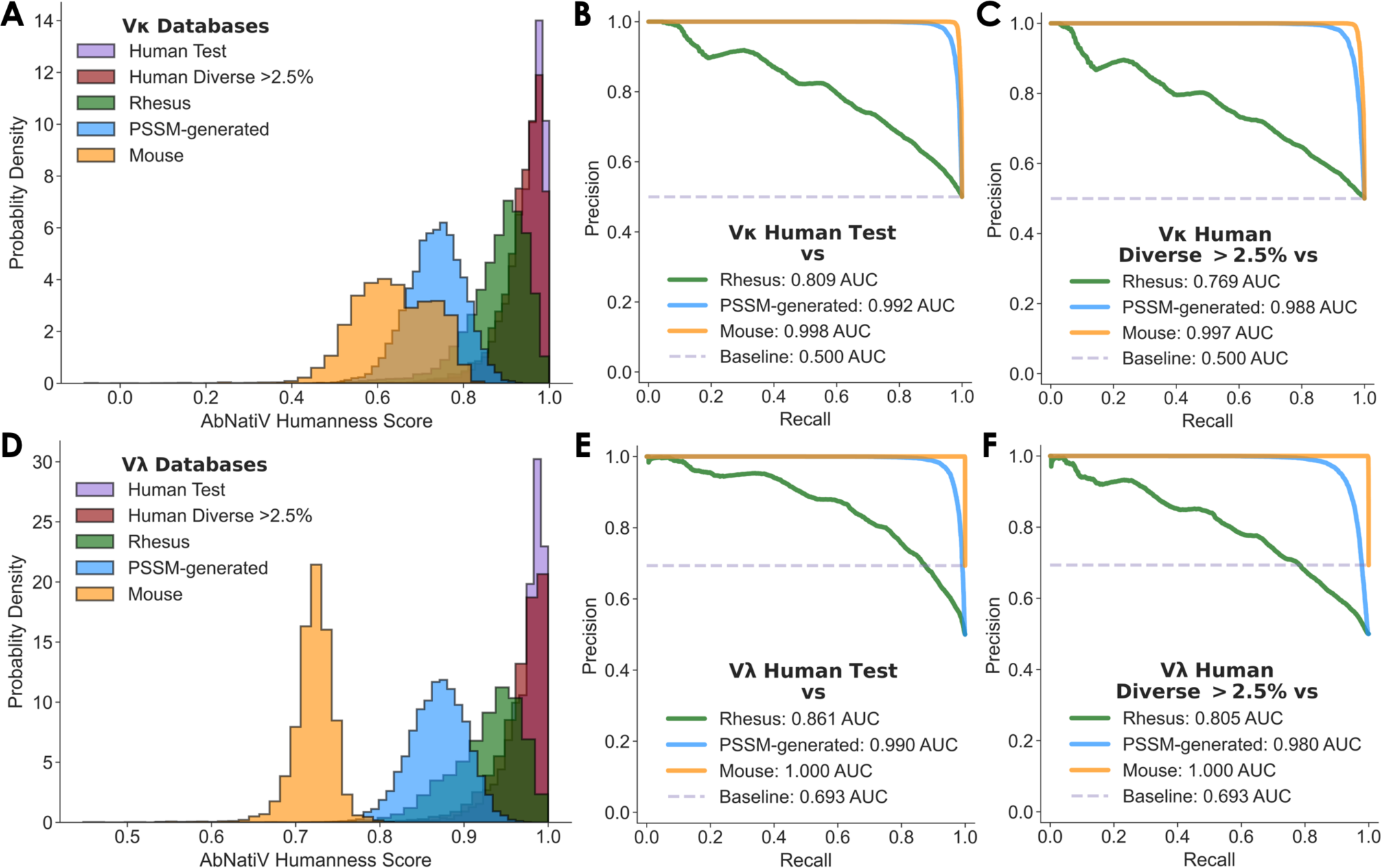
Performance on Vκ and Vλ sequence classification. (**A, D**) The AbNatiV humanness score distributions of the Human Test (purple), Human Diverse >2.5% (red), Rhesus (green), PSSM-generated (blue), and Mouse (orange) Vκ (**A**) and Vλ (**D**) antibody datasets. The PSSM-generated database is made of artificial sequences randomly generated using residue positional frequencies from the PSSM of the Human Test dataset. The Human Diverse >2.5% dataset is made of sequences from the Test and BioPhi datasets with a sequence identity difference of 2.5% from their respective closest sequence of the corresponding Training set (see Methods). Each dataset contains 10,000 sequences except Human Diverse >2.5% which contains 10,490 sequences for Vκ, and 10,459 for Vλ. (**B, C**) Plots of the PR curves computed to represent the ability of AbNatiV to distinguish the Vκ Human Test set (**B**) or Human Diverse >2.5% (**C**) from the other datasets (see legend, which also reports the area under the curve). (**E, F**) Same PR plots but for the Vλ model. The corresponding ROC curves are given in **Supplementary** Fig. 6C-F. The baseline (dashed line) corresponds to the performance that a random classifier would have with the Mouse dataset.

**Extended Data Fig. 2.**
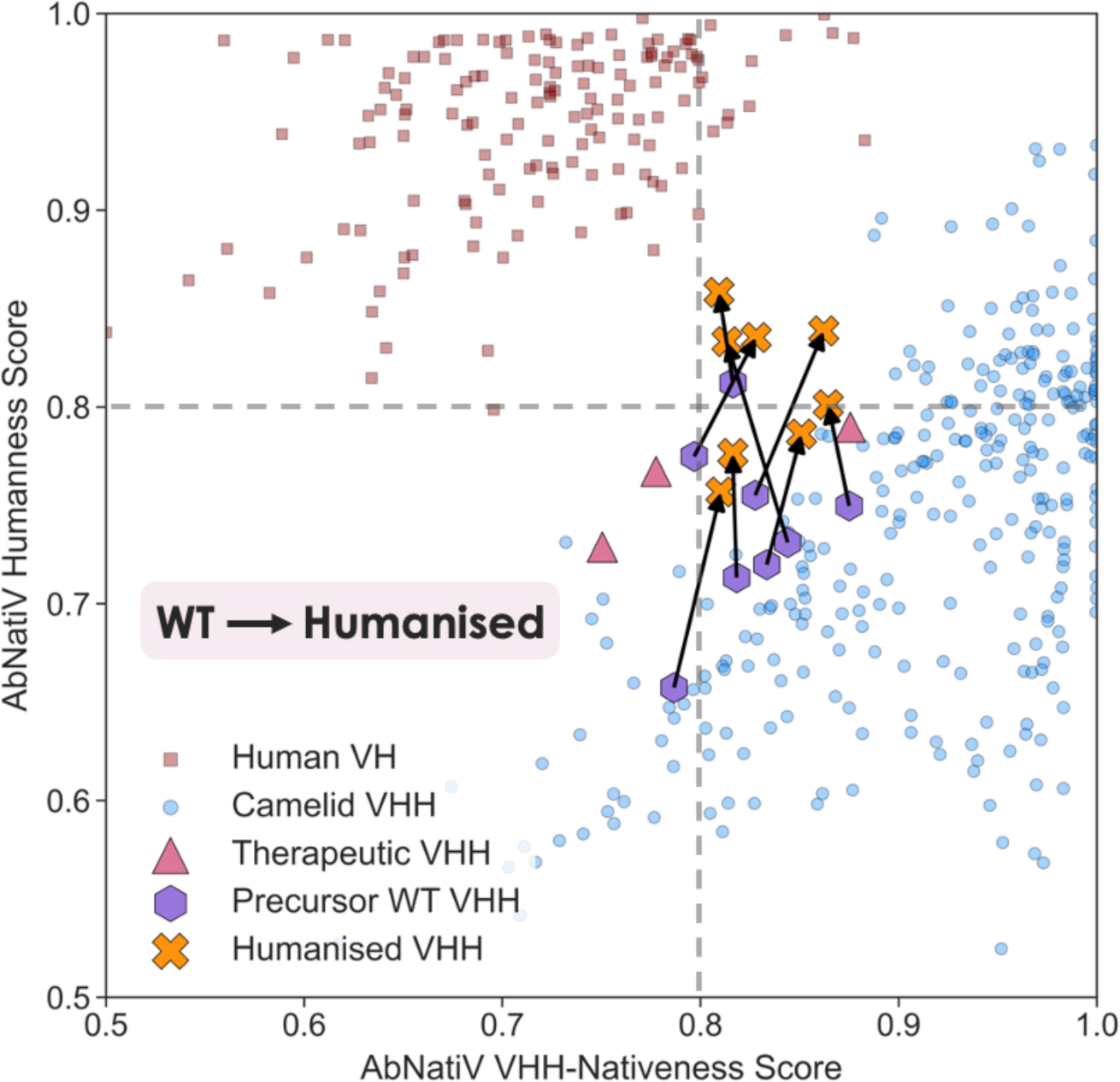
Combining AbNatiV humanness and VHH-nativeness. Plot of the AbNatiV humanness and VHH-nativeness scores of 300 sequences from the VH Human Test (in red), and VHH Camelid Test (in blue) datasets, along with the 3 nanobody therapeutics Envafolimab, Caplacizumab, and Rimteravimab (in pink), and 8 WT nanobodies (in purple) with their humanised counterpart (in orange) (45). An arrow is directed from the WT sequence to the humanised one. Two dashed lines at 0.8 represent the threshold that best separates native from non-native sequences as defined in Methods. Only sequences with a score in [0.5,1] are represented to improve readability. To provide a reference background distribution, 300 randomly selected human VH sequences and 300 camelid VHH sequences are also plotted. The cluster of human sequences that score relatively well in VHH-nativeness derive from the IGHV-3 germline gene (**Supplementary** Fig. 23), consistent with the genetic origin of natural camelid nanobodies (48).

## Notes

### Competing Interest Statement

The authors have declared no competing interest.

### Summary of Updates

Several updates including new humanisation pipelines and wet-lab experiments to validate algorithm

