## Supplementary figures for "AbNatiV: VQ-VAE-based assessment of antibody and nanobody nativeness for hit selection, humanisation, and engineering"

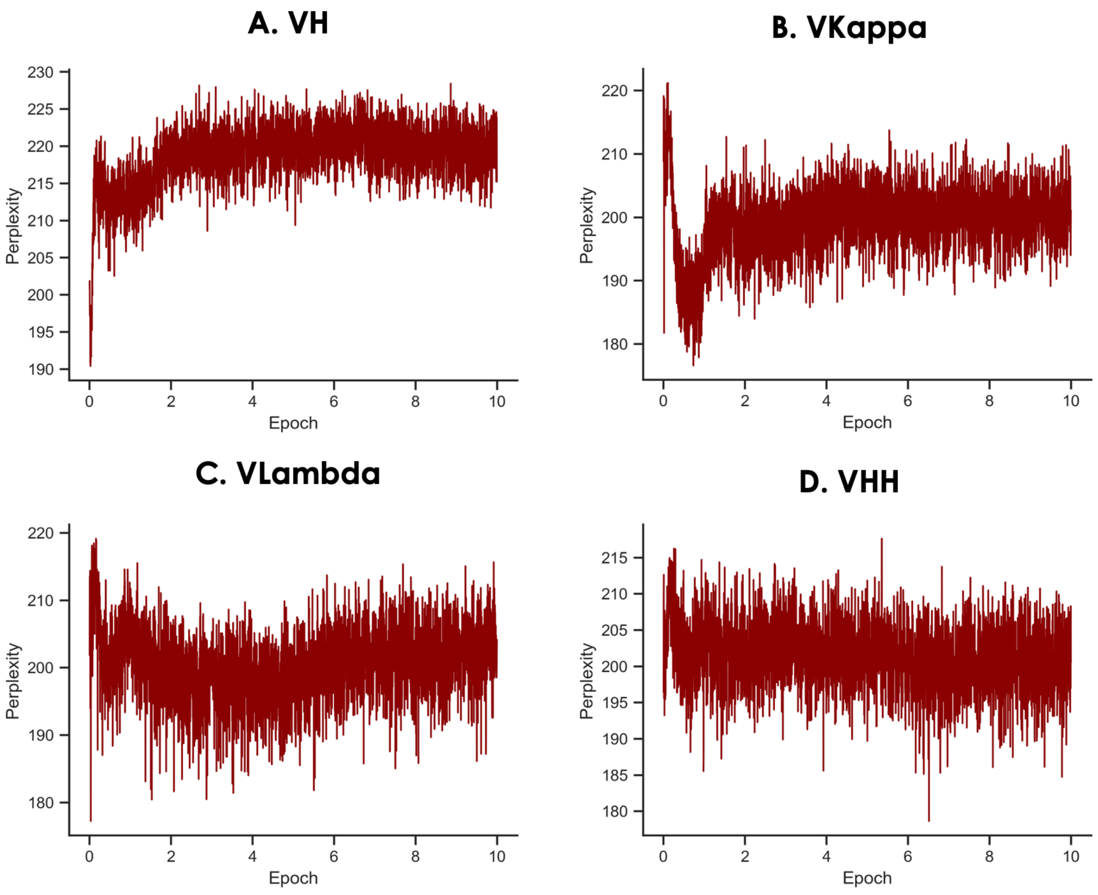


**Supplementary Fig. 1. Perplexity during the training of the AbNatiV models.** (**A**) Perplexity as a function of training epoch of AbNatiV on human VH sequences; (**B**) on human VKappa sequences; (**C**) on human VLambda sequences; (**D**) on camelid VHH sequences.

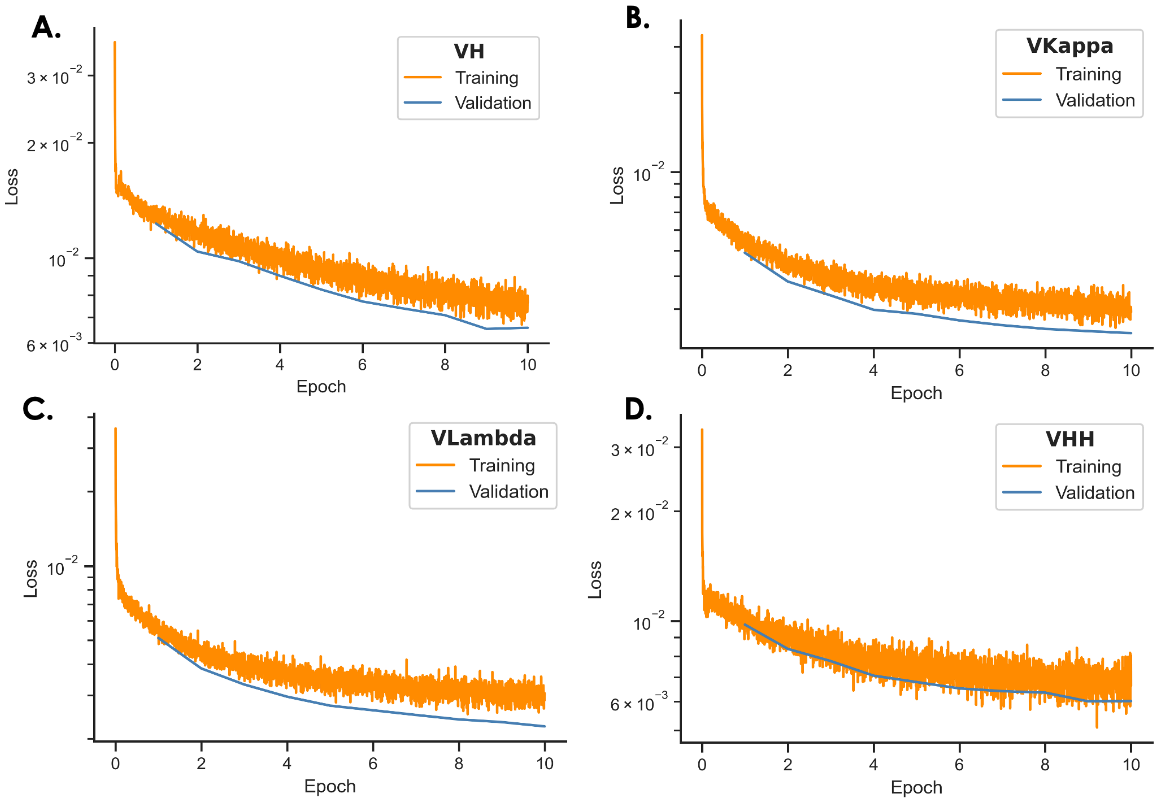


**Supplementary Fig. 2. AbNatiV training performance**. (**A**) Loss function as a function of training epoch evaluated for each training batch (in orange) and averaged on 50,000 validation sequences at each epoch (in blue). Plots are reported for AbNatiV trained on human VH sequences. (**B**) On human VKappa sequences. (**C**) On human VLambda sequences. (**D**) On camelid VHH sequences.

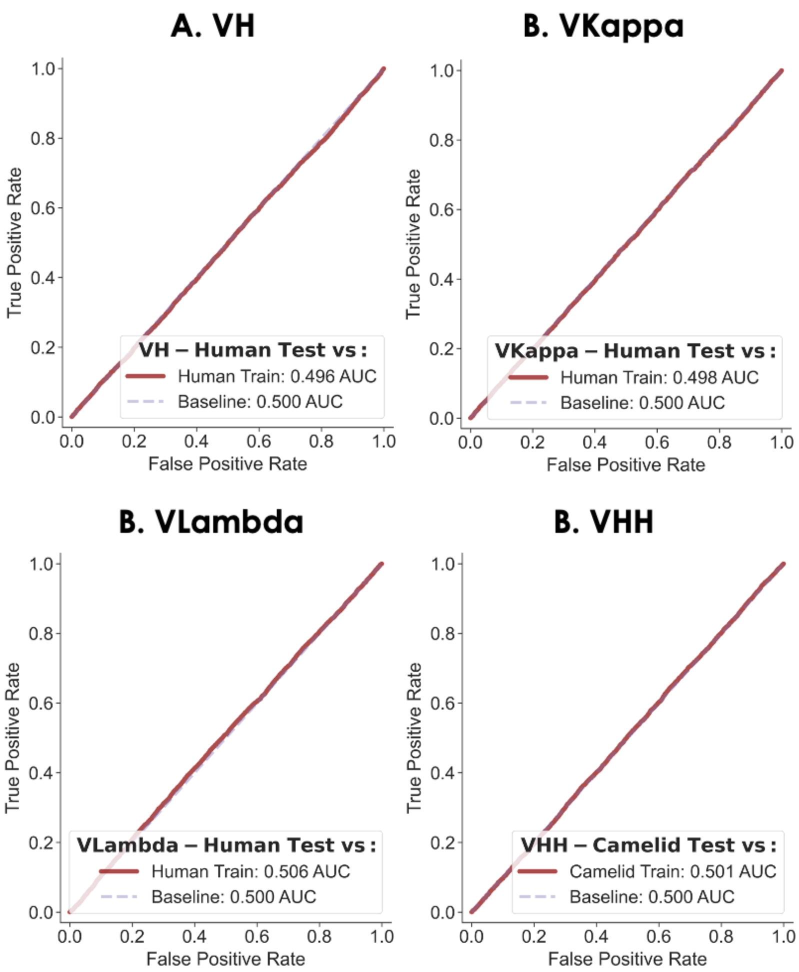


**Supplementary Fig. 3. Overfitting assessment of the AbNatiV models**. (**A**) ROC curves of the ability of AbNatiV to differentiate the score distribution of the Train dataset (random subset of 10 thousand sequences only) and the Test dataset for the AbNatiV models trained on human VH sequences. The analysis is also carried out (**B**) on human VKappa sequences, (**C**) on human VLambda sequences, and (**D**) on camelid VHH sequences. An AUC of 0.50 corresponds to perfectly overlapping distributions.

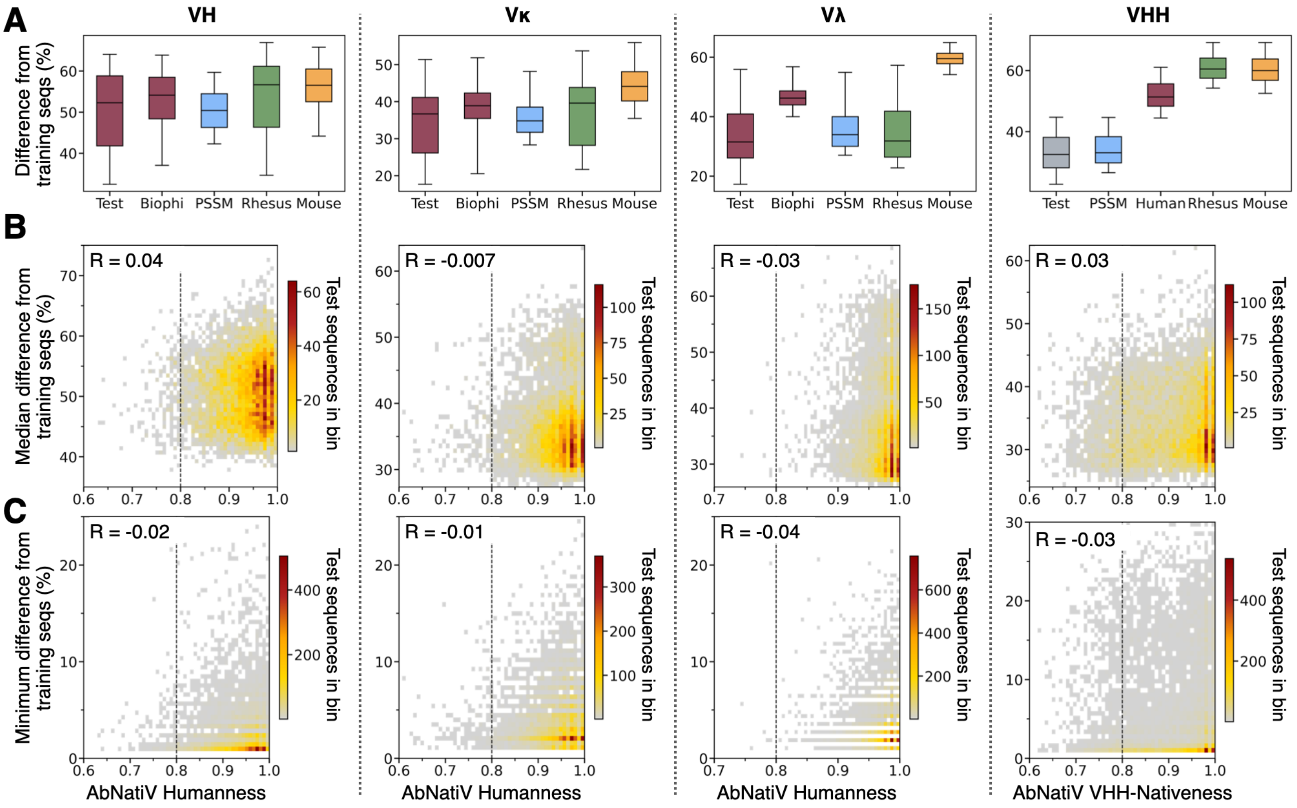


**Supplementary Fig. 4. Analysis of sequence difference across AbNatiV datasets.** The figure is divided in four columns corresponding to each of the four AbNatiV models (see title) introduced in this work. (**A**) The percent sequence difference (y-axis, see Methods) is calculated between all sequences in the training set of each model, and all sequences in each dataset used for the performance assessment of AbNatiV (x-axis). The Boxplots represent the distributions of the values of the percent sequence difference from the training sequences of each sequence in the dataset under scrutiny (x-axis). As calculations are memory-intensive (for instance the VH test set vs. training set only would be $2x10^{10}$ sequence difference values, meaning that each boxplot would need more than 80 Gb of memory), they are carried out in the following way. First, the distance between all sequences in the Training set and each sequence in the Test set (or in any other of the datasets on the x-axis) is calculated. This corresponds to 2 million percent differences (more for VHHs, see **Supplementary Table 5**). Then, only the values of the minimum and of the $5^{th}$, $25^{th}$, $50^{th}$ (i.e., median), $75^{th}$, and $95^{th}$ percentiles of these differences are saved. At the end of the calculation, these values are available for each sequence in the Test set, and the same goes for the other datasets examined (x-axis). In the boxplots, whiskers extend from the median of the values of the $5^{th}$ percentile to that of the values of the $95^{th}$ percentile, boxes from the median of the $25^{th}$s to that of the $75^{th}$s (i.e., the interquartile range), and the horizontal bar is the median of the medians. (**B**) Binned scatter plots of the median percent sequence difference of each sequence in the Test set from all sequences in the training set, plotted as a function of its AbNatiV humanness or VHH-nativeness score. The colour-bar represents the point density of each bin. The dashed vertical line is the AbNatiV score cut-off of 0.8. The Pearson's coefficient of correlation (R) is reported in each panel. (**C**) Same as **B** but reporting the minimum percent difference (that is the percent difference to the closest sequence in the training set) instead of the median.

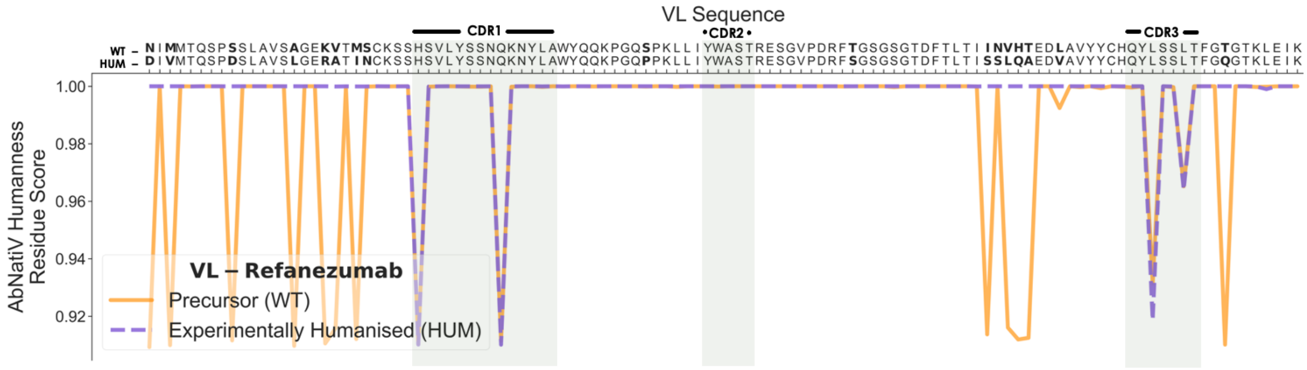


**Supplementary Fig. 5. AbNatiV nativeness profile of the Refanezumab light chain.** AbNatiV humanness residue profiles of the VL mouse precursor and of the humanised sequence of the Refanezumab therapeutic antibody.

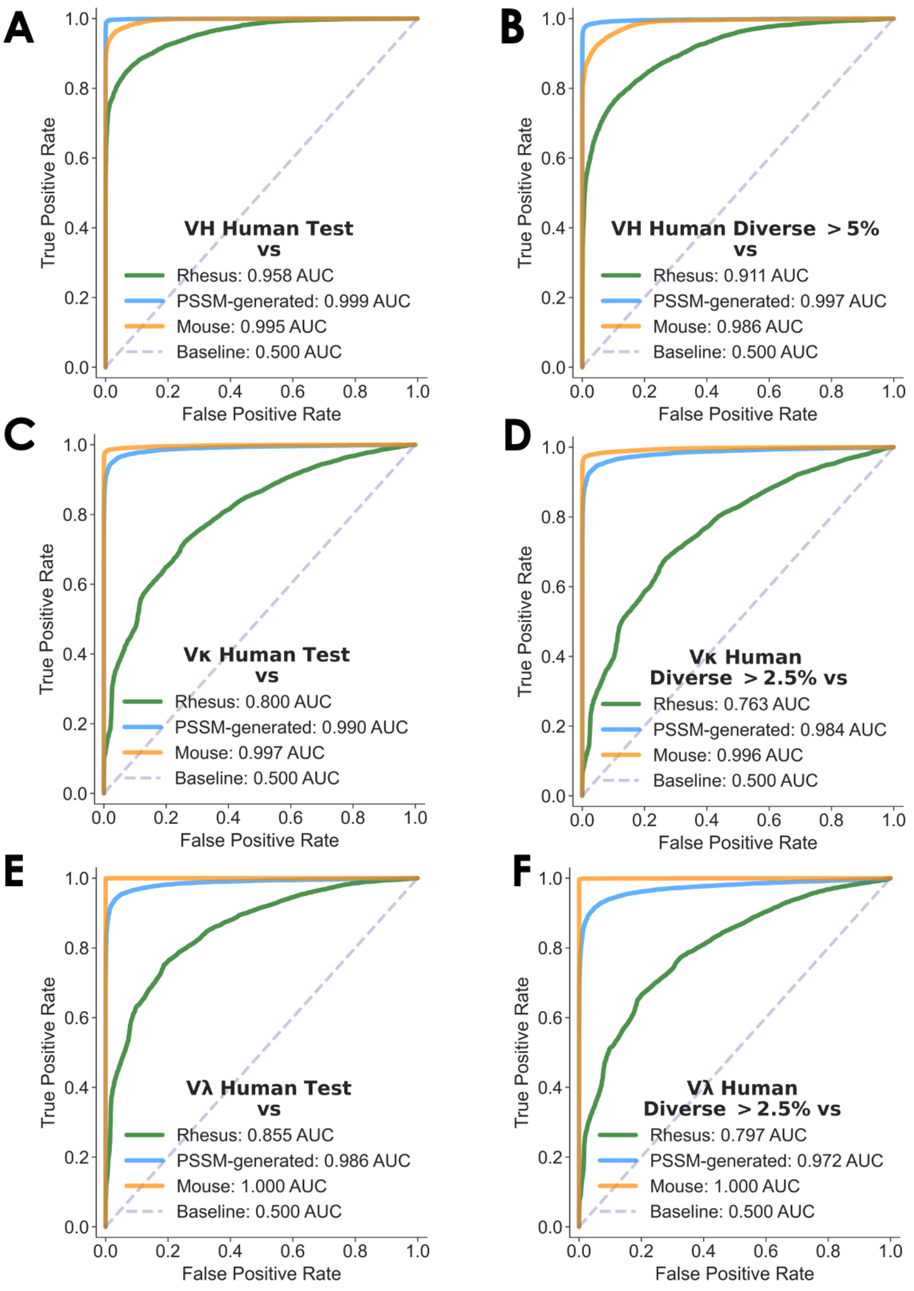


**Supplementary Fig. 6. ROC curves of AbNatiV classifications.** Plots of the ROC curves computed to represent the ability of AbNatiV to distinguish the Human Test sets (**A, C, E**) or Human Diverse sets (**B, D, F**) from the other datasets (see legend, which also reports the area under the curve) for respectively the VH (**A, B**), Vκ (**C, D**), and Vλ (**E, F**) models. The baseline (dashed line) corresponds to the performance of a random classifier.

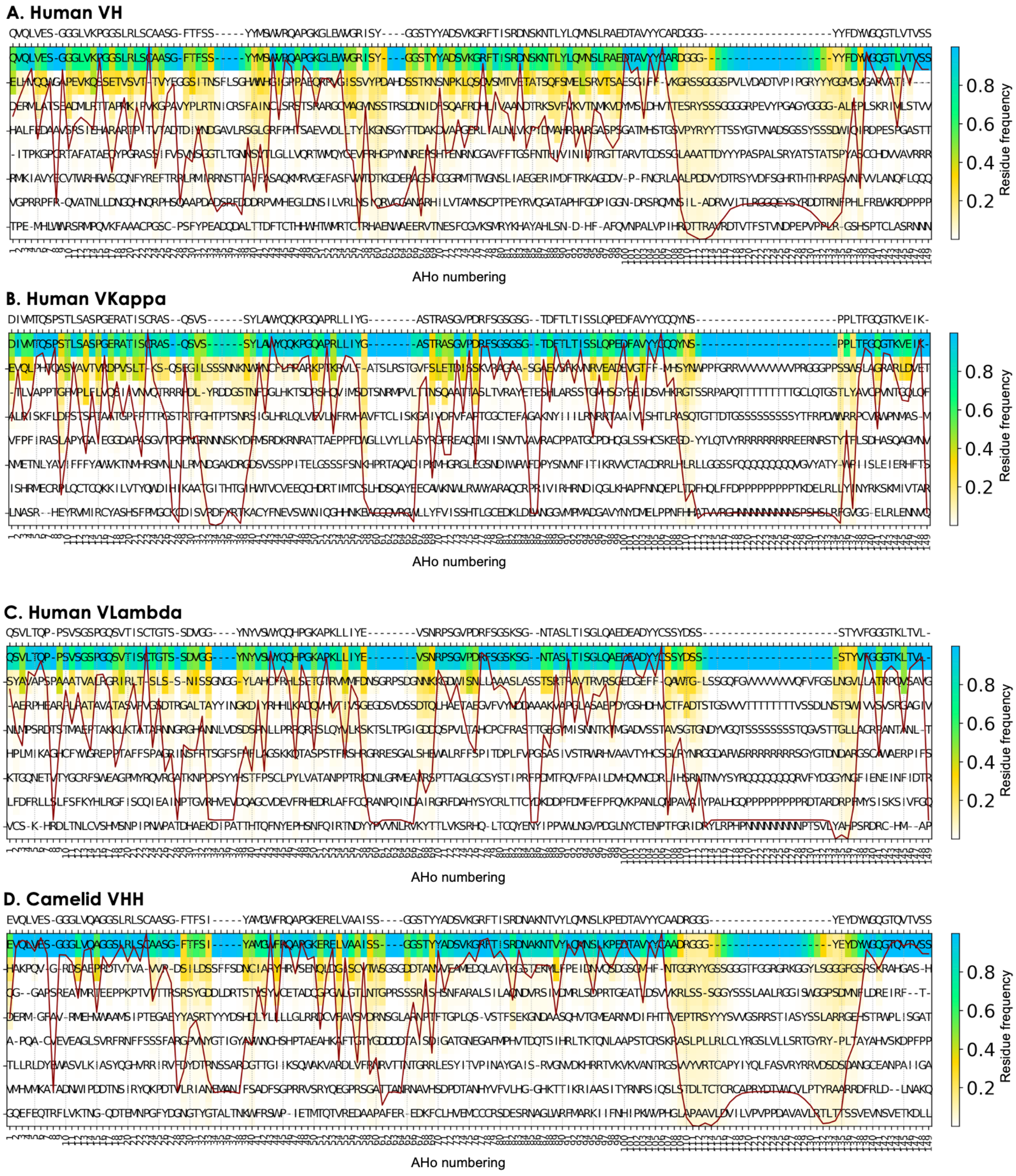


**Supplementary Fig. 7. Position-specific weight matrices (PWMs)**. PWMs are calculated from the training datasets of the human VH (**A**), human VKappa (**B**), human VLambda (**C**), and camelid VHH (**D**) models. The x-axis reports the AHo numbering scheme used for the alignment, the colour-bar is the amino acid frequency at each position. Each column is sorted from most to least frequent residue at that position, and only the top eight residues are shown. Residues are coloured based on their frequency at each position (see colour-bar). The consensus sequence, which consists in the most frequent residue at each position, is shown on the top. The continuous red line is the conservation index of each position (high means position highly conserved, low poorly conserved).

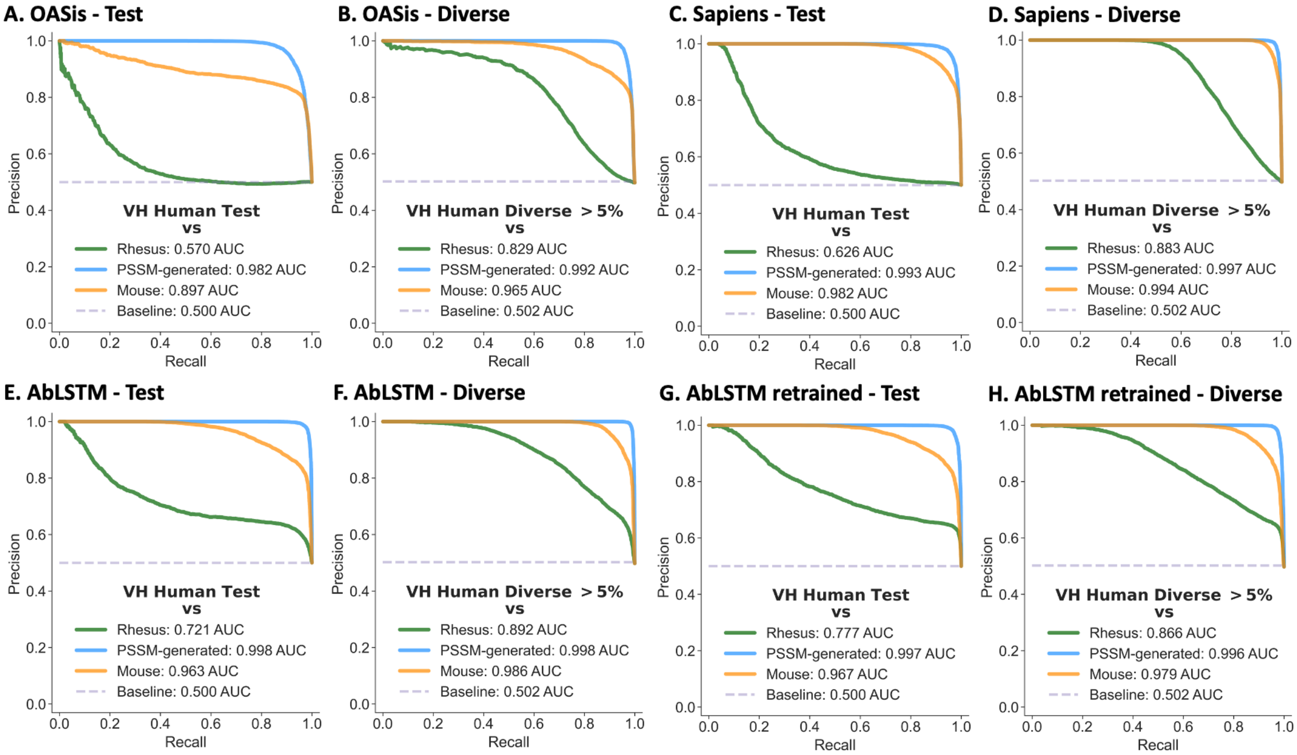


**Supplementary Fig. 8. Performance comparison on VH sequences**. (**A, B**) PR curves to assess the ability of OASis to distinguish the VH Human Test (**A**) or Human Diverse >5% (**B**) dataset from Rhesus (in green), PSSM-generated (in blue), and Mouse (in orange) VH antibody sequences. **(C, D)** ROC curves for Sapiens. (**E, F**) ROC curves for AbLSTM. (**G, H**) ROC curves for AbLSTM retrained on the AbNatiV training set. The baseline (dashed line) corresponds to the performance that a random classifier would have.

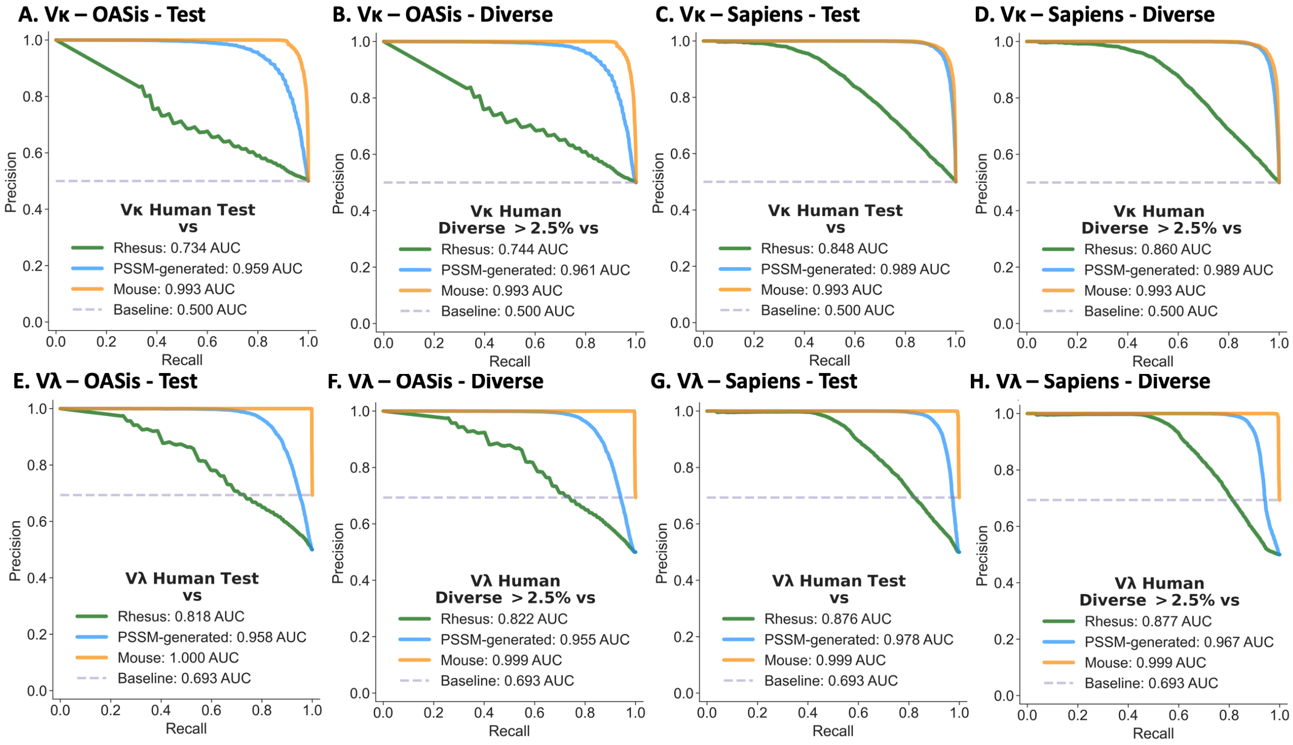


**Supplementary Fig. 9. Performance comparison on Vκ and Vλ sequences classification**. (**A, B, C, D**) PR curves to assess the ability of OASis (**A, B**) and Sapiens (**C, D**) to distinguish the Vκ Human Test (**A** and **C**) or Diverse >2.5% (**B** and **D**) dataset from Rhesus (in green), PSSM-generated (in blue), and Mouse (in orange) Vκ light chain antibody sequences. (**E, F, G, H**) Corresponding panels for the Vλ light chain sequences and respective datasets. The baseline (dashed line) corresponds to the performance that a random classifier would have with the Mouse dataset.

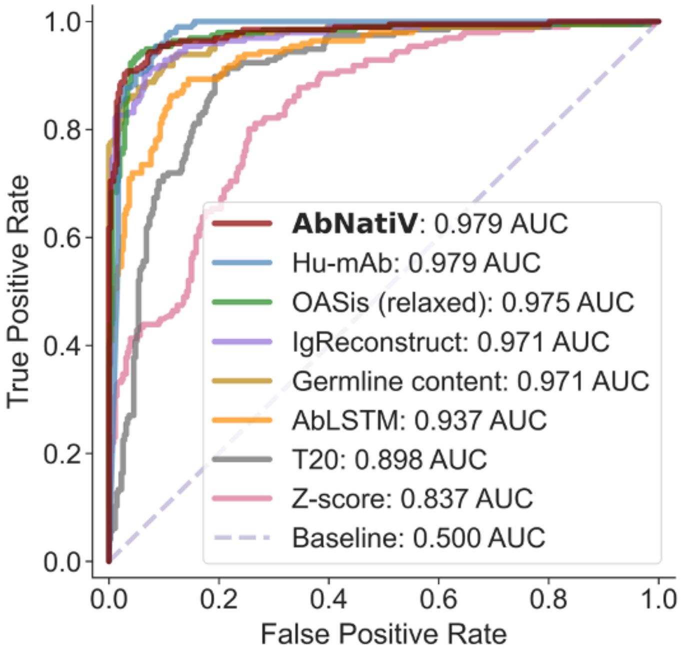


**Supplementary Fig. 10. Classification performance on antibody therapeutics.** Plot of the ROC curves of the classification of 196 human therapeutics from 353 non-human ones (from mouse, chimeric, and humanised) carried out by AbNatiV (in red) and seven other computational methods (see legend, which also reports the area under the curve). The baseline (dashed line) corresponds to the performance that a random classifier would have.

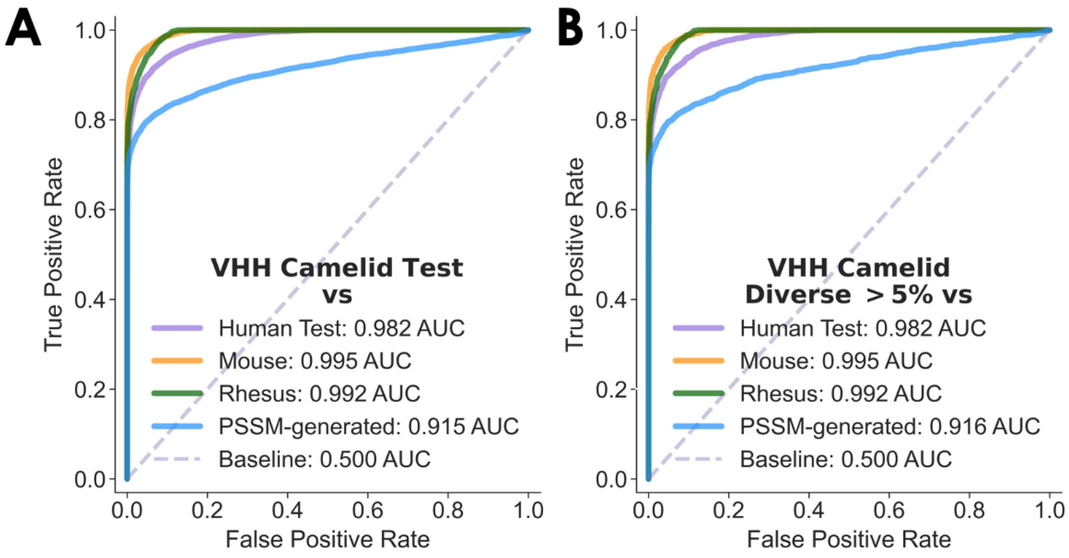


**Supplementary Fig. 11. ROC performance on VHH classification.** Plots of the ROC curves computed to represent the ability of AbNatiV to distinguish the VHH Test set (**B**) or VHH Diverse >5% (**C**) from the other datasets (see legend, which also reports the area under the curve). The baseline (dashed line) corresponds to the performance that a random classifier would have.

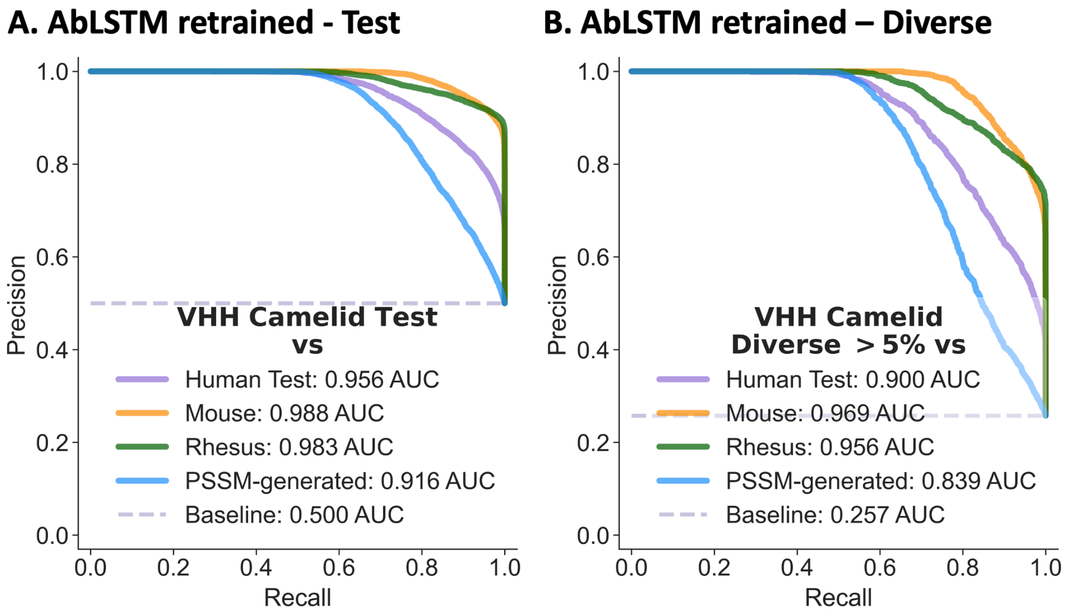


**Supplementary Fig. 12. Performance comparison on VHH sequence classification.** PR curves to assess the ability of AbLSTM retrained on the Camelid dataset to distinguish the VHH Test (**A**) or VHH Diverse >5% dataset (**B**) from the Human Test (in purple), Mouse (in orange) , Rhesus (in green), and PSSM-generated (in blue) VH antibody datasets. The baseline (dashed line) corresponds to the performance that a random classifier would have.

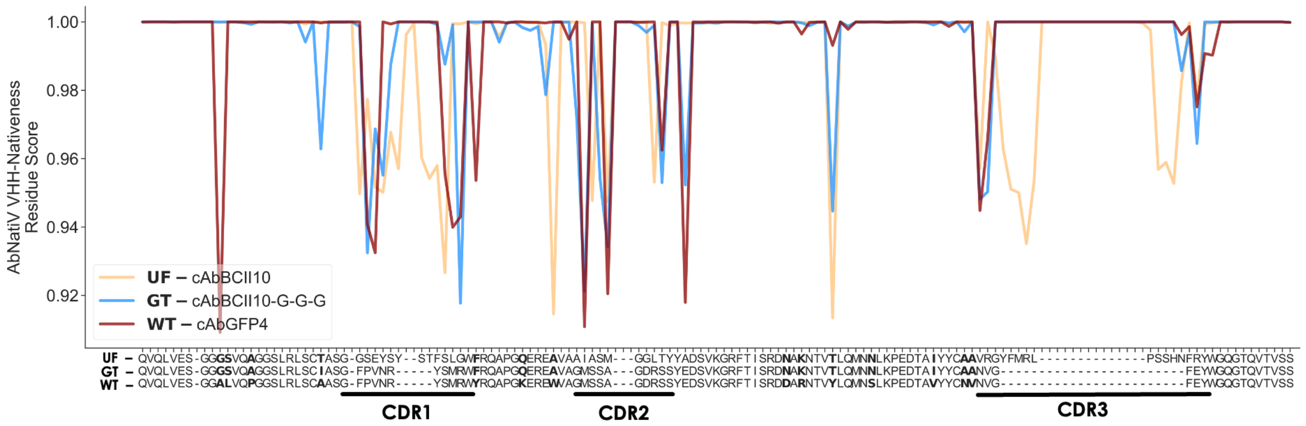


**Supplementary Fig. 13. VHH grafting sensitivity of AbNatiV**. AbNatiV VHH-nativeness profile of the VHH universal framework (UF) cAbCII10 (in orange), superimposed to the AbNatiV VHH-nativeness profile of the native cAbGFP4 nanobody (in red) and its grafted version (in blue), where all three CDRs have been grafted onto the UF scaffold (see Methods and main text).

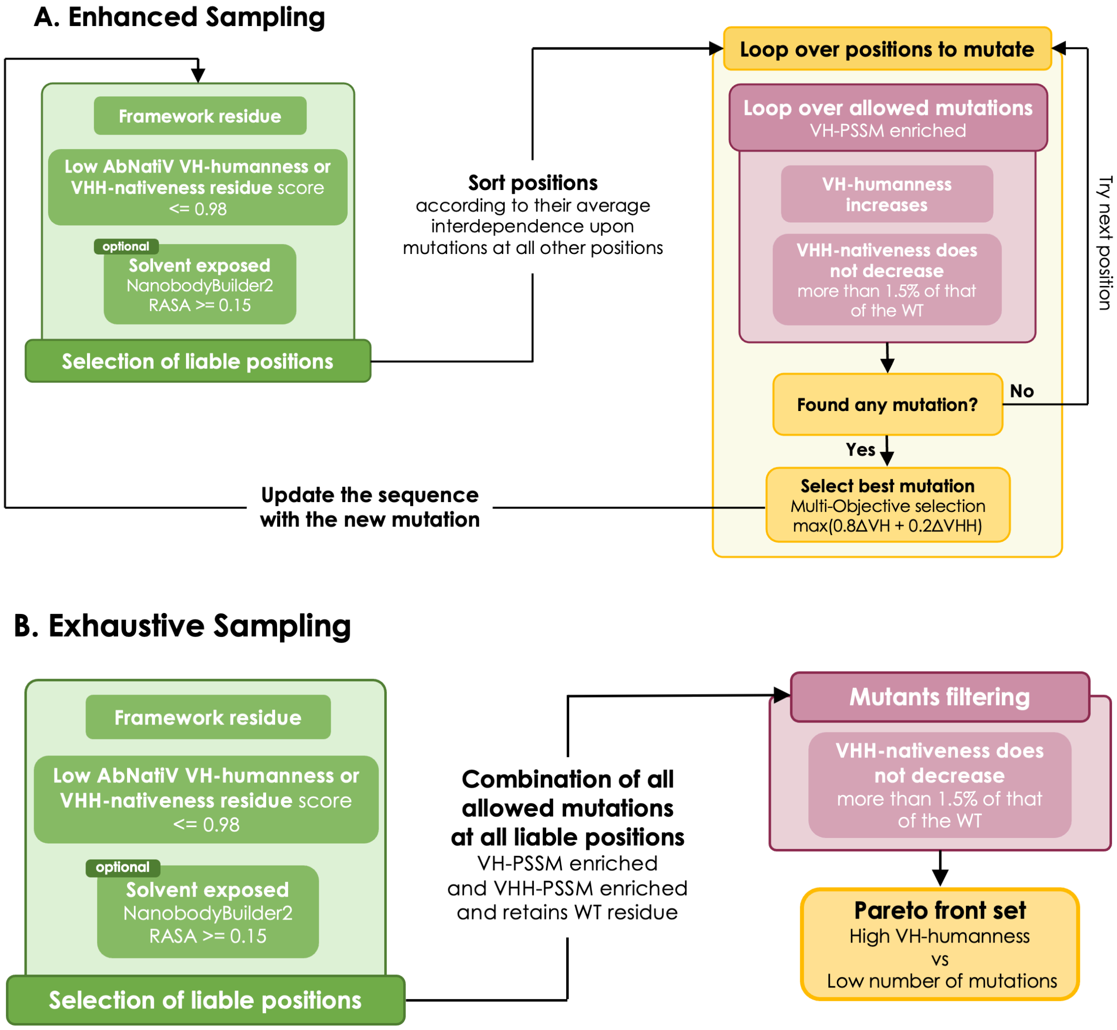


**Supplementary Fig. 14. Illustration of the Enhanced and Exhaustive sampling strategies used in the automated AbNatiV humanisation pipeline.** Both sampling strategy start by selecting the liable positions to mutate (in green). Liable positions are framework residues with a AbNatiV VH-humanness or VHH-nativeness residue score <= 0.98. Optionally, liable positions can be restricted to solvent-exposed only residues with a RASA score >= 0.15 upon NanoBuilder2 modelling of the structure (see Methods). Then the two strategies differ. (**A**) The Enhanced sampling approach iteratively explores the mutational space. The order at which positions are mutated is defined starting from those mutable positions whose score is least affected by mutations at other positions (see Methods). Subsequently, we loop over these sorted liable positions to mutate (in yellow). We mutate a given position with all the amino acids significantly enriched at that AHo position in the human VH PWM (i.e., with a PSSM log-likelihood score > 0 and a PWM frequency > 0.01, see **Supplementary Fig. 7**). Cysteines and methionines are not allowed for mutation. In pink, we keep mutations which increase the VH-humanness score of the sequence (i.e., ΔVH > 0), and which do not decrease the VHH-nativeness score by more than 1.5% (i.e., 1.5% decrease tolerance of ΔVHH). If no such mutations are found, the residue is left to WT, and the procedure continues to the next liable position. If more than one mutation is kept, we select the one that increases most the multi-objective function: $0.8\Delta VH + 0.2\Delta VHH$. The sequence is then updated with the selected mutation and the process of selecting mutable positions for mutation is repeated (in green). (**B**) The exhaustive approach assesses all combination of mutations allowed by the mutational space and selects the best variant. At each position, we only allow amino acids enriched in both human VH and VHH PWMs (i.e., with a PSSM log-likelihood score > 0 and a PWM frequency > 0.01, see **Supplementary Fig. 7**). Cysteines and methionines are excluded, and the WT residue is added to the options if not already present. All possible combinations of such mutations at all liable positions are then scored, and those that do not decrease the VHH-nativeness score by more than 1.5% of that of the WT are kept (in pink). Finally, we compute the Pareto front (in yellow) that maximises the VH-humanness score while minimising the number of mutations over all remaining mutation combinations. This approach returns a set of variants with the highest VH-humanness for each number of mutations that are beneficial to the VH-humanness (see **Supplementary Fig. 21**). Increasing the number of mutations is beneficial when it increases the VH-humanness score. In this work, only sequences sequence exhibiting the highest humanness score were experimentally tested.

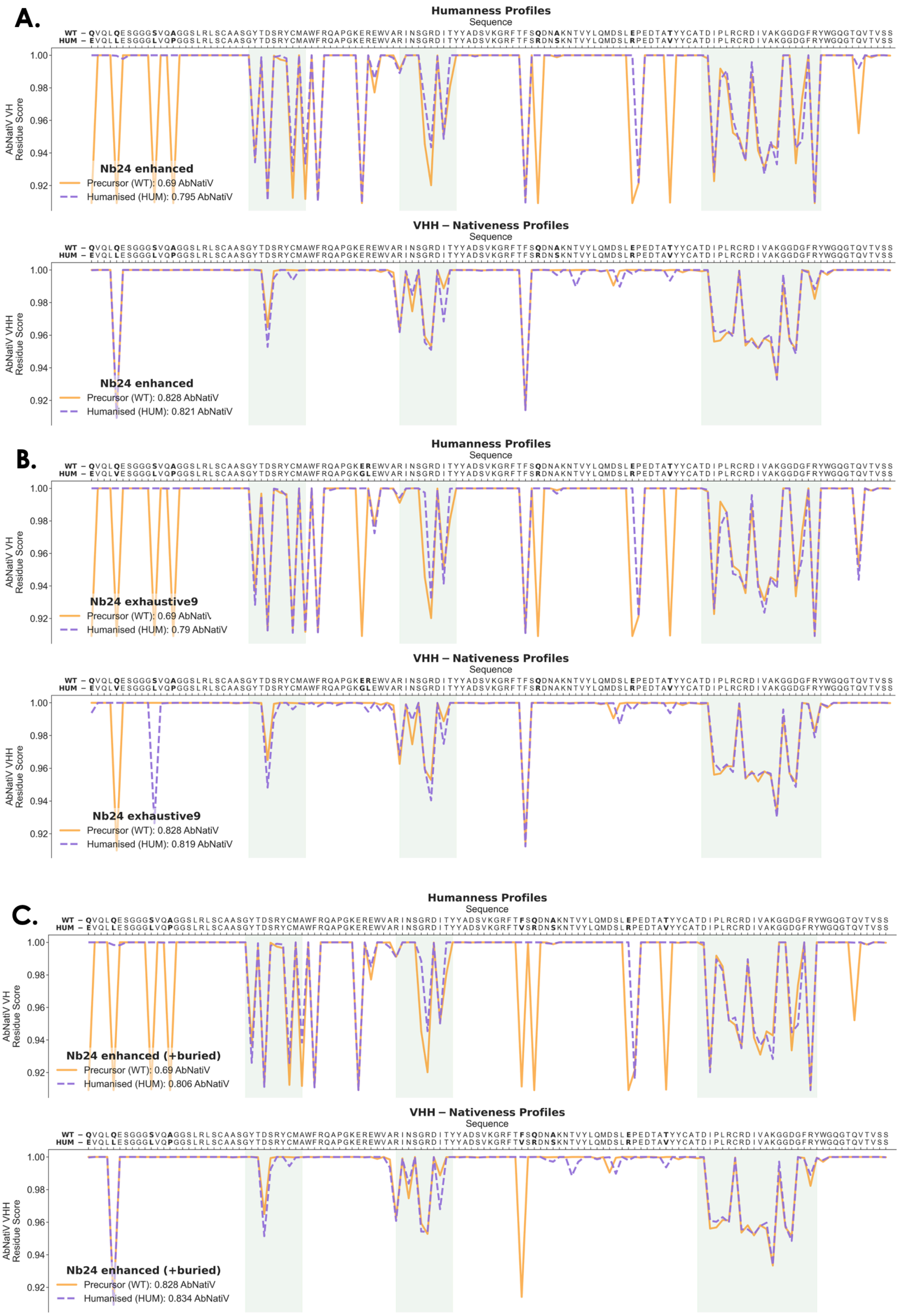
A.
Humanness Profiles
Sequence
WT – QVQLQESGGGSVQAGGSLRLSCAASGYTDSRYCMAWFRQAPGKEREWVARINSGRDITYYADSVKGRFTFSQDNAKNTVYLQMDSLEPEDTATYYCATDIPLRCRDIVAKGGDGFRYWGQGTQVTVSS
HUM – EVQLLESGGGLVQPGGSLRLSCAASGYTDSRYCMAWFRQAPGKEREWVARINSGRDITYYADSVKGRFTFSRDNSKNTVYLQMDSLRPEDTAVYYCATDIPLRCRDIVAKGGDGFRYWGQGTQVTVSS
AbNatiV VH Residue Score
1.00
0.98
0.96
0.94
0.92
Nb24 enhanced
Precursor (WT): 0.69 AbNatiV
Humanised (HUM): 0.795 AbNatiV
VHH – Nativeness Profiles
Sequence
WT – QVQLQESGGGSVQAGGSLRLSCAASGYTDSRYCMAWFRQAPGKEREWVARINSGRDITYYADSVKGRFTFSQDNAKNTVYLQMDSLEPEDTATYYCATDIPLRCRDIVAKGGDGFRYWGQGTQVTVSS
HUM – EVQLLESGGGLVQPGGSLRLSCAASGYTDSRYCMAWFRQAPGKEREWVARINSGRDITYYADSVKGRFTFSRDNSKNTVYLQMDSLRPEDTAVYYCATDIPLRCRDIVAKGGDGFRYWGQGTQVTVSS
AbNatiV VHH Residue Score
1.00
0.98
0.96
0.94
0.92
Nb24 enhanced
Precursor (WT): 0.828 AbNatiV
Humanised (HUM): 0.821 AbNatiV
B.
Humanness Profiles
Sequence
WT – QVQLQESGGGSVQAGGSLRLSCAASGYTDSRYCMAWFRQAPGKEREWVARINSGRDITYYADSVKGRFTFSQDNAKNTVYLQMDSLEPEDTATYYCATDIPLRCRDIVAKGGDGFRYWGQGTQVTVSS
HUM – EVQLVESGGGLVQPGGSLRLSCAASGYTDSRYCMAWFRQAPGKGLEWVARINSGRDITYYADSVKGRFTFSRDNAKNTVYLQMDSLRPEDTAVYYCATDIPLRCRDIVAKGGDGFRYWGQGTQVTVSS
AbNatiV VH Residue Score
1.00
0.98
0.96
0.94
0.92
Nb24 exhaustive9
Precursor (WT): 0.69 AbNati\
Humanised (HUM): 0.79 AbNatiV
VHH – Nativeness Profiles
Sequence
WT – QVQLQESGGGSVQAGGSLRLSCAASGYTDSRYCMAWFRQAPGKEREWVARINSGRDITYYADSVKGRFTFSQDNAKNTVYLQMDSLEPEDTATYYCATDIPLRCRDIVAKGGDGFRYWGQGTQVTVSS
HUM – EVQLVESGGGLVQPGGSLRLSCAASGYTDSRYCMAWFRQAPGKGLEWVARINSGRDITYYADSVKGRFTFSRDNAKNTVYLQMDSLRPEDTAVYYCATDIPLRCRDIVAKGGDGFRYWGQGTQVTVSS
AbNatiV VHH Residue Score
1.00
0.98
0.96
0.94
0.92
Nb24 exhaustive9
Precursor (WT): 0.828 AbNatiV
Humanised (HUM): 0.819 AbNatiV
C.
Humanness Profiles
Sequence
WT – QVQLQESGGGSVQAGGSLRLSCAASGYTDSRYCMAWFRQAPGKEREWVARINSGRDITYYADSVKGRFTFSQDNAKNTVYLQMDSLEPEDTATYYCATDIPLRCRDIVAKGGDGFRYWGQGTQVTVSS
HUM – EVQLLESGGGLVQPGGSLRLSCAASGYTDSRYCMAWFRQAPGKEREWVARINSGRDITYYADSVKGRFTVSRDNSKNTVYLQMDSLRPEDTAVYYCATDIPLRCRDIVAKGGDGFRYWGQGTQVTVSS
AbNatiV VH Residue Score
1.00
0.98
0.96
0.94
0.92
Nb24 enhanced (+buried)
Precursor (WT): 0.69 AbNatiV
Humanised (HUM): 0.806 AbNatiV
VHH – Nativeness Profiles
Sequence
WT – QVQLQESGGGSVQAGGSLRLSCAASGYTDSRYCMAWFRQAPGKEREWVARINSGRDITYYADSVKGRFTFSQDNAKNTVYLQMDSLEPEDTATYYCATDIPLRCRDIVAKGGDGFRYWGQGTQVTVSS
HUM – EVQLLESGGGLVQPGGSLRLSCAASGYTDSRYCMAWFRQAPGKEREWVARINSGRDITYYADSVKGRFTVSRDNSKNTVYLQMDSLRPEDTAVYYCATDIPLRCRDIVAKGGDGFRYWGQGTQVTVSS
AbNatiV VHH Residue Score
1.00
0.98
0.96
0.94
0.92
Nb24 enhanced (+buried)
Precursor (WT): 0.828 AbNatiV
Humanised (HUM): 0.834 AbNatiV

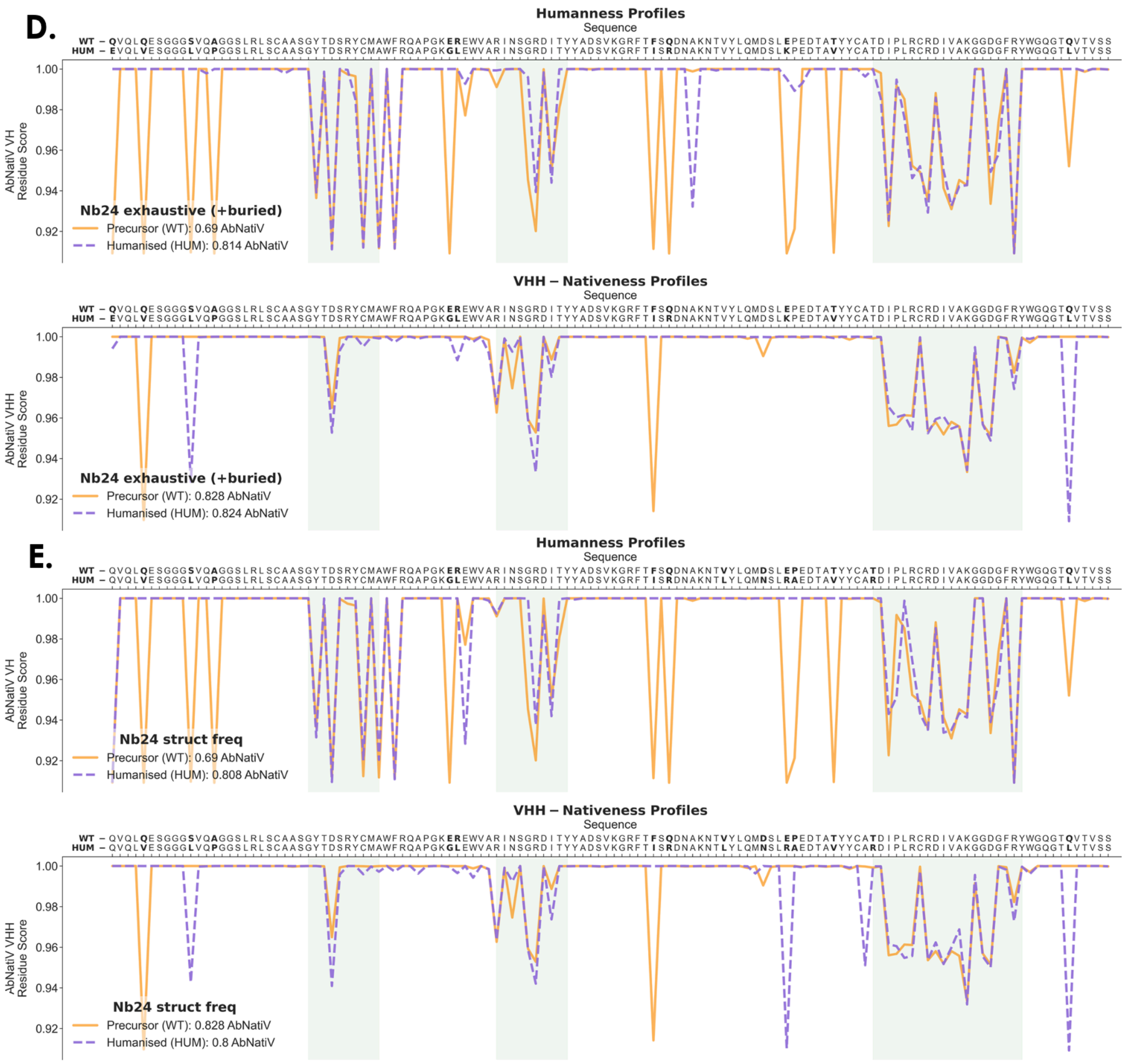


**Supplementary Fig. 15. AbNatiV VH humanness (top) and VHH nativeness (bottom) profiles of the humanised designs of Nb24.** AbNatiV profiles are represented for the WT (in yellow) and humanised counterpart (in purple) of Nb24. (**A**) AbNatiV enhanced sampling. (**B**) AbNatiV exhaustive sampling. (**C**) AbNatiV enhanced sampling allowing mutations at buried framework residues. (**D**) AbNatiV exhaustive sampling allowing mutations at buried framework residues. (**E**) AbNatiV profiles for the structure and frequency-based humanised variants (this variant was obtained from the Llamanade web server at http://35.208.211.136/, see Methods).

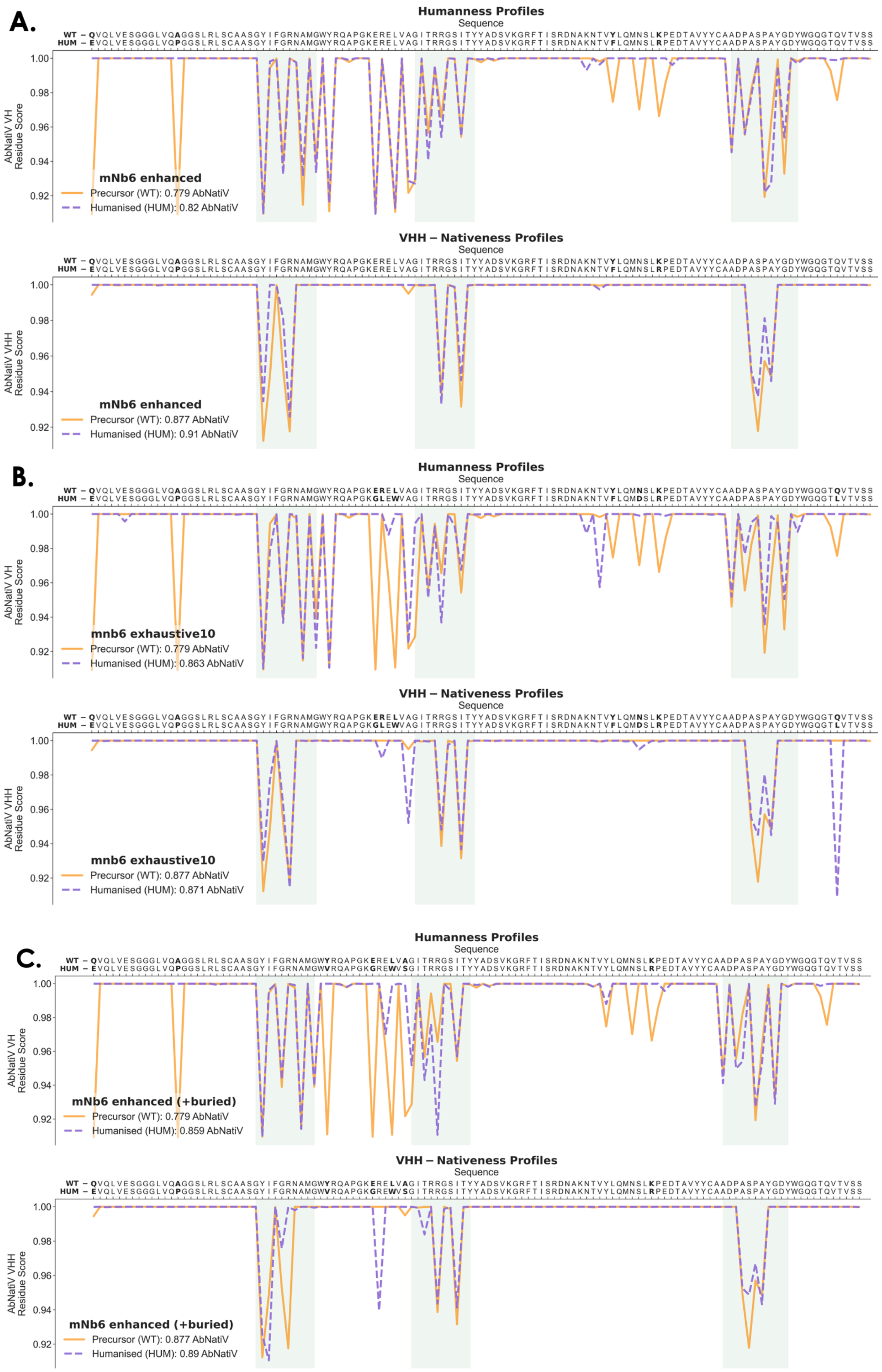
A.
Humanness Profiles
Sequence
WT – QVQLVESGGGLVQAGGSLRLSCAASGYIFGRNAMGWYRQAPGKERELVAGITRRGSITYYADSVKGRFTISRDNAKNTVYLQMNSLKPEDTAVYYCAADPASPAYGDYWGQGTQVTVSS
HUM – EVQLVESGGGLVQPGGSLRLSCAASGYIFGRNAMGWYRQAPGKERELVAGITRRGSITYYADSVKGRFTISRDNAKNTVFLQMNSLRPEDTAVYYCAADPASPAYGDYWGQGTQVTVSS
AbNatiV VH Residue Score
mNb6 enhanced
Precursor (WT): 0.779 AbNatiV
Humanised (HUM): 0.82 AbNatiV
VHH – Nativeness Profiles
AbNatiV VHH Residue Score
mNb6 enhanced
Precursor (WT): 0.877 AbNatiV
Humanised (HUM): 0.91 AbNatiV
B.
Humanness Profiles
WT – QVQLVESGGGLVQAGGSLRLSCAASGYIFGRNAMGWYRQAPGKERELVAGITRRGSITYYADSVKGRFTISRDNAKNTVYLQMNSLKPEDTAVYYCAADPASPAYGDYWGQGTQVTVSS
HUM – EVQLVESGGGLVQPGGSLRLSCAASGYIFGRNAMGWYRQAPGKGLEWVAGITRRGSITYYADSVKGRFTISRDNAKNTVFLQMDSLRPEDTAVYYCAADPASPAYGDYWGQGTLVTVSS
mnb6 exhaustive10
Precursor (WT): 0.779 AbNatiV
Humanised (HUM): 0.863 AbNatiV
VHH – Nativeness Profiles
mnb6 exhaustive10
Precursor (WT): 0.877 AbNatiV
Humanised (HUM): 0.871 AbNatiV
C.
Humanness Profiles
WT – QVQLVESGGGLVQAGGSLRLSCAASGYIFGRNAMGWYRQAPGKERELVAGITRRGSITYYADSVKGRFTISRDNAKNTVYLQMNSLKPEDTAVYYCAADPASPAYGDYWGQGTQVTVSS
HUM – EVQLVESGGGLVQPGGSLRLSCAASGYIFGRNAMGWVRQAPGKGREWVSGITRRGSITYYADSVKGRFTISRDNAKNTVYLQMNSLRPEDTAVYYCAADPASPAYGDYWGQGTQVTVSS
mNb6 enhanced (+buried)
Precursor (WT): 0.779 AbNatiV
Humanised (HUM): 0.859 AbNatiV
VHH – Nativeness Profiles
mNb6 enhanced (+buried)
Precursor (WT): 0.877 AbNatiV
Humanised (HUM): 0.89 AbNatiV
1.00
0.98
0.96
0.94
0.92

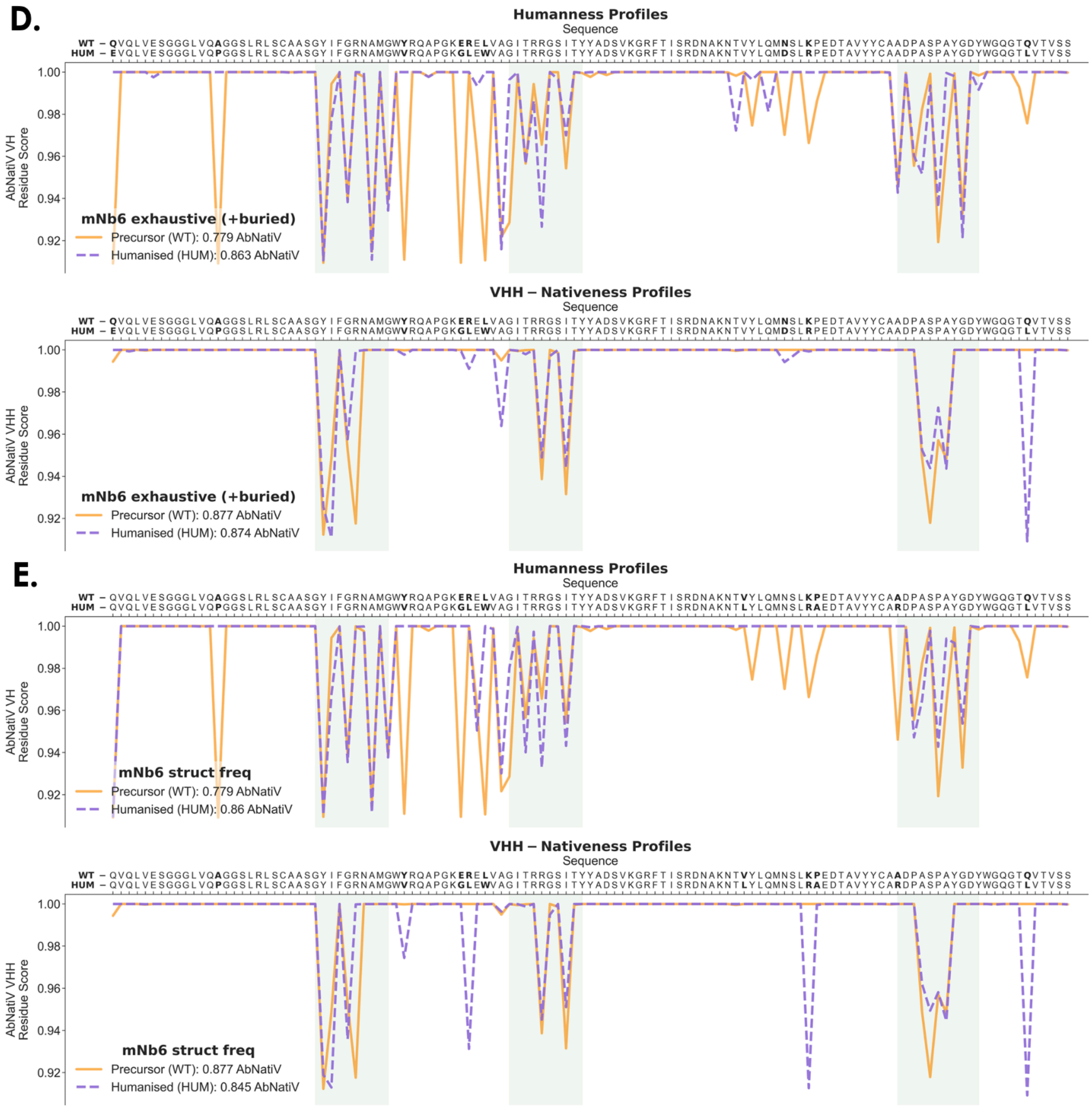


**Supplementary Fig. 16. AbNatiV VH and VHH profiles of the humanised designs of mNb6.** AbNatiV profiles are represented for the WT (in yellow) and humanised counterpart (in purple) of mNb6. (**A**) AbNatiV enhanced sampling. (**B**) AbNatiV exhaustive sampling. (**C**) AbNatiV enhanced sampling allowing buried framework residues. (**D**) AbNatiV exhaustive sampling allowing buried framework residues. (**E**) AbNatiV profiles for the structure and frequency-based humanised variants (this variant was obtained from the Llamanade web server at http://35.208.211.136/, see Methods).

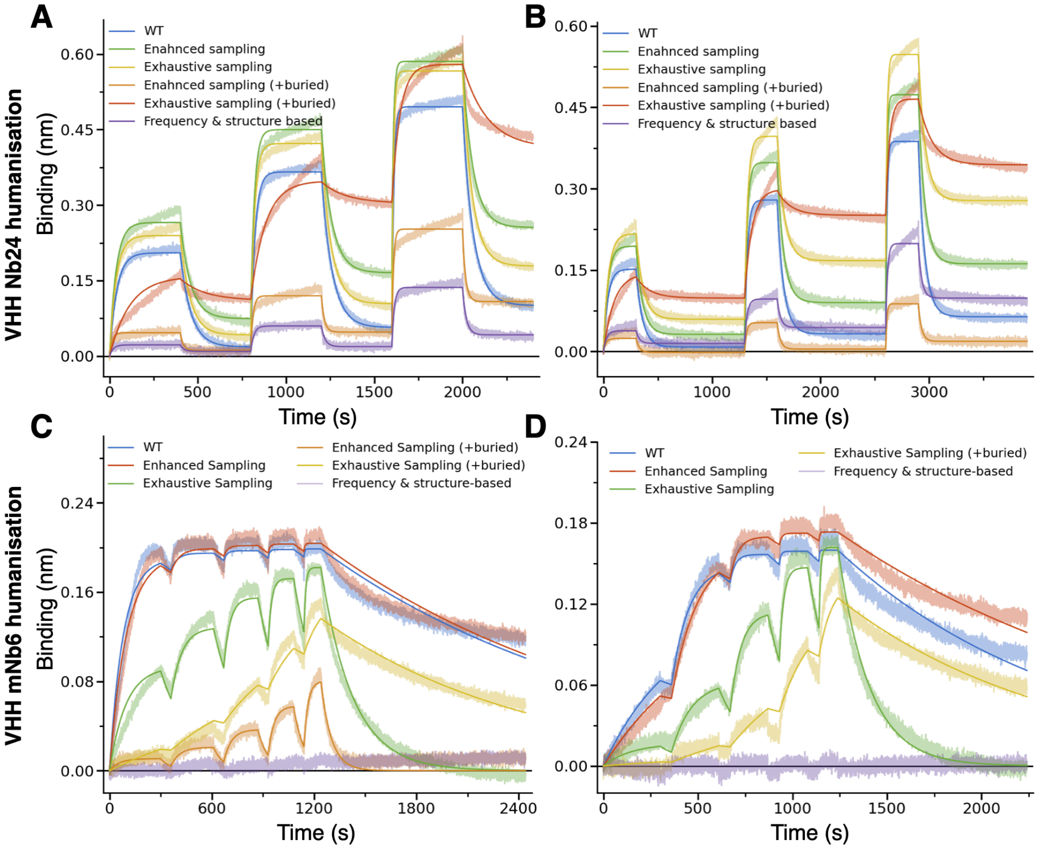


**Supplementary Fig. 17. Additional BLI binding experiments.** BLI binding traces (associations and dissociations phases) obtained with SA sensors loaded with biotinylated $\beta_2$-microglobulin (**A**, **B**) or biotinylated SARS-CoV-2 RBD (**C**, **D**). (**A, B**) Association was monitored in wells containing 44.4, 133.3 and 400 nM (**A**) or 33.3, 100, and 300 nM (**B**) of Nb24 nanobody variants (see legend). Data were fitted globally with a 1:1 partial dissociation binding model using $R_{max}$, on rate, and off rate as global parameters and $Y_{t \rightarrow inf}$ as local parameter. (**C, D**) Association was monitored in wells containing 18.75, 37.5, 75, 150 and 300 nM (**C**) or 3.7, 11.1, 33.3, 100, and 300 nM (**D**) of mNb6 nanobody variants (see legend). Data were fitted globally with a 1:1 binding model using $R_{max}$, on rate, and off rate as global parameters.

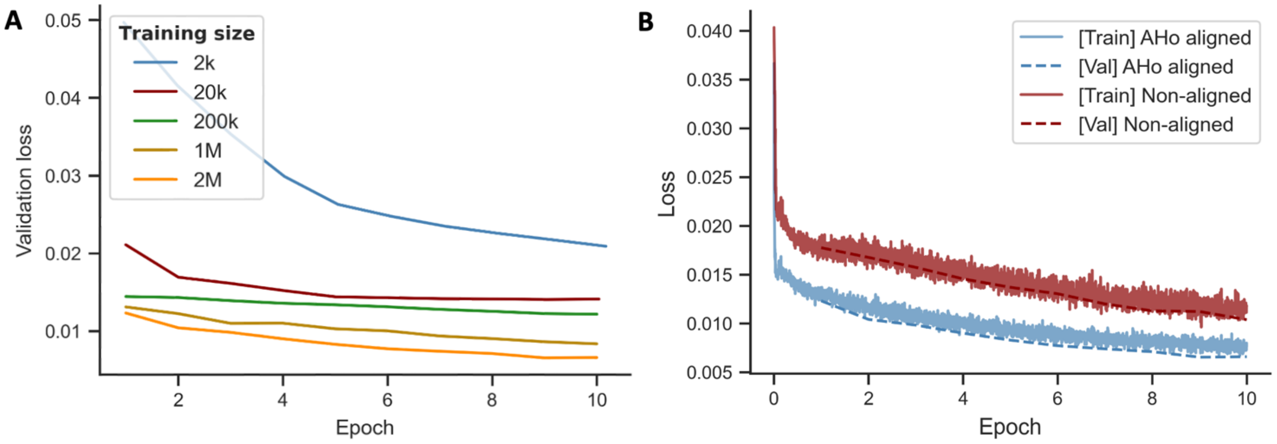


**Supplementary Fig. 18. Influence of training set size and sequence alignment on performance**. (**A**) Validation loss performance as a function of the number of training epochs of the AbNatiV human VH model trained on datasets of different size (see legend). Validation loss performance is always calculated on the same dataset of 50k validation sequences. (**B**) Loss performances on the training (solid line) and validation (dashed line) datasets of the AbNatiV human VH model trained on the same 2 million sequences that were aligned (in blue) or not aligned (in red) as a function of the number of training epochs.

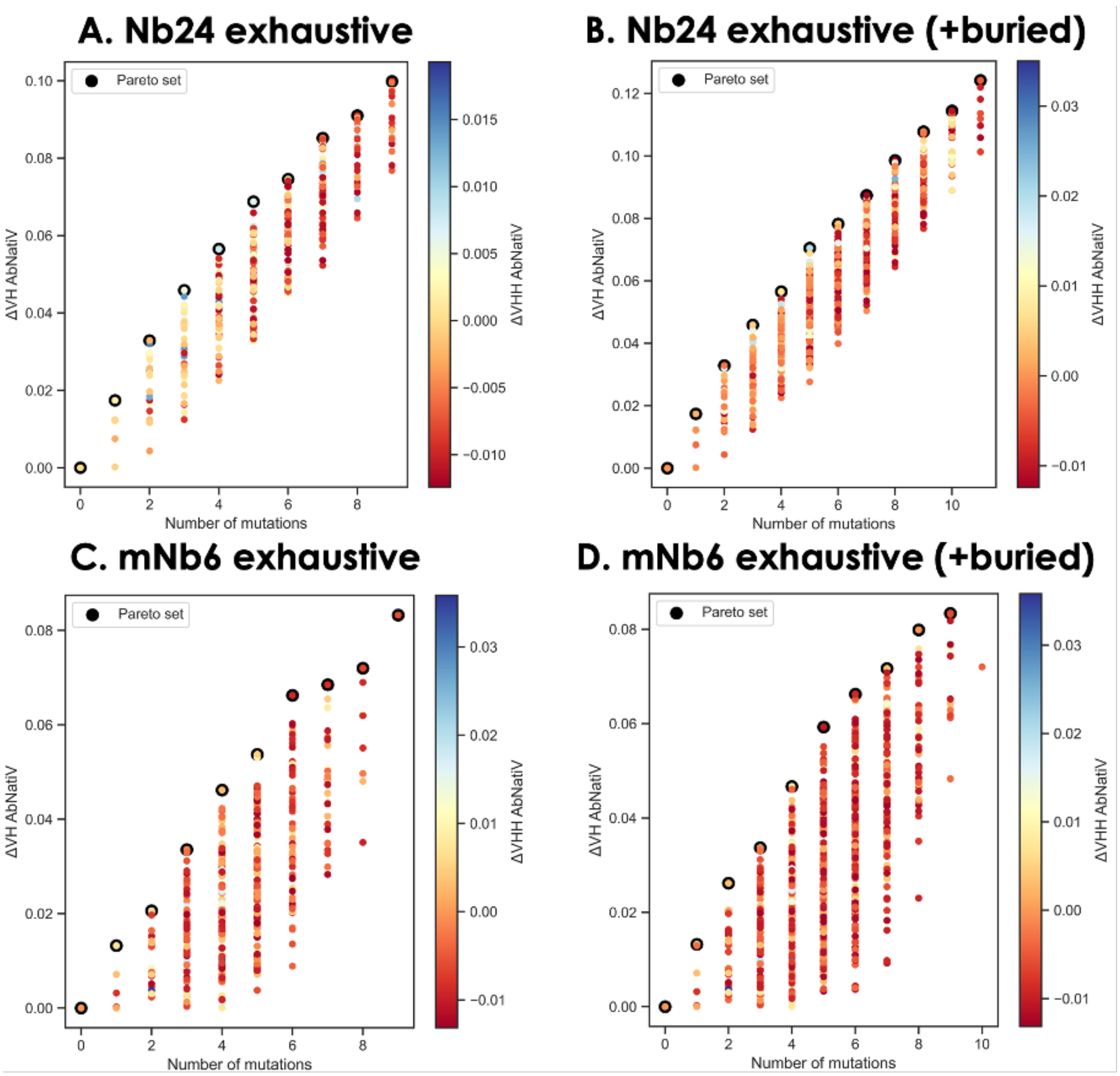


**Supplementary Fig. 19. Pareto front sets returned by the exhaustive sampling of the AbNatiV humanisation of nanobodies pipeline.** The ΔVH AbNatiV humanness score is plotted as a function of the number of mutations. The Pareto front computed maximises the VH-humanness improvement (ΔVH) while minimising the number of mutations over all remaining mutants. Sequences that constitute the Pareto front are circled in black. Each sequence is additionally coloured following its VH-nativeness variation (ΔVHH, see colour-bar). (**A**) Pareto set of the exhaustive sampling for Nb24. (**B**) Pareto set of the exhaustive sampling for Nb24 allowing mutations at buried framework positions. (**C**) Pareto set of the exhaustive sampling for mNb6. (**D**) Pareto set of the exhaustive sampling for mNb6 allowing mutations at buried framework positions.

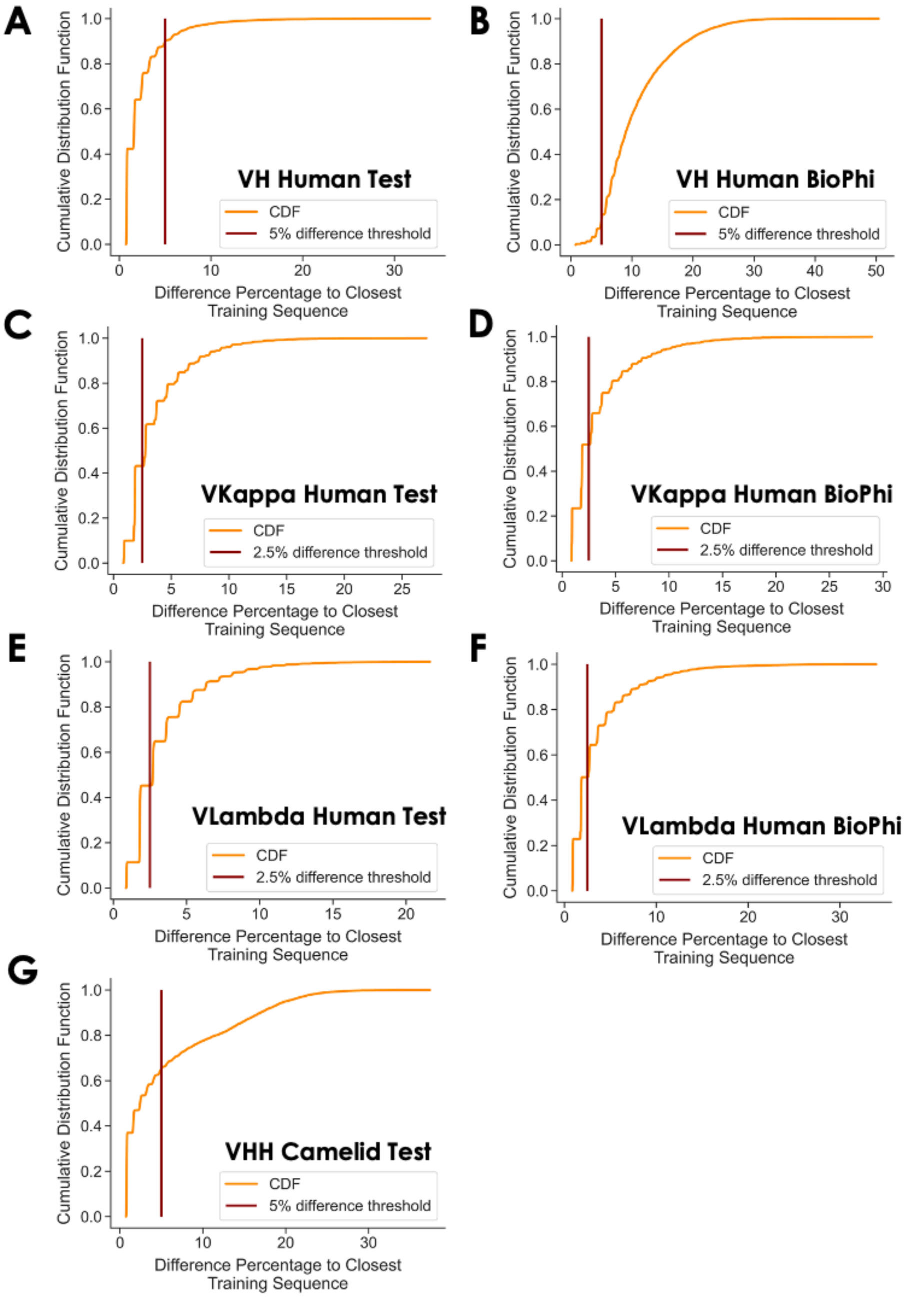


**Supplementary Fig. 20. Minimum percent difference from training sequences.** Each panel reports the cumulative distribution functions (CDFs) (in orange) for the VH Human Test (**A**), VH Human BioPhi (**B**), VKappa Human Test (**C**), VKappa Human BioPhi (**D**), VLambda Human Test (**E**), VLambda Human BioPhi (**F**) and VHH Camelid Test (**G**). The x-axis reports the percent difference to the closest sequence in the respective training dataset. In red is specified the minimum difference percentage threshold used to define, for each AbNatiV model, the corresponding Diverse >5% dataset (or >2.5% for the light chains). For VH, VKappa and VLambda, sequences with a higher difference percentage from both the Test and BioPhi datasets are combined to yield to the Diverse datasets.

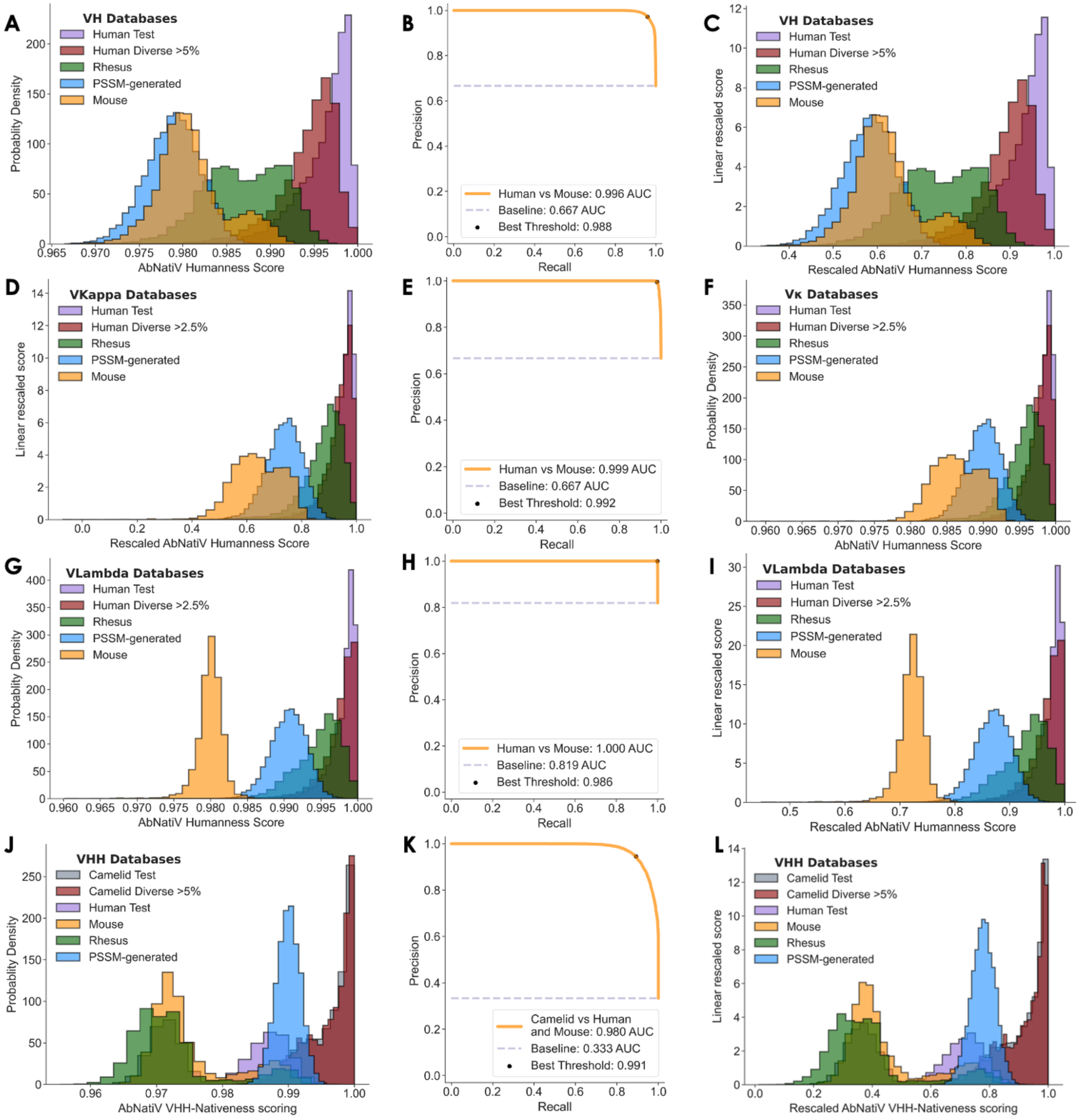


**Supplementary Fig. 21. Linear rescaling of the AbNatiV nativeness score**. (**A, D, G, J**) The AbNatiV nativeness score distributions of various datasets (see legend) after the exp $(-X)$ projection but before the linear rescaling for models trained on human VH (**A**), VKappa (**D**), VLambda (**G**) and camelid VHH (**J**) sequences. (**B, E, H, K**) The optimal thresholds $T_R$ (in black) extracted as the point closest to (1,1) in the PR curves when separating the Human Test sequences from the Mouse sequences for the models trained on human VH (**B**), VKappa (**E**), and VLambda (**H**) sequences, and when separating Camelid Test sequences from the Human Test and Mouse sequences for the model trained on camelid VHH sequences (**K**). (**C, F, I, L**) The AbNatiV nativeness score distributions (respectively from **A, D, G,** and **J**) after linear rescaling $((0.8 - 1) * (X - 1)/(T_R - 1) + 1)$ for each model following their respective $T_R$ (respectively from **C, F, I,** and **L).**

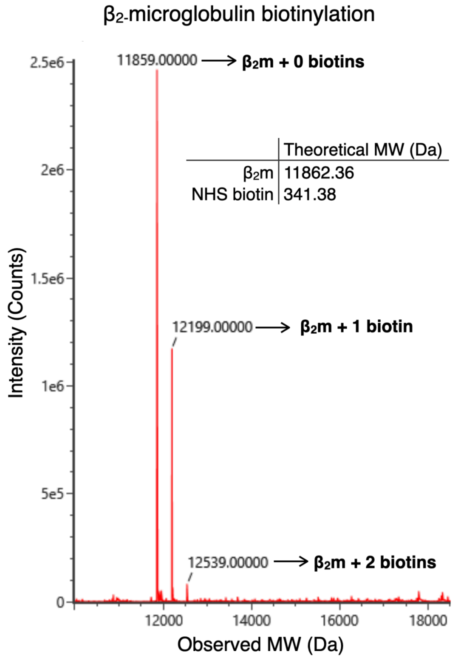


**Supplementary Fig. 22. LC-MS of biotinylated $\beta_2$-microglobulin.** Deconvoluted LC-MS spectrum of biotinylated $\beta_2$-microglobulin after SEC purification. Detected peaks correspond to the products indicated by arrows. The unreacted $\beta_2$-microglobulin ($\beta_2$m+ 0 biotins) will not load onto BLI SA sensors, meaning that most of the protein used in binding assays has exactly one biotin per protein, with a subpopulation with 2 biotins per protein.

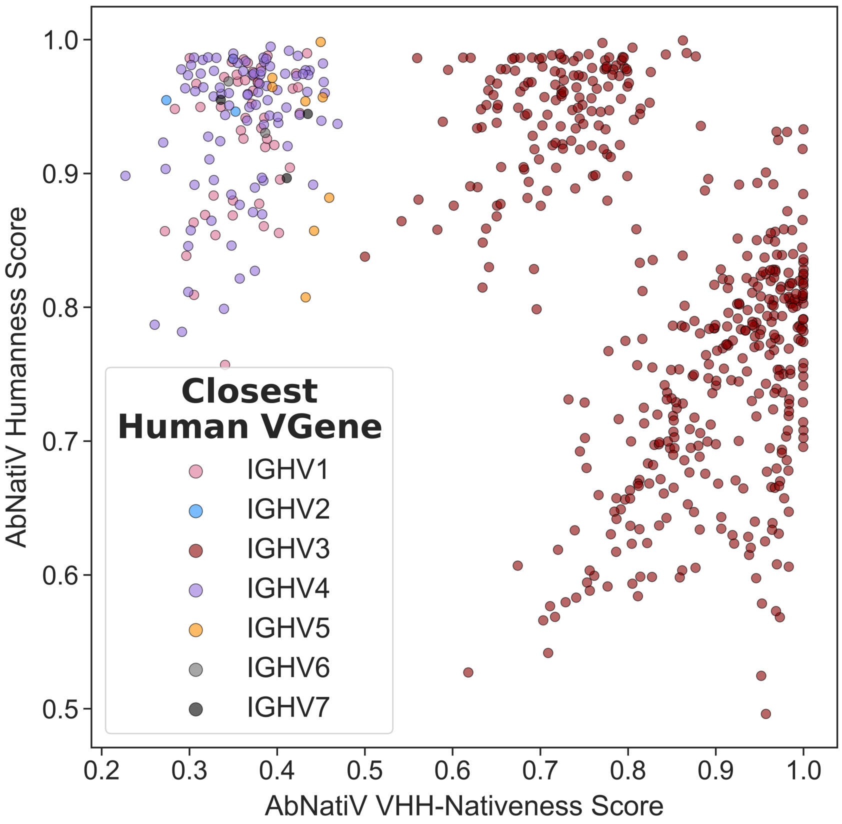


**Supplementary Fig. 23. Combining AbNatiV humanness and VHH-nativeness**. Plot of the AbNatiV humanness and VHH-nativeness scores of 300 sequences from the VH Human Test (in red), and VHH Camelid Test (in blue) datasets from **Extended Data Figure 2** of the main text. Datapoints (sequences) are colored based on their closest human germline V gene as identified by the ANARCI software.
